# Impact of Evolutionary Relatedness on Species Diversification and Tree Shape

**DOI:** 10.1101/2023.11.09.566365

**Authors:** Tianjian Qin, Luis Valente, Rampal Etienne

## Abstract

Slowdowns in lineage accumulation are often observed in phylogenies of extant species. One explanation is the presence of ecological limits to diversity and hence to diversification. Previous research has examined whether and how species richness (SR) impacts diversification rates, but rarely considered the evolutionary relatedness (ER) between species, although ER can affect the degree of interaction between species, which likely sets these limits. To understand the influences of ER on species diversification and the interplay between SR and ER, we present a simple birth-death model in which the speciation rate depends on the ER. We use different metrics of ER that operate at different scales, ranging from branch/lineage-specific to clade-wide scales. We find that the scales at which an effect of ER operates yield distinct patterns in various tree statistics. When ER operates across the whole tree, we observe smaller and more balanced trees, with speciation rates distributed more evenly across the tips than in scenarios with lineage-specific ER effects. Importantly, we find that negative SR dependence of speciation masks the impact of ER on some of the tree statistics. Our model allows diverse evolutionary trajectories for producing imbalanced trees, which are commonly observed in empirical phylogenies but have been challenging to replicate with earlier models.

## Introduction

Phylogenetic trees are important tools to estimate past processes that may explain the current species richness of clades, such as diversification rates. Over the last two decades, the increasing availability of DNA sequence data and of tools to reconstruct phylogenetic trees from these data has led to the development of birthdeath models that use molecular phylogenies as a source of information to study diversification dynamics (Nee et al., 1992; Purvis, 2008; Quental and Marshall, 2010; Etienne et al., 2012). Lineage-through-time (LTT) plots, semi-logarithmic plots that track the number of lineages that have descendants at the present through time, are a powerful way of summarizing diversification dynamics. If per-lineage rates of speciation and extinction have been constant through time, the accumulation of lineages increases through time exponentially, with an even stronger increase close to the present, a phenomenon called the ‘pull-of-the-present’ (Nee et al., 1994; Kubo and Iwasa, 1995). However, a large number of empirical phylogenies display a different pattern: they often show recent deceleration (Purvis, 2008; Phillimore and Price, 2008; Moen and Morlon, 2014; Aguilée et al., 2018), which contrasts with findings from the fossil record (Etienne et al., 2012; Louca and Pennell, 2021).

Several hypotheses have been put forward to explain the observed slowdowns in phylogenies of extant species (Moen and Morlon, 2014), such as time-dependent speciation rate (Moen and Morlon, 2014), protracted speciation (Etienne and Rosindell, 2012) and negative diversity-dependent diversification (Valentine, 1973; Sepkoski, 1978; Etienne et al., 2012). In negative diversity-dependent diversification models, the speciation rate declines with increasing species diversity (SR).

Current models of diversity-dependent diversification assume that speciation rate depends on global SR, regardless of evolutionary relationships of the species. The general underlying idea is that species diversification results in the occupation of available niche spaces, leaving limited opportunities for subsequent species to use those niches (Figure 1), because evolving clades compete for finite ecological resources (Wiens, 2011). In these models, SR acts as a proxy for more mechanistic factors such as functional traits, niches and ecological interactions, which ultimately may influence ecological limits (Srivastava et al., 2012; Kondratyeva et al., 2019). While past ecological interactions between species are difficult to infer, useful proxies are evolutionary relatedness (ER) metrics, which quantify the phylogenetic distance between taxa. ER can represent the ecological roles of the species well (Helmus et al., 2007; Pigot et al., 2016), as closely related species are more likely to share trait states and ecological functions than distantly related species (Srivastava et al., 2012), and evolutionary distance may be related to fitness and niche differences (Cadotte et al., 2012), which in turn can affect speciation and extinction. Here, we may learn from insights from phylogenetic community ecology (Webb et al., 2002; Mayfield and Levine, 2010; HilleRisLambers et al., 2012) and invasion biology. For example, alien species that are evolutionarily distant from those in the local community may have a greater chance of establishing (Zheng et al., 2018), so using ER as a proxy to quantify open niche spaces may explain the success of plant invaders (Henn et al., 2019; Qin et al., 2020).

**Figure 1.**
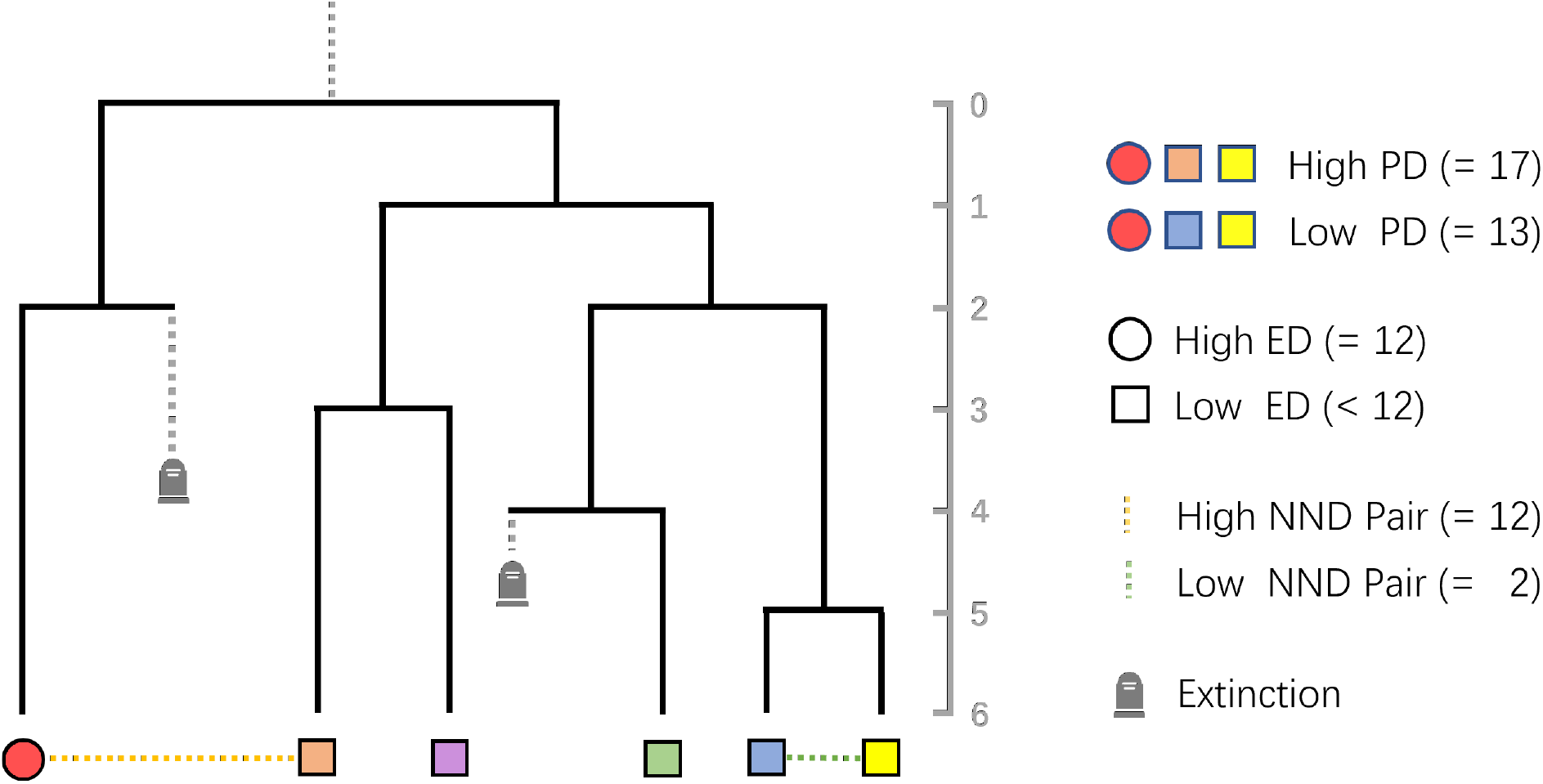
Illustration of how evolutionary relatedness (ER) is computed within clades and species in a phylogeny, as measured by three metrics: PD (phylogenetic diversity), ED (evolutionary distinctiveness), and NND (nearest neighbor distance). The tree on the left represents a phylogeny with extant and extinct (marked by tombstone icons) species. The numbered axis in the middle marks the branch lengths in evolutionary time (million-years). The colored circle and squares at the tips of the phylogeny represent corresponding species. On the right of the plot, the species denoted by a grey circle, an orange square and a yellow square form a combination of high PD (17), while the species denoted by the grey circle, the blue square and the yellow square form a combination of low PD (13). The species with the highest ED value (12) is represented by a circle, and those with lower ED values (less than 12) are represented by a square. The yellow and green dashed-lines between species illustrate two pairs of species, one with a low NND value (2) and one with a high NND value (12).

The potentially powerful phylogenetic proxies for ecological interactions, however, have been the subject of intense debate because the underlying assumptions are often not robustly supported (Gerhold et al., 2015; Pigot et al., 2016). Indeed, there is mixed evidence regarding the effectiveness of these proxies. On the one hand, Tucker et al. (2018) concluded that phylogenetic diversity is a useful proxy for functional diversity, because phylogenetic diversity correlates more strongly with functional diversity when trait dimension increases (multidimensional trait space), although that correlation is weakened when trait evolution models become increasingly complex. On the other hand, Mazel et al. (2018) found that in many cases phylogenetic diversity only poorly captures functional diversity, even less so than random selection. Furthermore, Venail et al. (2015) found that trait and functional variation among species is largely explained by SR but not phylogenetic relatedness (a concept similar to ER). Recently, Pie et al. (2023) found that sympatry with closely-related species does not lead to decreasing speciation rates in a variety of vertebrate clades, implying that the underlying mechanisms of diversity-dependent diversification remain unconfirmed.

While these concerns indicate that support for a role of ER on diversification is not clear, this may be because we do not know what signal ER is expected to leave in phylogenetic trees and extant communities (Rabosky, 2009b; Ricklefs, 2010; Wiens, 2011). Most studies have only considered diversity-dependence within clades of phylogenetically closely related species (Rabosky and Glor, 2010; Etienne et al., 2012; Foote et al., 2018), whereas only a few have considered diversity-dependence between clades of phylogenetically disparate species (Pires et al., 2017). A recent study (Etienne et al., 2023) using empirical data on island frogs found that a model with diversity-dependence between closely related species was preferred over one with diversity-dependence also occurring with more distantly related species, indicating that interactions of species with close relatives, but not with distant relatives, negatively affect colonization and diversification. This suggests an important role of ER in species diversification and the need for distinguishing between different levels of ER and scales at which it may have an effect on diversification. However, none of the above-mentioned studies has investigated whether ER directly impacts macro-evolutionary dynamics (Rillo and Etienne, 2022).

Assuming ER affects diversification rates, what signatures does this effect leave in phylogenetic trees of communities? What emergent phylogenetic patterns are expected if species facilitate or compete more strongly with their close relatives? Can phylogenetic limits imposed by ER be differentiated from those imposed by SR? And how do ER effects operating at the whole clade versus more lineage-specific scales affect phylogenies? To address these questions, here we present a new phylogenetic birth-death simulation model that incorporates both SR and ER. The model allows for positive, neutral, or negative effects of both SR and ER on diversification rates.

We measure ER using three different mechanisms of how ER can affect diversification (see Methods) each considering a different scale of the effect of ER. We use the new model to simulate communities under a stochastic branching process where speciation rate can vary according to SR and ER, and analyze whether ER leaves a signature on the diversification dynamics by looking at various tree summary statistics. We aim to provide the first expectations for the effects of these processes on phylogenetic trees by developing a simple simulation model where the effects of SR and ER can be studied independently or in combination. While inferring parameters from empirical datasets is beyond the scope of this study, our model presents a tool that can be used in future simulation-based parameter estimation methods, for example for generating training datasets for neural network models. Our R package “evesim” on GitHub contains functions to generate simulated phylogenies using our model (Hildenbrandt and Qin, 2024).

## Methods

### Model

We employed a phylogenetic stochastic model that simulates the processes of species birth and death over time (Nee et al., 1994). The model is a mechanistic process-based model that is parameterized by speciation and extinction rates along with the effect sizes of SR and ER. It allows us to explore different evolutionary trajectories from phylogenetic trees given specific parameter settings at the start of the simulation. We assumed that all species on the same tree belong to the same clade, thus our study focuses on species diversification patterns within a clade, and the phylogenies simulated include all the extant species in the clade. We investigated different types of effects of ER on species diversification by considering three measures of ER: phylogenetic diversity (PD, community-level metric), evolutionary distinctiveness (ED, per-lineage metric), and nearest neighbor distance (NND, per-lineage metric).

PD measures the amount of evolutionary history represented by a group of species and is commonly used to determine how species occupy different niches (Tucker et al., 2018). We calculated PD using Faith’s index (Faith, 1992), which represents the total branch length of a phylogenetic tree reconstructed from all species in a clade. ED quantifies the uniqueness of each of the species relative to other species and is a valuable tool in conservation efforts (Cadotte and Davies, 2010). Per species, ED is defined as the sum of the pairwise distances between focal species and all other species, divided by the number of species minus one. NND also quantifies the uniqueness of a species, but only on a very local phylogenetic scale, as it is defined as the phylogenetic distance (branching length of the path) between a focal species and its nearest neighbor. NND measures the degree to which each of the species is locally clustered within specific clades, and is less sensitive to higher-level phylogenetic structures than ED (Webb, 2000). Unlike PD, which is a clade-level metric and assigns a single value for all species in the clade, ED and NND are lineage-specific metrics, with each lineage assigned its own value.

There are many other choices of ER metrics, e.g. the “Fair Proportion” index (Redding et al., 2008) and the “Evo-Heritage” metric (Rosindell et al., 2024). However, we aimed to use metrics that are computationally efficient.

*λ*_*i*_, the speciation rate of a specific lineage *i*, is given by

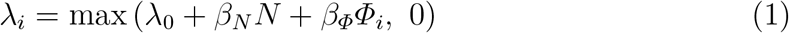

where *λ*_0_ is the intrinsic speciation rate of all the lineages, *N* is the species richness (across all lineages), *Φ*_*i*_ represents ER (either PD, ED or NND) of the lineage *i, β*_*N*_ is a coefficient to adjust the effect size of species richness on the speciation rate and *β*_*Φ*_ is a coefficient to adjust the effect size of ER on the speciation rate. *β*_*N*_ and *β*_*Φ*_ can be positive, zero, or negative. We note that by negative ER we mean negative *β*_*Φ*_ and thus that as species become evolutionarily less related, speciation rate decreases. Furthermore, if we set both *β*_*N*_ and *β*_*Φ*_ to zero, the model reduces to a standard birth-death model.

When using PD as an ER metric, we assume that *Φ*_*i*_ is the same for every lineage *i*, so the speciation rates of all lineages are equal. Unlike PD, both ED and NND are calculated separately for every lineage.

To account for the strong correlation between phylogenetic diversity and time when using PD, we included an offset method in our model to compensate for the inflation of branch lengths in the phylogenetic tree. The method subtracts the tree age *t* from *Φ*_*t*_, which is the phylogenetic diversity at time *t*. The adjusted 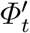 is then given by

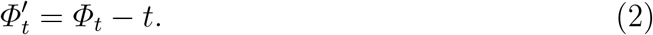

We assumed that the extinction rate *µ* is fixed to the intrinsic rate of extinction which is constant through time and across lineages:

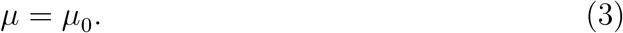

### Simulations

We ran a series of simulations of the model under different scenarios in order to investigate the effect and signature of ER on various phylogenetic summary statistics. We started the simulations with two ancestral lineages, setting the values of ER of both lineages to zero. We used the Gillespie algorithm (Gillespie, 1976), in which the waiting time between two evolutionary events is sampled from an exponential distribution with a mean equal to the inverse of the sum of the rates of all possible events. The probability of each event is proportional to its own rate relative to the sum of the rates of all possible events.

Two types of events can occur: speciation and extinction. When a speciation event happens, a lineage at the tip of the tree bifurcates into two lineages. When an extinction event happens, a species is marked as extinct. The simulation lasts for a predetermined time, which equals the crown age of the final phylogeny. A successful simulation is conditional on survival of both crown lineages; the simulation will start over if one of the crown lineages goes extinct entirely (see Figure 2 for the illustration of the simulation). Our GitHub repository “eve” (Qin, 2023) contains the codebase for the current study.

**Figure 2.**
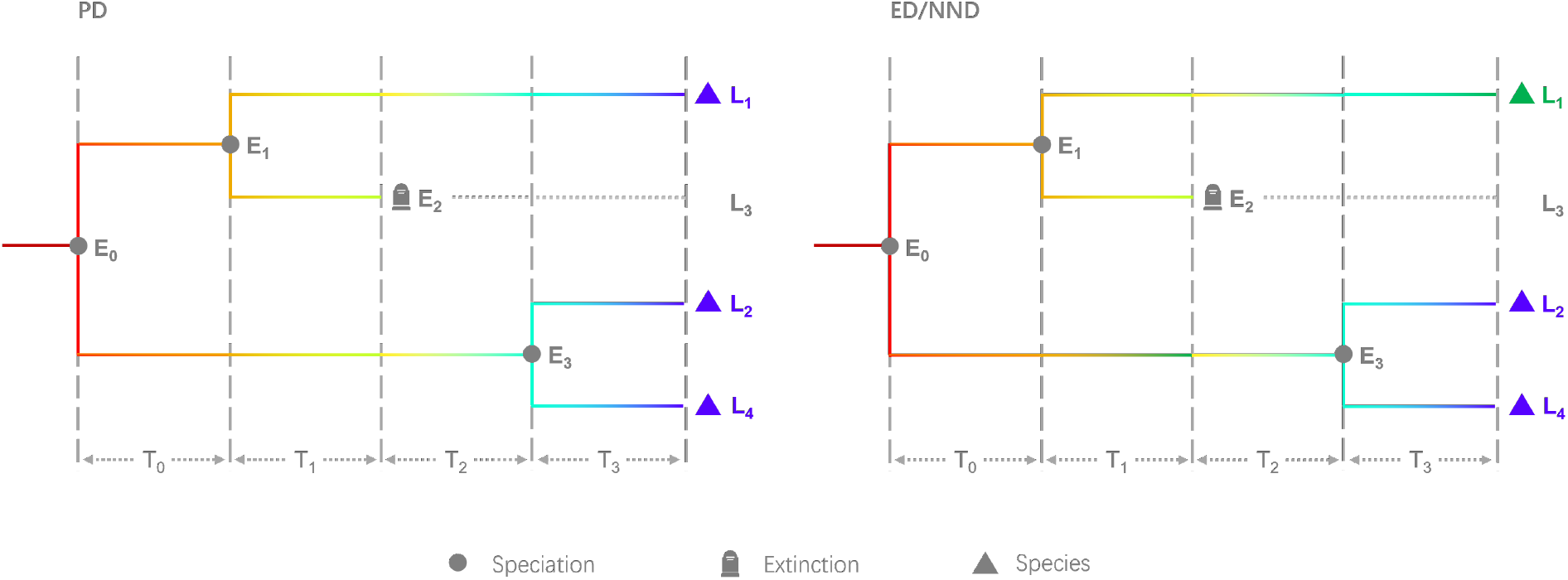
Illustration representing a stochastic simulation under the assumptions of positive speciation and extinction rates alongside a negative coefficient of evolutionary relatedness, exemplified through a scenario where phylogenetic diversity (PD, left panel) or evolutionary distinctiveness (ED, right panel) or nearest neighbor distance (NND, also right panel) acts as clade-wide (PD) or lineage-specific (ED and NND) constraint. Although ED and NND scenarios are illustrated in the same panel, the underlying processes are different. Two event types are depicted: speciation (solid grey circles) and extinction (tombstone symbols). During a speciation event, a lineage at a tree tip bifurcates into two lineages (e.g., *E*_0_, *E*_1_ and *E*_3_). An extinction event marks a species as extinct (e.g., *E*_2_). In the left panel, the branch color transition from red to blue signifies the variation in the speciation rate of lineages along PD. In the right panel, the branch color transition from red to blue or green signifies the variation of per-lineage speciation rates within the phylogeny due to lineage specific (ED or NND) phylogenetic diversity-dependence. The simulation unfolds as follows: it initiates from *E*_0_ with two ancestral lineages *L*_1_ and *L*_2_, setting the initial PD value to 0. Speciation rates for all extant lineages (initially *L*_1_ and *L*_2_) are derived from PD. The first time interval, *T*_0_, extends the branch lengths of *L*_1_ and *L*_2_. Prior to sampling event *E*_1_, PD is recalculated based on extant lineages, and speciation rates are updated. Event *E*_1_ illustrates *L*1 bifurcating into *L*_1_ and *L*_3_. The subsequent time interval *T*_1_ further extends the branch lengths of *L*_1_, *L*_2_ and *L*_3_. Before sampling each event, PD and speciation rates are updated. Event *E*_2_ marks the extinction of *L*_3_, halting its branch growth while *L*_1_ and *L*_2_ continue to extend through time interval *T*_2_. Event *E*_3_ illustrates the speciation of *L*_2_ into *L*_2_ and *L*_4_. The final time interval, *T*_3_, stops the simulation because the cumulative time (*T*_0_ + *T*_1_ + *T*_2_ + *T*_3_) surpasses a pre-determined time threshold, *T*. *T*_3_ is then set to *T −* (*T*_0_ + *T*_1_ + *T*_2_). The branch lengths of *L*_1_, *L*_2_, and *L*_4_ extend by *T*_3_, marking the simulation endpoint.

We simulated phylogenies using a variety of parameter combinations (see Table 1). We assumed a crown age of 6 time units, which can be interpreted as 6 million years. All combinations of parameters in Table 1 were used, to a total of 135 combinations, and each was repeated for the three different ER scenarios: PD, ED and NND. Thus, we had a total of 405 parameter sets. For each set we simulated 100 phylogenetic trees using the Peregrine high performance computing cluster of the University of Groningen. In our preliminary tests, we increased the number of replicates from 100 to 300 and then 1000 for the fastest parameter settings, but we did not observe noticeable trend changes with the increased number of replicates. Due to hardware limitations, we retained the number of 100 for consistency across all combinations. The parameter sets were chosen such that the simulation can be finished within the time and resource limits of the cluster. Moreover, the effects of *β*_*Φ*_ on the diversification process are inherently non-symmetrical because positive *β*_*Φ*_ results in a positive feedback, which has a much greater influence on the final size of the phylogenies. For this reason, we kept the positive *β*_*Φ*_ values relatively small. Typically, when setting *λ*_0_ = 0.6, *µ*_0_ = 0, *β*_*N*_ = 0, and *β*_*Φ*_ *< −*0.1, the output phylogeny will be very small (often less than five lineages) and hence not very meaningful. If *β*_*Φ*_ *>* 0.002, then the resulting phylogenies will be very large (often more than 500 lineages), which creates computational problems (many simulation steps with compututionally demanding calculation of the phylogenetic metrics).

**Table 1:** Parameters used in the simulations.

| Parameter | Value |
| --- | --- |
| Intrinsic speciation rate | 0.4, 0.5, 0.6 |
| Intrinsic extinction rate | 0, 0.1, 0.2 |
| Crown age | 6 |
| Coefficient species richness $N$ ($\beta_N$) | -0.04, -0.02, 0 |
| Coefficient evolutionary relatedness ($\beta_\Phi$) | -0.04, -0.02, 0, 0.001, 0.002 |
| Evolutionary relatedness metric | PD, ED, NND |

We deliberately chose a relatively simple model to assess the effect of ER on diversification, rather than including a variety of realistic ecological interactions linked with ER. It allows us to explore parameter space better, and it may facilitate future parameter estimation from phylogenies. Most importantly, it allows gaining broad generalizable insights based on the fundamental processes we are interested in (i.e., ER and SR effects on diversification).

### Data analysis

Raw output from the simulations was processed using the “eve” package into data compatible with statistics functions in other R packages. For each parameter set, the “treestats” (Janzen and Etienne, 2024) and “eve” packages were used to calculate summary statistics for all extant trees, excluding those with only two extant lineages. The statistics chosen were the following: the J One balance index (Lemant et al., 2022), the Gamma statistic (Pybus and Harvey, 2000), mean branch length, mean pairwise distance (MPD) (Webb et al., 2002) and the Rogers J index of imbalance (Rogers hereafter) (Rogers, 1996), see glossary for their definitions.

The effects of SR and ER were measured by the values of *β*_*N*_ and *β*_*Φ*_, respectively. The effects of the scale at which the effect operates, from whole-clade, to species-specific were determined by the ER mechanism (one of the three ER measures: PD (whole-clade), ED (intermediate) and NND (species-specific).

### Speciation rate evenness

In the ED and NND scenarios, speciation rates are expected to vary between tree tips. We measured how these rates are distributed in phylogenies by adopting a concept similar to measuring species evenness in a community, but instead quantifying the evenness of speciation rates across lineages weighted by their phylogenetic distances. Each phylogeny of *n* lineages was given by a correlation matrix ***C*** with each lineage *i*’s speciation rate represented by *λ*_*i*_ (*i* = 1, 2, 3, …, *n*). The phylogenetic evenness index *E* is then defined as

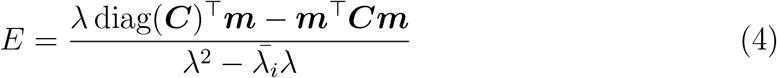

which was originally proposed by Helmus et al. (2007). In the equation, *λ* denotes sum(*λ*_1_, *λ*_2_, *λ*_3_, …, *λ*_*n*_) and 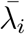 denotes mean(*λ*_1_, *λ*_2_, *λ*_3_, …, *λ*_*n*_), diag(***C***) denotes a column vector comprising the diagonal elements of the correlation matrix ***C*** (see below), ***m*** denotes an *n ×* 1 column vector containing values of *λ*_*i*_. The value of *E* ranges between 0 and 1. The maximum value (i.e., 1) of *E* only occurs when speciation rates among lineages are equal and the tree is completely balanced (star-like). Values of *E* less than 1 represent increasing unevenness of speciation rates among the lineages. *E* is sensitive to tree topology such that trees with uniform speciation rates can have different values of evenness.

The correlation matrix ***C*** is the standardized pairwise distance matrix. Because we only consider ultrametric (no extinct lineages) and binary (fully resolved) phylogenies, ***C*** can be computed as:

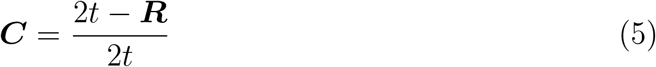

where ***R*** is the pairwise distance matrix of all the lineages of the phylogeny of *n* lineages, with *r*_*ij*_ (*i* = 1, 2, 3, …, *n*; *j* = 1, 2, 3, …, *n*) represents the pairwise phylogenetic distance between lineage *i* and *j*:

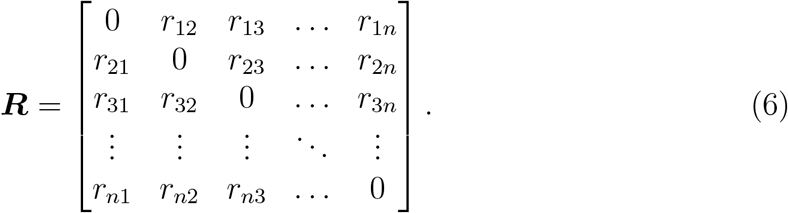

As shown in Equation 5 and Equation 6, all elements of diag(***C***) are equal to one, Equation 4 can thus be simplified as:

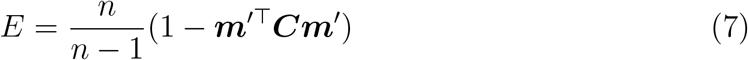

where ***m***^*′*^ = ***m****/λ*.

Note that this simplification is only possible for ultrametric trees with *n ≥* 2, a formal proof of Equation 5 and the derivation of Equation 7 are provided in Appendix F and Appendix G.

In the case of simulated data, the models and parameters are already known to differ. Testing for statistical significance between treatments is thus meaningless; the distributional changes sufficiently demonstrate the influence of the parameters. Therefore, we focused on how summary statistics vary with the strength of ER and SR to explain the model’s power and effects.

### Data visualization

For visualization purposes, we selected one representative tree for each tree set of 100 trees representing a single parameter set. This tree was chosen among the complete trees based on its index vector, which has the smallest mean Mahalanobis distance (De Maesschalck et al., 2000) to the other trees in the same tree set (as defined in Equation 8 below). In order to calculate the Mahalanobis distance for each tree, we first defined the index vector for the *i*-th tree in a set of trees resulted from *k*-th parameter set, where *k* denotes one of the parameter sets in the 405 combinations in our simulation, as

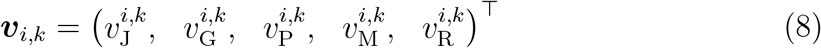

where each element of ***v***_*i,k*_ represents a summary statistic of the *i*-th tree in the *k*-th set. 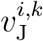 denotes the J One balance index, 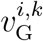 denotes the Gamma statistic, denotes PD, 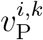 denotes the mean pairwise distance, 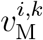 denotes the Rogers balance index and *⊤* denotes the transpose of vector, see glossary for descriptions and citations for these measures. These five statistics were selected based on a clustering dendrogram of various summary statistics, where they were found to be less correlated and more evenly spaced than other statistics (see Appendix B).

The Mahalanobis distance of ***v***_*i,k*_ is given as

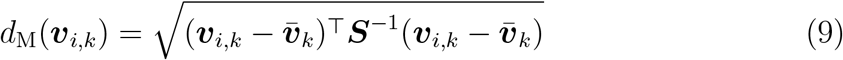

where ***v***_*i,k*_ is the index vector for the *i*-th tree of the *k*-th set. 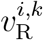 comprises five mean values of each element respectively in the index vectors in the *k*-th set, *⊤* denotes the vector transpose, ***S*** is the covariance matrix of these vectors and ***S***^*−*1^ denotes the inverse of ***S***.

For each representative tree (i.e. the tree with the smallest Mahalanobis distance to the other trees), we mapped the historical speciation rates onto the tree using the “eve” package. The variation of speciation rates along tree lineages is represented by a continuous color scale. The actual rate values corresponding to colors were calculated as mean speciation rates (mean value of speciation rates at the beginning and at the end) of the time frames between evolutionary events in a simulation.

LTT plots were generated using the “eve” package for each tree set by summarizing all the trees in the set and then displaying the mean LTT curve along with shading representing the confidence interval of the set. The LTT plots were based on extant species only.

## Results

### Simulation performance

All simulation jobs were completed within seven hours on the computing cluster, with most of the parameter sets finishing within one hour.

When extinction rates are 0 and no species richness effect is present, a deceleration in lineage accumulation is observed in the PD scenario, but only when *β*_*Φ*_ is negative, that is, higher ER in the communities results in lower speciation rate on all lineages (see Figure 3 and the animations in Appendix D).

**Figure 3.**
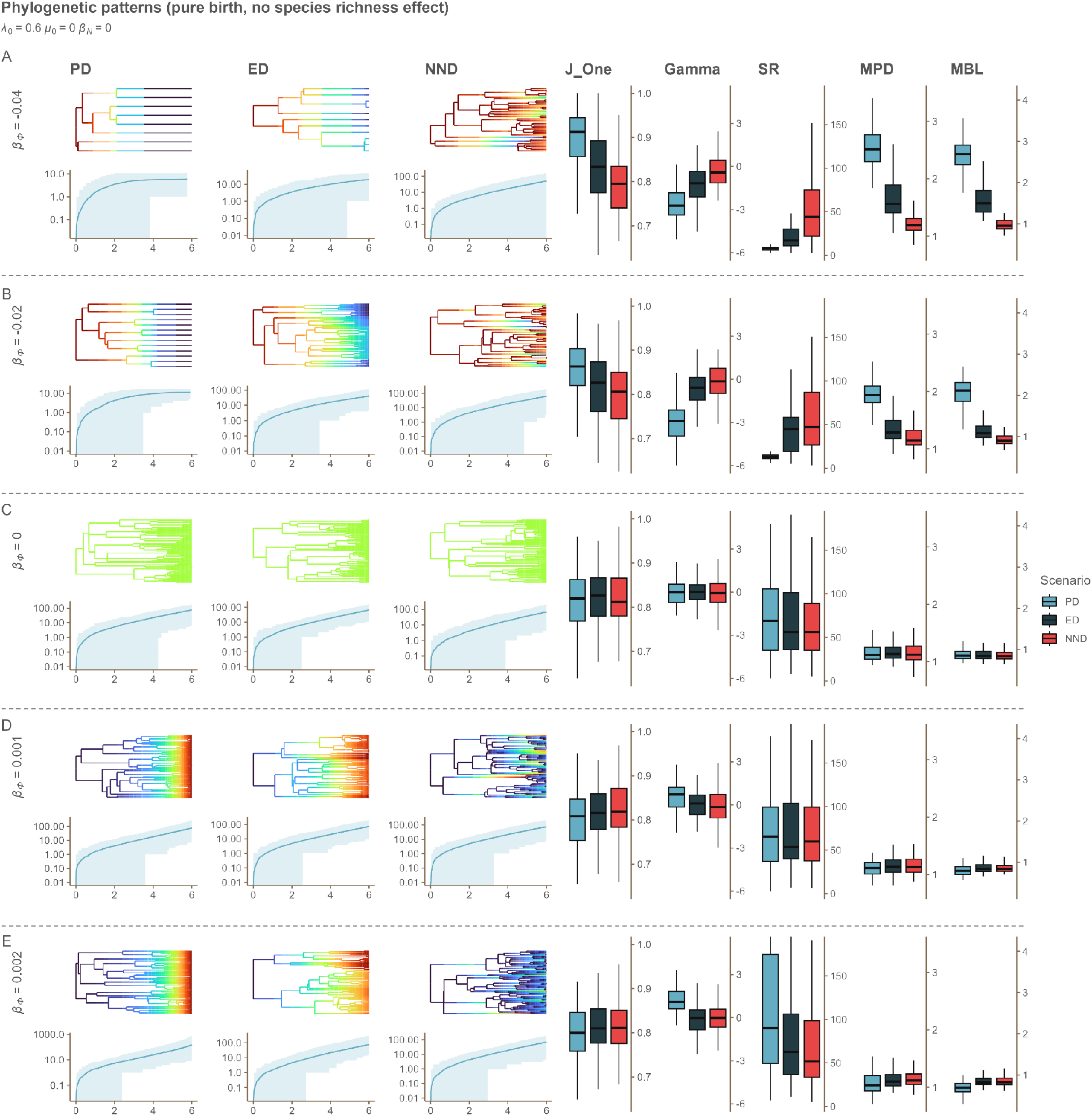
Overview of results of simulations of a pure birth process (no extinction, see Appendix D for other speciation and extinction rate combinations), with dependence of speciation on evolutionary relatedness. For all parameter sets shown in the figure, the intrinsic speciation rate at the start of the simulation (*λ*_0_) is 0.6, the intrinsic extinction rate (*μ*_0_) is 0, and the coefficient of the species richness effect (*β*_N_) is 0. The results are grouped by rows according to varying levels of *β*_Φ_ (the coefficient of the evolutionary relatedness effect). On the left, representative simulated phylogenies and associated lineage-through time (LTT) plots are shown for dependence of speciation rates on three metrics of ER (left - phylogenetic diversity (PD), middle – evolutionary distinctiveness (ED), and right - nearest neighbor distance (NND) scenario). The colors mapped onto the trees represent the values of the speciation rate for each of the lineages during simulation time. The rates increase from blue to red. The trees coloured in green are cases where the speciation rate remains unchanged throughout the simulation. Blue line in the LTT plots represents lineage accumulation through time and the shaded area is the 95% confidence interval. On the right of each row, boxplots for the corresponding summary statistics are shown: J One - J One balance index; Gamma - Gamma statistic; SR - total number of extant lineages; MPD - mean pairwise distance; MBL - the mean branch length.

### Effects of evolutionary relatedness on phylogenetic patterns

We observed a hierarchical structuring in tree topology patterns, from PD to ED to NND. Under the PD scenario, trees exhibit a higher degree of balance (interpreted from the J One balance index) than in the ED scenario, which in turn exhibits more balanced trees than the NND scenario. This pattern is more prominent when *β*_*Φ*_ decreases from zero towards more negative values, that is, when ER reduces the speciation rates even more. As *β*_*Φ*_ increases from negative to positive values, that is, the ER effect on the speciation rates transitions from a reduction to augmentation, the differences in tree balance between PD, ED and NND scenarios decrease, and the trees exhibit generally lower degree of balance (see the animations in Appendix A ending with J One).

As *β*_*Φ*_ increases from -0.04 to 0 (ER effect becomes neutral), a general decrease in the mean pairwise distances of the trees is observed. However, no substantial change occurs when *β*_*Φ*_ shifts from 0 to positive values (i.e., a beneficial effect of ER on speciation). When *β*_*Φ*_ is negative, the trees under the PD scenario exhibit larger mean pairwise distances than those in the ED scenario and the trees under the ED scenario exhibit larger mean pairwise distances than those in the NND scenario. As *β*_*Φ*_ increases from -0.04 to 0, the disparities in mean pairwise distances among PD, ED, and NND scenarios decrease (see the animations in Appendix A ending with MPD and MBL). The patterns of mean branch lengths across *β*_*Φ*_ settings are similar to those of mean pairwise distances.

The distribution of internal nodes, or the concentration of speciation events, were interpreted using the Gamma statistic. When *β*_*Φ*_ is negative, the speciation events in the PD scenario are located more closely to the root than in the ED scenario, and the events in the ED scenario are closer to the root than those in the NND scenario. These differences are reduced as *β*_*Φ*_ increases from negative values towards zero. This pattern changes as *β*_*Φ*_ shifts from zero towards positive values, particularly when *β*_*N*_ is 0. As *β*_*Φ*_ increases from -0.04 to 0.002, the distributions of speciation events in all three scenarios change from being closer to the root to being more evenly distributed throughout the temporal range of the phylogeny (see the animations in Appendix A ending with Gamma).

The trees become notably larger (more species) as *β*_*Φ*_ increases from -0.04 to 0. This increase is minimized when *β*_*Φ*_ increases from 0 to 0.002. When *β*_*Φ*_ is negative, the tree sizes are generally largest in the NND scenario, second largest in the ED scenario and smallest in the PD scenario. This pattern does not persist when *β*_*Φ*_ is zero or positive. The pattern is reversed when *β*_*Φ*_ is positive, in which case the PD scenario generally has the largest tree sizes and the NND scenario the smallest (see the animations in Appendix A ending with SR).

Of all model parameters, *β*_*Φ*_ has the strongest effect on the evenness of the speciation rates across lineages. In all parameter combinations, the disparities of speciation rate evenness between PD, ED and NND scenarios decline with increasing *β*_*Φ*_ (see Figure 4B and the animations in Appendix A ending with ERE).

**Figure 4.**
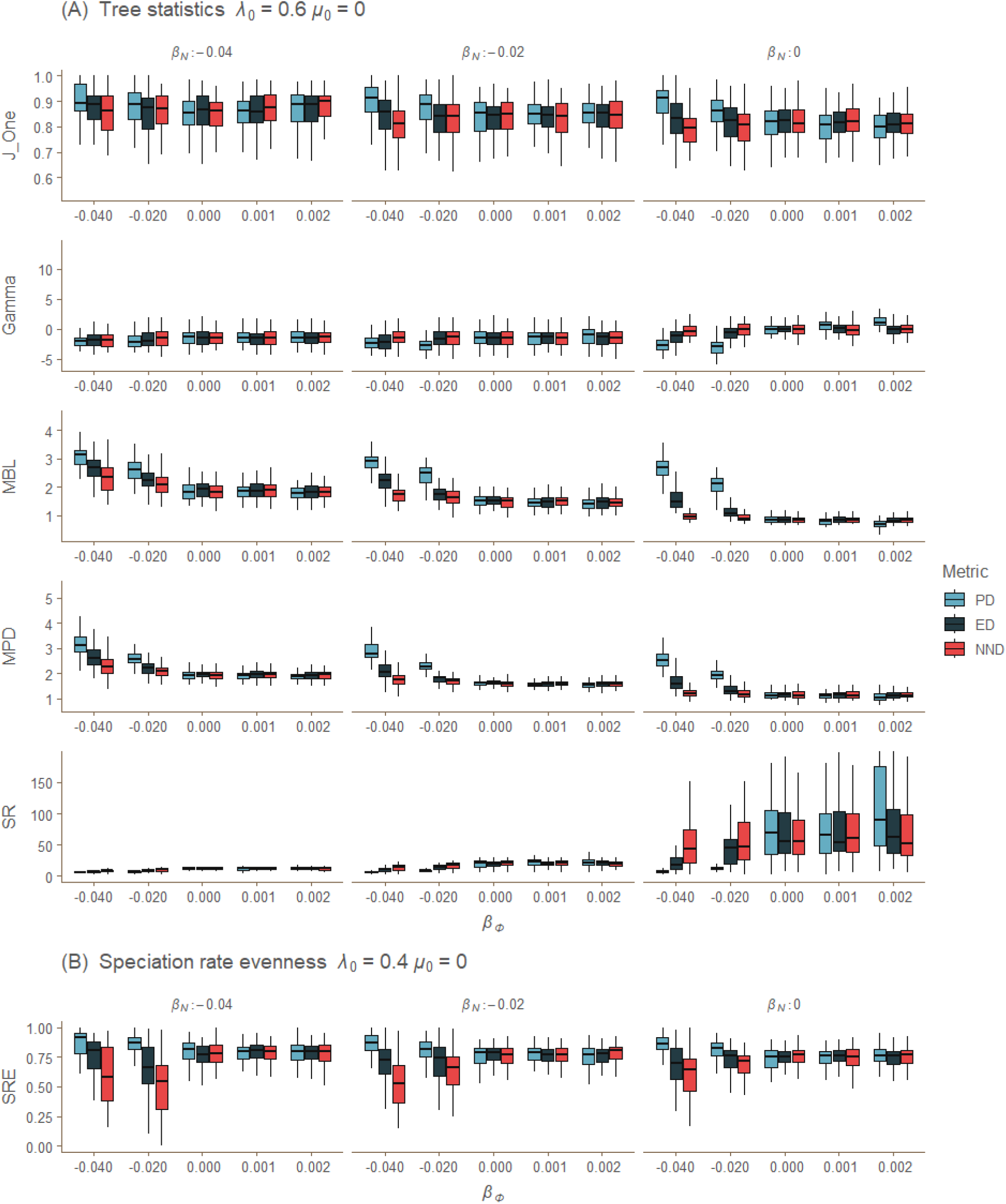
Tree summary statistics for the three scenarios of dependence of speciation rates (dependence on phylogenetic diversity (PD), evolutionary distinctiveness (ED) and nearest neighbor distance dependence (NND)), for various levels of the evolutionary relatedness effect (*β*_Φ_) and the species richness effect (*β*_N_). *λ*_0_ – initial speciation rate; *μ*_0_ - fixed extinction rate; x-axis: strength of ER effects; y-axis: value of the statistics. The following statistics are shown in panel A: J One - J One balance index; Gamma - Gamma statistic; MBL – the mean branch length; MPD - mean pairwise distance; SR - total number of extant lineages. The folloing statistic is shown in panel B: SRE - speciation rate evenness. SRE is shown in a separated panel because it has different patterns and is in a different parameter setting. We pick SRE from this parameter setting to better present the disparities among different scenarios and across different levels of the ER effect.

### Interaction between the effects of evolutionary relatedness and species richness

The effects of *β*_*Φ*_ (for instance the degree of disparities of the statistics between PD, ED and NND scenarios, and the changes of the statistics along with varying *β*_*Φ*_) appear more prominent when *β*_*N*_ imposes a less pronounced (i.e. less negative) effect on speciation, that is, the SR effect on the speciation rates is less pronounced, with the strongest effects observed at *β*_*N*_ = 0 which means the SR effect is neutral. From another point of view, the influence of *β*_*N*_ on the tree summary statistics is more prominent when *β*_*Φ*_ shifts towards more negative values (see Figure 4A).

When negative SR effect on speciation rate becomes weaker (*β*_*N*_ increases from - 0.04 to 0, SR effect on the speciation rates shifts from reduction to neutral), the trees become larger and less balanced, mean branch lengths and mean pairwise distances of the trees become smaller, the distribution of speciation events changes from being closer to the root to being more evenly distributed, and the degrees of disparities of speciation rate evenness among PD, ED and NND increase (see Figure 4A for details and see all the animations in Appendix A starting with beta n for animated comparisons). Generally, stronger negative SR effect reduces the differences of tree statistics among PD, ED and NND scenarios, it also reduces the differences of the statistics between different *β*_*Φ*_ settings. This is particularly true for the Gamma statistic and J One balance index.

The above described patterns, however, do not apply to speciation rate evenness. When the negative effect of SR on speciation rate becomes stronger (*β*_*N*_ decreases from 0 to -0.04), the disparities of speciation rate evenness among PD, ED and NND scenarios appear more prominent when *β*_*Φ*_ value becomes more negative.

### Effects of intrinsic speciation and extinction rates

Higher intrinsic speciation rates (*λ*_0_) lead to less balanced phylogenetic trees and more disparities among the scenarios. Higher intrinsic extinction rates (*µ*_0_), however, lead to more balanced trees and less disparities. The effects of *λ*_0_ and *µ*_0_ are intertwined with both *β*_*Φ*_ and *β*_*N*_. The effects are described in detail in Appendix E. Besides Figure 4, we also produced a set of animated plots (Appendix A), showing how tree summary statistics we measured change along with *β*_*Φ*_, *β*_*N*_, *λ*_0_ and *µ*_0_, to help interpret the patterns resulting from different levels of evolutionary relatedness effects (as determined by *β*_*Φ*_) and between the three different versions of ER dependence.

## Discussion

### From clade-wide to lineage-specific: ecological limits shape evolutionary trajectories

Evidence for ecological limits on diversity has been found or suggested in many studies (Rabosky, 2009a; Mittelbach et al., 2007), and ecological limits may exert influence on diversification by directly impacting rates of speciation (Wiens, 2011). Our simulations further suggested that, from clade-wide to lineage-specific, different extents of ecological limits result in unique phylogenetic tree properties, when using ER as a proxy.

When phylogenetic diversity acts as a proxy regulating speciation rate of all lineages (PD scenario), a negative ER effect (or *β*_*Φ*_, the corresponding parameter in our simulations) can be interpreted as a niche space limitation on clade expansion as phylogenetic diversity increases, which applies equally to all species in the clade. Under this interpretation, an increase in phylogenetic diversity within a clade leads to saturation of niches. Conversely, a positive ER effect provides the clade with the potential to exploit additional resources as phylogenetic diversity increases, potentially resulting in a rapid accumulation of species as the potential for biotic interactions increases.

By relating (clade-wide, PD scenario) speciation rate to ER, the PD scenario exhibits similarities to models with diversity carrying capacity (e.g. diversity-dependent diversification model, DDD), with negative ER effect (negative *β*_*Φ*_) on the speciation rates imposing an upper limit of phylogenetic diversity carrying capacity. The role of negative SR effect (negative *β*_*N*_) in our model is similar to the concept of carrying capacity in the DDD model, negative ER and SR effects in our simulations both markedly reduce average tree sizes, given a constant simulation time.

In the ED and NND scenarios, a negative ER effect indicates a scenario where more distantly related species are less likely to speciate, which may reflect speciation through an adaptive dynamics scenario where competition for similar resources leads to trait divergence to escape a fitness value (Geritz et al., 1999; Bolnick and Fitz-patrick, 2007). It may also reflect environmental filtering (Thakur and Wright, 2017), where closely related species matching the environment are more likely to speciate. In contrast, a positive ER effect indicates that more distantly related species are more likely to produce descendants, for example, due to species adapting to different environmental conditions and thereby exploring new niche space (Kozak and Wiens, 2006).

By relating lineage-specific speciation rates to ER, the ED and NND scenarios can account for evolutionary trajectories in which niches are either conserved (more similar than expected), constrained (diverging within a restricted range of available niches), or divergent (less similar than expected), based on different settings of ER effects (see the illustration of niche conservatism by Pyron et al. (2015)). Negative ER effect also trims the average tree sizes in the ED scenario, but this effect seems diminished or even absent in the NND scenario. Negative effects of ER decrease from PD scenario to ED scenario to NND scenario; along this “gradient” of metrics, ecological limits exert increasingly smaller influence on overall species richness of the communities, due to their impacts being more concentrated to close relatives (see also the speciation rate transitioning in Figure 3).

The unevenness of speciation rates among the tips increases from PD to ED and NND (note that although in PD scenario all tips of a tree have the same speciation rate, PD trees from the same parameter setting could still have different unevenness due to different tree topologies). When the ER effect is negative, the more distinct lineages are less likely to spawn new lineages, leading to a “clustering” where closer lineages have closer speciation rates. As the effect of ER goes from PD to ED to NND, large “clusters” break apart into small and separate clades on the tree, thereby increasing unevenness. When we shift the ER effect to positive, more distinct lineages are more prone to speciate, which may cause an “overdispersion” of the rates rather than a “clustering” (Figure 4B). In the ED and NND scenarios, the speciation rates among the tips are directly determined by their distances to the clade (ED) or their immediate neighbors (NND), so the observed unevenness could also be interpreted as an unevenness of ER.

In general, we observed hierarchical patterns in various summary statistics (e.g. J One balance index, Gamma statistic and speciation rate evenness) from clade-wide to lineage-specific ecological limits (scenarios, i.e. PD, ED, NND) when ER effect is negative. These patterns fade as ER effect shifts from negative to neutral (*β*_*Φ*_ = 0). We did not symmetrically explore the positive ER effects due to computational limitations as explained above.

We observed prominent difference of tree sizes between scenarios, with PD trees being the smallest, ED trees being noticeably larger and NND trees being the largest on average. Tree size often shows strong correlation with phylogenetic tree statistics, even after applying size correction methods, regardless of the underlying model (Janzen, 2023). This may explain why *β*_*Φ*_ impacts the balance and mean pairwise distances of PD trees, even though speciation rates across lineages are consistently uniform. To verify if tree size confounds our interpretation, we compared all the statistics from our results with the tree size correlation curves by Janzen (2023). We found that the correlations between statistics and tree size in our results are opposite to patterns observed by Janzen (2023) as tree size increases from 10 to 100. This implies that our findings are unlikely to be due to an effect of tree size. Rather, if the effect of tree size would be corrected for, we would expect our patterns to be even more pronounced.

### Species richness effect reduces evolutionary relatedness signature

Debates in ecology persist regarding the efficacy of phylogenetic metrics serving as proxies for species richness or functional diversity (Mazel et al., 2018; Owen et al., 2019). Our model introduces a more flexible scenario wherein species richness and phylogenetic (or evolutionary) relatedness operate concurrently. Their impacts on species diversification can be independently positive, neutral, or negative. We observed a stronger impact from ER as SR effect becomes neutral (*β*_*N*_ shifts from negative values to 0) on speciation rate in communities, with the most pronounced ER impact occurring when the SR effect is neutral (*β*_*N*_ = 0), where SR has no influence on species diversification. Furthermore, we noted that as SR imposes a more negative effect, the disparities in various tree statistics (e.g., J One balance index, mean branch length, mean pairwise distance, and Gamma statistic) among PD, ED, and NND scenarios become less evident or even disappear. Based on these patterns, we suggest that when speciation rate is limited by species richness, the signature of evolutionary relatedness is concealed. However, evolutionary relatedness can still play a complementary role in explaining macroevolutionary patterns if the impact of species richness is minor and the impact of evolutionary relatedness is substantial.

### More diverse evolutionary trajectories, increased tree imbalance

Imbalanced phylogenies are frequently observed in empirical research, and their occurrence has been attributed to various factors, including errors in phylogenetic data, incomplete species sampling, and biases introduced by reconstruction methods; additionally, such imbalance may reflect variations in evolutionary rates within trees (Stam, 2002; Blum and François, 2006). Simulations based on earlier stochastic models often show discrepancies when compared to empirical evidence, highlighting a potential misalignment between model predictions and actual observations (Blum and François, 2006), although there is significant variation in the degree of imbalance between empirical clades (Janzen and Etienne, 2024).

Our model offers more diverse evolutionary trajectories including lineage-specific scenarios that allow each lineage within a tree to have distinct speciation rates. Our analysis reveals notable differences in speciation rate variations within trees across different scenarios (see Figure 4B, the differences in speciation rate evenness), from PD through ED to NND. This hierarchical pattern reflects a shift in the impact of evolutionary relatedness–becoming increasingly concentrated among closer relatives– and a corresponding increase in speciation rate variation among lineages. Phylogenies simulated under our model can be more balanced or less balanced compared to a simple birth-death scenario. For example, in the NND scenario where trees often exhibit large speciation rate variation when ER imposes negative effect, stronger negative ER effect results in higher speciation rate variation and the corresponding trees exhibit greater imbalance. See Appendix C for a comparison of phylogenetic imbalance between empirical trees and trees simulated under our model.

### Phylogenetic diversity-dependence of extinction and empirical application

Although macroevolutionary dynamics are often studied through dependence of speciation rates on (phylogenetic) diversity metrics, the speed at which species go extinct may also be linked to ecological factors. Competition may play a non-negligible role in increasing the extinction risk of species, but its effects can vary depending on the context. For example, empirical evidence from Bengtsson (1989) on rock pool zooplankton demonstrates that interspecific competition increases local extinction rates; Dangremond et al. (2010) found that the proximity of an invasive grass increased seed predation on an endangered lupine species and accelerated its decline; Timmermann (2020) reviewed the extinction of Neanderthals and highlighted how competitive pressures, coupled with resource exploitation efficiency, played a significant role in Neanderthals’ demise. Overall, competition for finite ecological resources can accelerate species extinction, particularly for species that are more vulnerable to antagonistic ecological interactions.

If ER-dependent extinction were incorporated in our simulation, we expect that under the PD scenario, negative ER effects should lead to larger trees due to decreasing community-wide extinction rates, similar to the positive feedback loop of ER-dependent speciation. Positive ER effects on extinction would likely result in smaller trees. For the ED scenario, negative ER effects would mean distinct species are less likely to go extinct, making trees less balanced, while positive ER effects would promote the extinction of distinct species, leading to an auto-balancing process. For the NND scenario, ER effects are more local, which may result in different effects than for PD or ED.

Enabling both ER-dependent speciation and extinction, with all possible positive/negative combinations, could lead to even more diverse evolutionary trajectories. However, without data, drawing concrete implications would be challenging. We stress that increased extinction can eliminate some of the information contained in phylogenies, which would further burden our interpretation from phylogenetic trees.

In some real-world scenarios, it is possible that ER effects are so strong that SR is not dominating, which could explain the significant imbalance observed in empirical phylogenies, even when SR effects might otherwise obscure this pattern. However, fitting complex models to empirical phylogenies, such as through maximum-likelihood estimation or approximate Bayesian computation, presents mathematical and technical challenges (Etienne et al., 2014; Xie et al., 2023). Alternative methods such as neural networks can directly infer parameters from empirical phylogenies, but as models grow more complex, it becomes more difficult to recover parameters accurately, regardless of the method used (Qin et al., 2024). Nonetheless, it should still be possible to identify which ER scenarios generate evolutionary patterns that are closest to empirical data through neural network classification tasks. Such analyses may enable us to re-evaluate how ecological factors shape biodiversity.

## Conclusion

We have introduced a novel perspective on the influence of ecological limits, represented by the effects of evolutionary relatedness on species diversification patterns, which lead to a variety of macro-evolutionary trajectories. Our results show that the effects of ecological limits associated with phylogenetic distance between species are highly dependent on the scale at which these limits operate, as well as whether the effects of evolutionary relatedness on speciation rates are positive or negative. When operating at a smaller scale (branch or lineage specific), the phylogenies are expected to be more imbalanced, as there is larger variation of speciation rates within trees.

We have also presented evidence of the complex interplay between the effects of species richness and evolutionary relatedness on diversification, suggesting that strong species richness effects can reduce the influence of evolutionary relatedness on species diversification, thereby complicating interpretations of diversity slowdowns in macroevolutionary studies. Our findings suggest that a re-evaluation of how these factors operate concurrently to shape biodiversity patterns is needed.

## Glossary

**ED** Evolutionary distinctiveness, a measure of how unique or distinct a species is in terms of its evolutionary history. ED is computed by summing the pairwise distances between each lineage and all other lineages, divided by the number of lineages minus one. A species with high ED has fewer close relatives and represents a larger amount of independent evolutionary history than a species with low ED (Cadotte and Davies, 2010).

**ER** Evolutionary relatedness, a measure of evolutionary relationships between species. In our study, ER can be measured as PD, ED or NND (see definitions below).

**Gamma** Gamma statistic, a measure of internal node distribution. The value of gamma is calculated based on the distribution of node heights. A negative gamma value suggests a decrease in diversification rate over time, while a positive gamma value indicates an increase in diversification rate (Pybus and Harvey, 2000).

**J One** J One balance index, a measure of the degree of balance of a phylogeny. It is defined for trees with any degree distribution, and enables meaningful comparison of trees with different numbers of tips (Lemant et al., 2022).

**MBL** Mean branch length, a measure of average evolutionary change or divergence represented in a phylogenetic tree. It is calculated by summing the lengths of all the branches in a phylogenetic tree and dividing by the total number of branches (Webb et al., 2002).

**MPD** Mean pairwise distance, a measure of the average pairwise phylogenetic distance between species in a clade. It calculates the average of all pairwise distances between species in a phylogenetic tree, providing a measure of the overall phylogenetic spread (Webb et al., 2002).

**NND** Nearest neighbor distance, a measure of how distinct a species is in terms of its evolutionary history with respect to the most closely related species. NND is computed by taking the branch length distance on a phylogenetic tree from one species to its nearest neighbor on the tree. While ED measures distinctiveness on a global phylogenetic level, NND measures it on a local phylogenetic level (Webb, 2000).

**PD** Phylogenetic diversity, a measure of biodiversity that takes into account the evolutionary relationships between species. Specifically, PD is the minimum total length of all the phylogenetic branches required to span a given set of taxa on the phylogenetic tree. It provides insight into the evolutionary history represented by a group of species (Faith, 1992).

**Rogers** Rogers balance index, a measure of “patristic distance”, which represents the sum of branch lengths between two taxa in a phylogenetic tree which reflects the total amount of change along the evolutionary path between two taxa (Rogers, 1996).

**SR** Species richness, a measure of the number of different species present in a particular area or ecological community. It is a simple count of species.

## Appendix A Animation of main simulation

Animated plots were generated to aid the explanation of model behaviors and phylogenetic patterns of the main simulation. The plots were named in a format of “parameter statistic.gif” where parameter refers to the changing parameter in the animation and statistic refers to the summary statistic the plot shows. For example, “beta n Gamma.gif” is an animation showing the Gamma statistics among different parameter settings and transitioning in *β*_*N*_.

## Appendix B Heatmaps

Heatmaps were plotted to visualize the correlation matrix of a number of tree statistics under different ER scenarios. A representative tree was selected based on the heatmaps, using the statistics that are less correlated and more evenly spaced on the heatmap. There are three heatmaps, each presenting one scenario.

## Appendix C Tree imbalance

**Figure C5:**
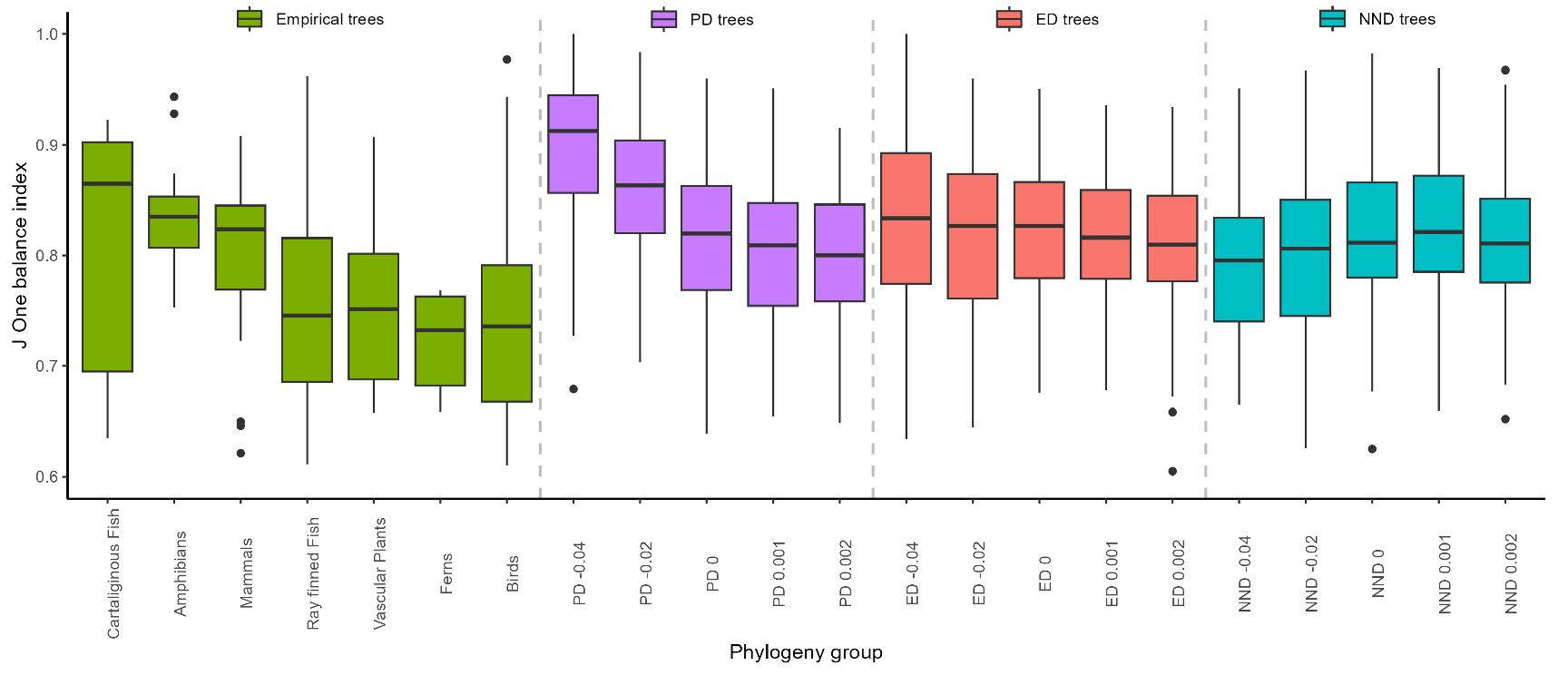
Comparison of phylogenetic tree imbalance between empirical clades and clades simulated using our model. The y-axis shows the J One balance index, with lower values indicating higher degree of imbalance. The leftmost group of boxes shows the tree imbalance computed from empirical trees obtained from Janzen and Etienne (2024). The x-axis indicates the taxonomic group of the empirical trees; there are seven empirical sub-clades. In the remaining groups of boxes, the x-axis indicates the ER scenario (PD, ED, or NND) and the effect size of ER (*β*_*Φ*_) used for simulating the trees. The remaining parameters are fixed: speciation rate *λ*_0_ = 0.6, extinction rate *µ*_0_ = 0 and effect size of SR *β*_*N*_ = 0. The plots show that under certain parameter values, we can generate trees with higher levels of imbalance, bringing the simulated phylogenies closer to empirical phylogenies. For example, with a net diversification rate (*λ*_0_ *− µ*_0_) = 0.6, *β*_*Φ*_ *>* 0.001, and *β*_*N*_ = 0, we can generate phylogenies that are more imbalanced than those of mammals and amphibians under the PD and ED scenarios. Further increasing the speciation rate and *β*_*Φ*_ could yield even more imbalanced trees. Under the NND scenario we estimate that *β*_*Φ*_ *< −*0.01 can produce phylogenies close to empirical trees. We observed substantial variation in imbalance between empirical clades, with some clades, such as ferns and birds, much more imbalanced, while others, such as cartilaginous fish, more balanced. Given the smaller sample size and potential biases in empirical phylogenies, these findings should be interpreted with caution.

## Appendix D Tree statistics and LTT plots with representative trees

Tree statistics and LTT plots with representative trees were plotted for all the parameter settings of the main simulation. The plots were named in a format of “*λ*_0_ *µ*_0_ *β*_*N*_ .png”. For example, “0.6 0 0.png” presents the results under the parameter setting *λ*_0_ = 0.6, *µ*_0_ = 0 and *β*_*N*_ = 0.

## Appendix E Effects of intrinsic speciation and extinction rates

We found that the intrinsic speciation rate (*λ*_0_) also showed a substantial impact on tree statistics. The effects of this parameter are intertwined with both *β*_*Φ*_ and *β*_*N*_ (see animations in Appendix A starting with lambda and mu).

As *λ*_0_ increases, the differences in tree balance across scenarios increase, particularly when *β*_*Φ*_ has smaller positive effect or the effect is more negative. The trees become generally less balanced as *λ*_0_ increases. The effects of *λ*_0_ are stronger when *β*_*Φ*_ is more negative and *β*_*N*_ is closer to zero, with the most profound case being *β*_*Φ*_ = *−*0.04 and *β*_*N*_ = 0 where increasing *λ*_0_ results in most evident increase of disparities of balance between PD, ED and NND scenarios (see the animations in Appendix A starting with lambda and ending with J One).

In contrast to *λ*_0_, an increase in the intrinsic extinction rate (*µ*_0_) results in more balanced trees and less disparities of balance among the PD, ED and NND scenarios. The reduction of disparities is more prominent when *β*_*Φ*_ becomes more negative. The most prominent case is when *β*_*Φ*_ = *−*0.04 and *β*_*N*_ = 0 where increasing *µ*_0_ results in most evident reduction of disparities of balance between PD, ED and NND scenarios (see the animations in Appendix A ending with J One) (see the animations in Appendix A starting with mu and ending with J One).

An increase in *λ*_0_ from 0.4 to 0.6 generally leads to a reduction in mean branch length and mean pairwise distance, while an increase in *µ*_0_ from 0 to 0.2 generally leads to an increase in these statistics (see the animations in Appendix A starting with lambda and mu, ending with MPD and MBL).

The disparities of speciation events distributed across the phylogeny among PD, ED and NND scenarios, as measured by the gamma statistic, become more prominent as *λ*_0_ increases and less prominent as *µ*_0_ increases. The distribution of speciation events changes from being closer to the root to being more evenly distributed as *µ*_0_ increases (see the animations in Appendix A starting with lambda and mu, ending with Gamma).

When *β*_*Φ*_ effect is negative, increasing *λ*_0_ has the largest effect of increasing the tree sizes in the NND scenario. This effect is smaller in the ED scenario and even smaller in the PD scenario. Increasing *µ*_0_ has the largest effect of reducing the tree sizes in the NND scenario; this effect is smaller in the ED scenario and even smaller in the PD scenario. When *β*_*Φ*_ effect is zero or positive, these trends are diminished (see the animations in Appendix A starting with lambda and mu, ending with SR). Increasing *λ*_0_ only has the effect of reducing the disparities of speciation rate evenness among PD, ED and NND when *β*_*Φ*_ is negative and *β*_*N*_ is 0. *µ*_0_ has no obvious effect on the evenness of the speciation rates (see Figure 4B and the animations in Appendix A starting with lambda and mu, ending with ERE).

## Appendix F Correlation Matrix in Ultrametric Phylogenetic Trees

This section presents a proof of the relationship between the correlation matrix and the phylogenetic distance matrix for any ultrametric phylogeny under the Brownian motion model of trait evolution (Felsenstein, 1985; Martins and Hansen, 1997). Specifically, we prove that the correlation matrix ***C*** is given by

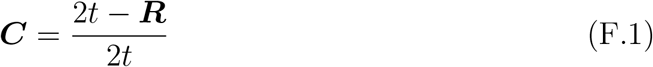

where *t* is the crown age of the phylogeny, and ***R*** is the phylogenetic distance matrix.

### Definition 1

(Phylogenetic Tree). *A phylogenetic tree is denoted as 𝒯* = (*𝒩, ε*), *where 𝒩 is the set of nodes, including the root, internal nodes, and tips (leaves) representing taxa, and E is the set of edges (branches) connecting the nodes. Each edge e ∈ E has an associated branch length l*_*e*_ *>* 0.

### Definition 2

(Ultrametric Tree). *An ultrametric tree is a rooted phylogenetic tree in which all tips (leaves) are equidistant from the root. This implies that the total path length from the root to any tip is the same for all tips*.

### Definition 3

(Brownian Motion Model). *In the Brownian motion model of trait evolution, the variance of trait change along a branch of length l*_*e*_ *is σ*^2^*l*_*e*_, *where σ*^2^ *is the evolutionary rate. Trait changes along different branches are independent unless they share common ancestry*.

### Definition 4

(Variance of a Trait). *For each tip i ∈ℒ, let P*_*i*_ *denote the set of edges (branches) on the path from the root to tip i. The variance of a neutral trait X*_*i*_ *at tip i is defined as*

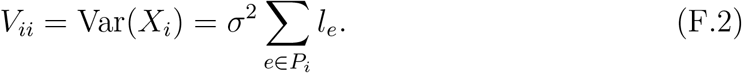

### Definition 5

(Covariance Between Traits). *The covariance between traits at tips I and j is defined as*

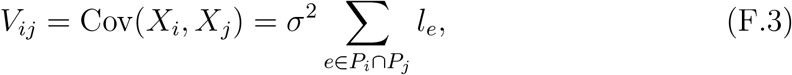

*where P*_*i*_ *∩ P*_*j*_ *is the set of shared edges on the paths from the root to tips i and j*.

### Definition 6

(Correlation Coefficient). *The correlation coefficient between tips i and j is defined as*

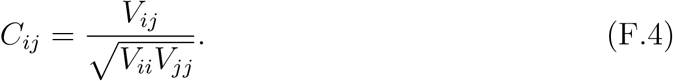

### Definition 7

(Phylogenetic Distance). *The phylogenetic distance R*_*ij*_ *between tips I and j is defined as*

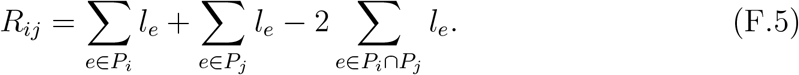

*By virtue of F*.*2 this simplifies to:*

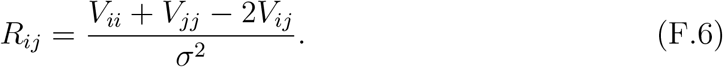

**Theorem**. *For any ultrametric phylogeny under the Brownian motion model, the correlation matrix* ***C*** *is given by:*

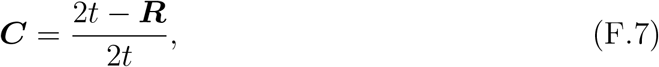

*where t is the crown age of the phylogeny, and* ***R*** *is the phylogenetic distance matrix*.

*Proof*. We begin by rearranging Equation F.6 as

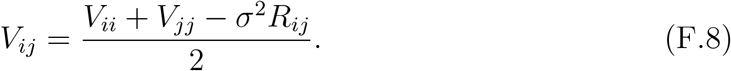

Substituting Equation F.8 into the correlation coefficient formula in Equation F.4 leads to:

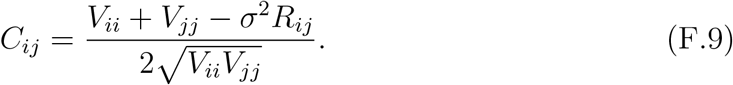

Because the tree is ultrametric, the total path length from the root to any tip *i* equals the crown age *t*. Hence, for any tip *i* and *j*, recall from Equation F.2, we get:

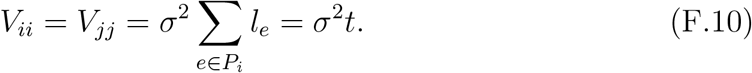

Substituting *V*_*ii*_ = *V*_*jj*_ = *σ*^2^*t* into Equation F.9 leads to

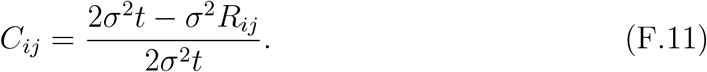

Then we factor out *σ*^2^:

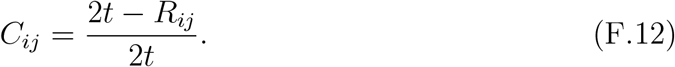

Because *C*_*ij*_ denotes all the elements in the correlation matrix ***C***, and *R*_*ij*_ denotes all the elements in the phylogenetic distance matrix ***R***, Equation F.12 implies:

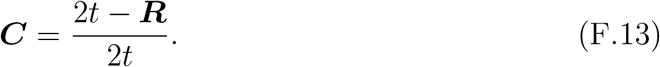

This completes the proof.

## Appendix G Simplification of the speciation rate evenness computation

In this section we continue with the simplification of Equation 4. For binary trees, the number of tips *n ≥* 2. If diag(***R***) represents the vector comprising the diagonal elements of ***R***, by definition of ***R***, all elements in diag(***R***) are equal to 0, thus all elements in diag(***C***) are equal to 1. Then we have

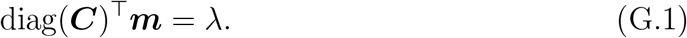

Recall that

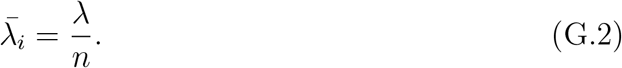

So Equation 4 can be rewritten as

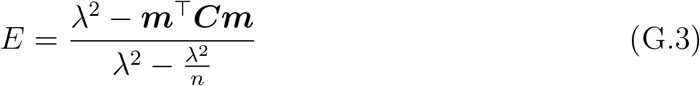

Factoring out *λ*^2^ leads to

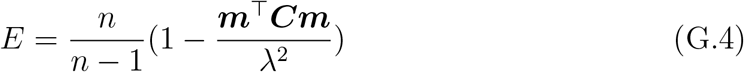

Defining***m***^*′*^ = ***m****/λ*, we get

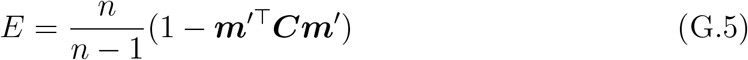

## References

Aguilée, R., Gascuel, F., Lambert, A., Ferriere, R., 2018. Clade diversification dynamics and the biotic and abiotic controls of speciation and extinction rates. Nature Communications 9, 1–13.

Bengtsson, J., 1989. Interspecific competition increases local extinction rate in a metapopulation system. Nature 340, 713–715. URL: https://www-nature-com.proxy-ub.rug.nl/articles/340713a0, doi:10.1038/340713a0.

Blum, M.G.B., François, O., 2006. Which Random Processes Describe the Tree of Life? A Large-Scale Study of Phylogenetic Tree Imbalance. Systematic Biology 55, 685–691. URL: 10.1080/10635150600889625, doi:10.1080/10635150600889625.

Bolnick, D.I., Fitzpatrick, B.M., 2007. Sympatric speciation: models and empirical evidence. Annual Review of Ecology, Evolution, and Systematics 38, 459–487.

Cadotte, M.W., Davies, T.J., 2010. Rarest of the rare: advances in combining evolutionary distinctiveness and scarcity to inform conservation at biogeographical scales. Diversity and Distributions 16, 376–385.

Cadotte, M.W., Dinnage, R., Tilman, D., 2012. Phylogenetic diversity promotes ecosystem stability. Ecology 93, S223–S233.

Dangremond, E.M., Pardini, E.A., Knight, T.M., 2010. Apparent competition with an invasive plant hastens the extinction of an endangered lupine. Ecology 91, 2261–2271. URL: https://onlinelibrary-wiley-com.proxy-ub.rug.nl/doi/abs/10.1890/09-0418.1, doi:10.1890/09-0418.1.

De Maesschalck, R., Jouan-Rimbaud, D., Massart, D.L., 2000. The Mahalanobis distance. Chemometrics and Intelligent Laboratory Systems 50, 1–18.

Etienne, R.S., Haegeman, B., Dugo-Cota, A., Vilà, C., Gonzalez-Voyer, A., Valente, L., 2023. The Phylogenetic Limits to Diversity-Dependent Diversification. Systematic Biology 72, 433–445.

Etienne, R.S., Haegeman, B., Stadler, T., Aze, T., Pearson, P.N., Purvis, A., Phillimore, A.B., 2012. Diversity-Dependence Brings Molecular Phylogenies Closer to Agreement with the Fossil Record. Proceedings of the Royal Society B: Biological Sciences 279, 1300–1309.

Etienne, R.S., Morlon, H., Lambert, A., 2014. Estimating the Duration of Speciation from Phylogenies. Evolution 68, 2430–2440. URL: 10.1111/evo.12433, doi:10.1111/evo.12433.

Etienne, R.S., Rosindell, J., 2012. Prolonging the Past Counteracts the Pull of the Present: Protracted Speciation Can Explain Observed Slowdowns in Diversification. Systematic Biology 61, 204.

Faith, D.P., 1992. Conservation evaluation and phylogenetic diversity. Biological conservation 61, 1–10.

Felsenstein, J., 1985. Phylogenies and the Comparative Method. The American Naturalist 125, 1–15. URL: https://www.journals.uchicago.edu/doi/10.1086/284325, doi:10.1086/284325.

Foote, M., Cooper, R.A., Crampton, J.S., Sadler, P.M., 2018. Diversity-dependent evolutionary rates in early Palaeozoic zooplankton. Proceedings of the Royal Society B: Biological Sciences 285, 20180122.

Gerhold, P., Cahill, J.F., Winter, M., Bartish, I.V., Prinzing, A., 2015. Phylogenetic patterns are not proxies of community assembly mechanisms (they are far better). Functional Ecology 29, 600–614.

Geritz, S.A.H., van der Meijden, E., Metz, J.A.J., 1999. Evolutionary Dynamics of Seed Size and Seedling Competitive Ability. Theoretical Population Biology 55, 324–343. URL: https://www.sciencedirect.com/science/article/pii/S0040580998914095, doi:10.1006/tpbi.1998.1409.

Gillespie, D.T., 1976. A general method for numerically simulating the stochastic time evolution of coupled chemical reactions. Journal of Computational Physics 22, 403–434.

Helmus, M.R., Bland, T.J., Williams, C.K., Ives, A.R., 2007. Phylogenetic measures of biodiversity. The American Naturalist 169, E68.

Henn, J.J., Yelenik, S., Damschen, E., 2019. Environmental gradients influence differences in leaf functional traits between native and non-native plants. Oecologia 191, 397–409.

Hildenbrandt, H., Qin, T., 2024. evesim: Evolutionary relatedness dependent diversification simulation powered by the Rcpp backend SimTable. URL: https://github.com/EvoLandEco/evesim. xoriginal-date: 2024-09-17T07:04:41Z.

HilleRisLambers, J., Adler, P., Harpole, W., Levine, J., Mayfield, M., 2012. Rethinking Community Assembly through the Lens of Coexistence Theory. Annual Review of Ecology, Evolution, and Systematics 43, 227–248.

Janzen, T., 2023. Tree Size. URL: https://github.com/thijsjanzen/treestats/blob/master/articles/tree_size.html.

Janzen, T., Etienne, R.S., 2024. Phylogenetic tree statistics: A systematic overview using the new R package ‘treestats’. Molecular Phylogenetics and Evolution 200, 108168. URL: https://www.sciencedirect.com/science/article/pii/S105579032400160X, doi:10.1016/j.ympev.2024.108168.

Kondratyeva, A., Grandcolas, P., Pavoine, S., 2019. Reconciling the concepts and measures of diversity, rarity and originality in ecology and evolution. Biological Reviews 94, 1317–1337.

Kozak, K.H., Wiens, J., 2006. Does Niche Conservatism Promote Speciation? A Case Study in North American Salamanders. Evolution 60, 2604–2621.

Kubo, T., Iwasa, Y., 1995. Inferring the rates of branching and extinction from molecular phylogenies. Evolution 49, 694–704.

Lemant, J., Le Sueur, C., Manojlović, V., Noble, R., 2022. Robust, universal tree balance indices. Systematic biology 71, 1210–1224.

Louca, S., Pennell, M.W., 2021. Why extinction estimates from extant phylogenies are so often zero. Current Biology 31, 3168–3173. e4.

Martins, E.P., Hansen, T.F., 1997. Phylogenies and the Comparative Method: A General Approach to Incorporating Phylogenetic Information into the Analysis of Interspecific Data. The American Naturalist 149, 646– 667. URL: https://www-journals-uchicago-edu.proxy-ub.rug.nl/doi/abs/10.1086/286013, doi:10.1086/286013.

Mayfield, M.M., Levine, J.M., 2010. Opposing effects of competitive exclusion on the phylogenetic structure of communities. Ecology Letters 13, 1085–1093.

Mazel, F., Pennell, M.W., Cadotte, M.W., Diaz, S., Dalla Riva, G.V., Grenyer, R., Leprieur, F., Mooers, A.O., Mouillot, D., Tucker, C.M., Pearse, W.D., 2018. Prioritizing phylogenetic diversity captures functional diversity unreliably. Nature Communications 9, 2888.

Mittelbach, G.G., Schemske, D.W., Cornell, H.V., Allen, A.P., Brown, J.M., Bush, M.B., Harrison, S.P., Hurlbert, A.H., Knowlton, N., Lessios, H.A., McCain, C.M., McCune, A.R., McDade, L.A., McPeek, M.A., Near, T.J., Price, T.D., Ricklefs, R.E., Roy, K., Sax, D.F., Schluter, D., Sobel, J.M., Turelli, M., 2007. Evolution and the latitudinal diversity gradient: speciation, extinction and biogeography. Ecology Letters 10, 315–331.

Moen, D., Morlon, H., 2014. Why does diversification slow down? Trends in Ecology and Evolution 29, 190–197.

Nee, S., Holmes, E.C., May, R.M., Harvey, P.H., 1994. Extinction Rates Can Be Estimated from Molecular Phylogenies. Philosophical Transactions of the Royal Society of London. Series B: Biological Sciences 344, 77–82.

Nee, S., Mooers, A.O., Harvey, P.H., 1992. Tempo and mode of evolution revealed from molecular phylogenies. Proceedings of the National Academy of Sciences, USA 89, 8322–8326.

Owen, N.R., Gumbs, R., Gray, C.L., Faith, D.P., 2019. Global conservation of phylogenetic diversity captures more than just functional diversity. Nature Communications 10, 859.

Phillimore, A.B., Price, T.D., 2008. Density-Dependent Cladogenesis in Birds. PLoS Biology 6, e71.

Pie, M.R., Divieso, R., Caron, F.S., 2023. Clade density and the evolution of diversity-dependent diversification. Nature Communications 14, 4576.

Pigot, A.L., Bregman, T., Sheard, C., Daly, B., Etienne, R.S., Tobias, J.A., 2016. Quantifying species contributions to ecosystem processes: a global assessment of functional trait and phylogenetic metrics across avian seed-dispersal networks. Proceedings of the Royal Society B: Biological Sciences 283, 20161597.

Pires, M.M., Silvestro, D., Quental, T.B., 2017. Interactions within and between clades shaped the diversification of terrestrial carnivores. Evolution 71, 1855–1864.

Purvis, A., 2008. Phylogenetic approaches to the study of extinction. Annual Review of Ecology, Evolution, and Systematics 39, 301–319.

Pybus, O.G., Harvey, P.H., 2000. Testing macro–evolutionary models using incomplete molecular phylogenies. Proceedings of the Royal Society B: Biological Sciences 267, 2267–2272.

Pyron, R.A., Costa, G.C., Patten, M.A., Burbrink, F.T., 2015. Phylogenetic niche conservatism and the evolutionary basis of ecological speciation. Biological Reviews 90, 1248–1262.

Qin, T., 2023. eve - Evolution emulator, phylogenetically dependent. URL: https://github.com/EvoLandEco/eve, doi:10.5281/zenodo.7991231.

Qin, T., Benthem, K.J.v., Valente, L., Etienne, R.S., 2024. Performance and Robustness of Parameter Estimation from Phylogenetic Trees Using Neural Networks. URL: https://www.biorxiv.org/content/10.1101/2024.08.02.606350v3, doi:10.1101/2024.08.02.606350. pages: 2024.08.02.606350 Section: New Results.

Qin, T., Zhou, J., Sun, Y., Müller-Schärer, H., Luo, F., Dong, B., Li, H., Yu, F., 2020. Phylogenetic diversity is a better predictor of wetland community resistance to Alternanthera philoxeroides invasion than species richness. Plant Biology 22, 591–599.

Quental, T.B., Marshall, C.R., 2010. Diversity dynamics: molecular phylogenies need the fossil record. Trends in Ecology and Evolution 25, 434–441.

Rabosky, D.L., 2009a. Ecological limits and diversification rate: alternative paradigms to explain the variation in species richness among clades and regions. Ecology Letters 12, 735–743.

Rabosky, D.L., 2009b. Heritability of Extinction Rates Links Diversification Patterns in Molecular Phylogenies and Fossils. Systematic Biology 58, 629–640. URL: 10.1093/sysbio/syp069, doi:10.1093/sysbio/syp069.

Rabosky, D.L., Glor, R.E., 2010. Equilibrium speciation dynamics in a model adaptive radiation of island lizards. Proceedings of the National Academy of Sciences, USA 107, 22178–22183.

Redding, D.W., Hartmann, K., Mimoto, A., Bokal, D., DeVos, M., Mooers, A.O., 2008. Evolutionarily distinctive species often capture more phylogenetic diversity than expected. Journal of theoretical biology 251, 606–615.

Ricklefs, R.E., 2010. Evolutionary diversification, coevolution between populations and their antagonists, and the filling of niche space. Proceedings of the National Academy of Sciences, USA 107, 1265–1272.

Rillo, M., Etienne, R., 2022. Diversity-Dependent Diversification. Oxford Bibliographies Online doi:10.1093/OBO/9780199941728-0141.

Rogers, J.S., 1996. Central Moments and Probability Distributions of Three Measures of Phylogenetic Tree Imbalance. Systematic Biology 45, 99–110.

Rosindell, J., Manson, K., Gumbs, R., Pearse, W.D., Steel, M., 2024. Phylogenetic Biodiversity Metrics Should Account for Both Accumulation and Attrition of Evolutionary Heritage. Systematic Biology 73, 158–182. URL: 10.1093/sysbio/syad072, doi:10.1093/sysbio/syad072.

Sepkoski, J.J., 1978. A kinetic model of Phanerozoic taxonomic diversity I. Analysis of marine orders. Paleobiology 4, 223–251.

Srivastava, D.S., Cadotte, M.W., Macdonald, A.A.M., Marushia, R.G., Mirotchnick, N., 2012. Phylogenetic diversity and the functioning of ecosystems. Ecology Letters 15, 637–648.

Stam, E., 2002. Does Imbalance in Phylogenies Reflect Only Bias? Evolution 56, 1292–1295. URL: https://onlinelibrary.wiley.com/doi/abs/10.1111/j.0014-3820.2002.tb01440.x, doi:10.1111/j.0014-3820.2002.tb01440.x.

Thakur, M.P., Wright, A.J., 2017. Environmental Filtering, Niche Construction, and Trait Variability: The Missing Discussion. Trends in Ecology and Evolution 32, 884–886.

Timmermann, A., 2020. Quantifying the potential causes of Neanderthal extinction: Abrupt climate change versus competition and interbreeding. Quaternary Science Reviews 238, 106331. URL: https://www.sciencedirect.com/science/article/pii/S0277379120302936, doi:10.1016/j.quascirev.2020.106331.

Tucker, C.M., Davies, T.J., Cadotte, M.W., Pearse, W.D., 2018. On the relationship between phylogenetic diversity and trait diversity. Ecology 99, 1473–1479.

Valentine, J.W., 1973. Evolutionary paleoecology of the marine biosphere. Englewood Cliffs, New Jersey.

Venail, P., Gross, K., Oakley, T.H., Narwani, A., Allan, E., Flombaum, P., Isbell, F., Joshi, J., Reich, P.B., Tilman, D., 2015. Species richness, but not phylogenetic diversity, influences community biomass production and temporal stability in a reexamination of 16 grassland biodiversity studies. Functional Ecology 29, 615–626.

Webb, C.O., 2000. Exploring the phylogenetic structure of ecological communities: an example for rain forest trees. The American Naturalist 156, 145–155.

Webb, C.O., Ackerly, D.D., McPeek, M.A., Donoghue, M.J., 2002. Phylogenies and community ecology. Annual review of ecology and systematics 33, 475–505.

Wiens, J.J., 2011. The causes of species richness patterns across space, time, and clades and the role of “ecological limits”. The Quarterly review of biology 86, 75–96.

Xie, S., Valente, L., Etienne, R.S., 2023. Can We Ignore Trait-Dependent Colonization and Diversification in Island Biogeography? Evolution 77, 670–681. URL: 10.1093/evolut/qpad006, doi:10.1093/evolut/qpad006.

Zheng, Y.L., Burns, J.H., Liao, Z.Y., Li, Y.P., Yang, J., Chen, Y.J., Zhang, J.L., Zheng, Y.G., 2018. Species composition, functional and phylogenetic distances correlate with success of invasive Chromolaena odorata in an experimental test. Ecology Letters 21, 1211–1220.

