## Supplementary figures and images for "Impact of Evolutionary Relatedness on Species Diversification and Tree Shape"

### Appendix A beta_n_ERE.gif

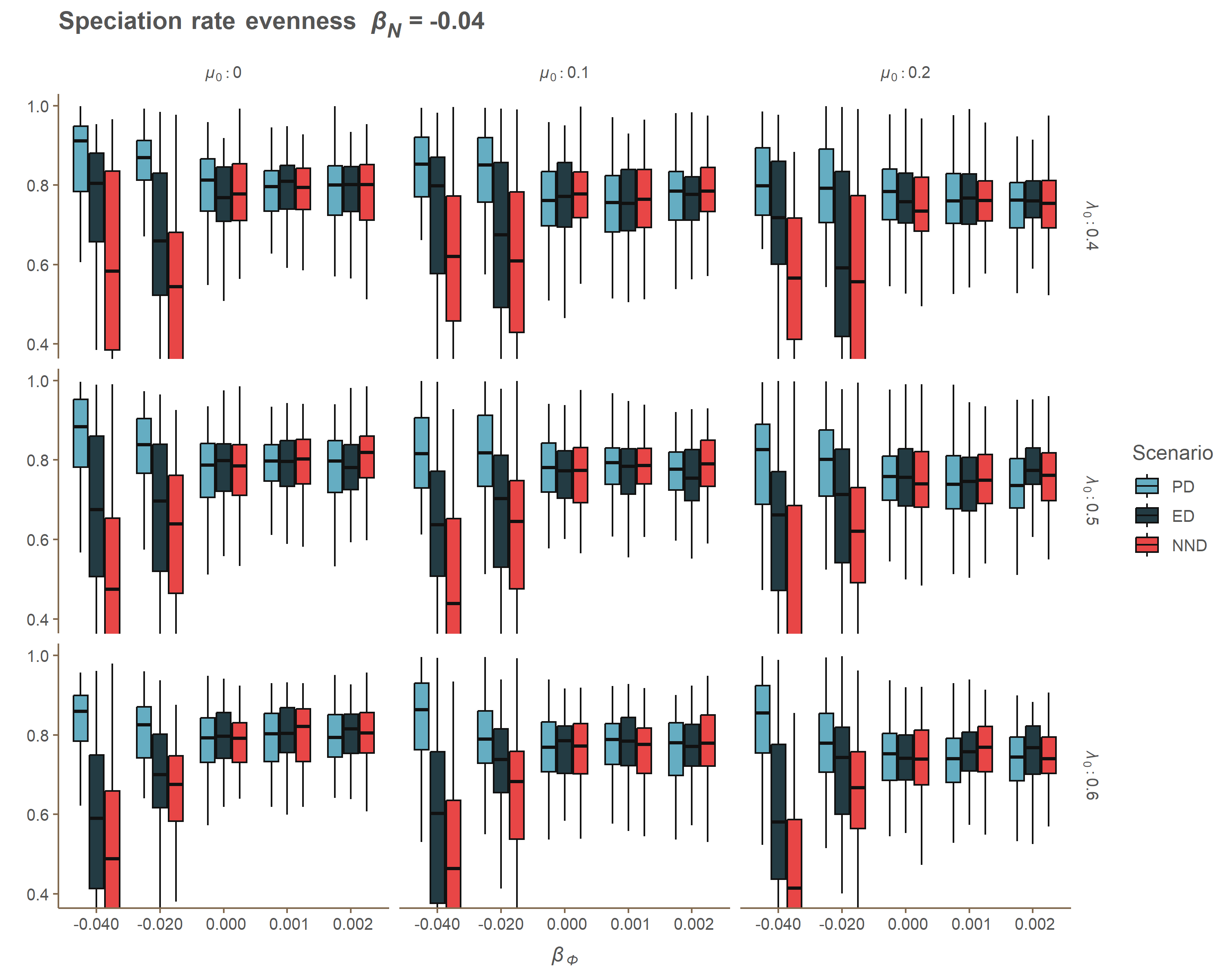

### Appendix A beta_n_Gamma.gif

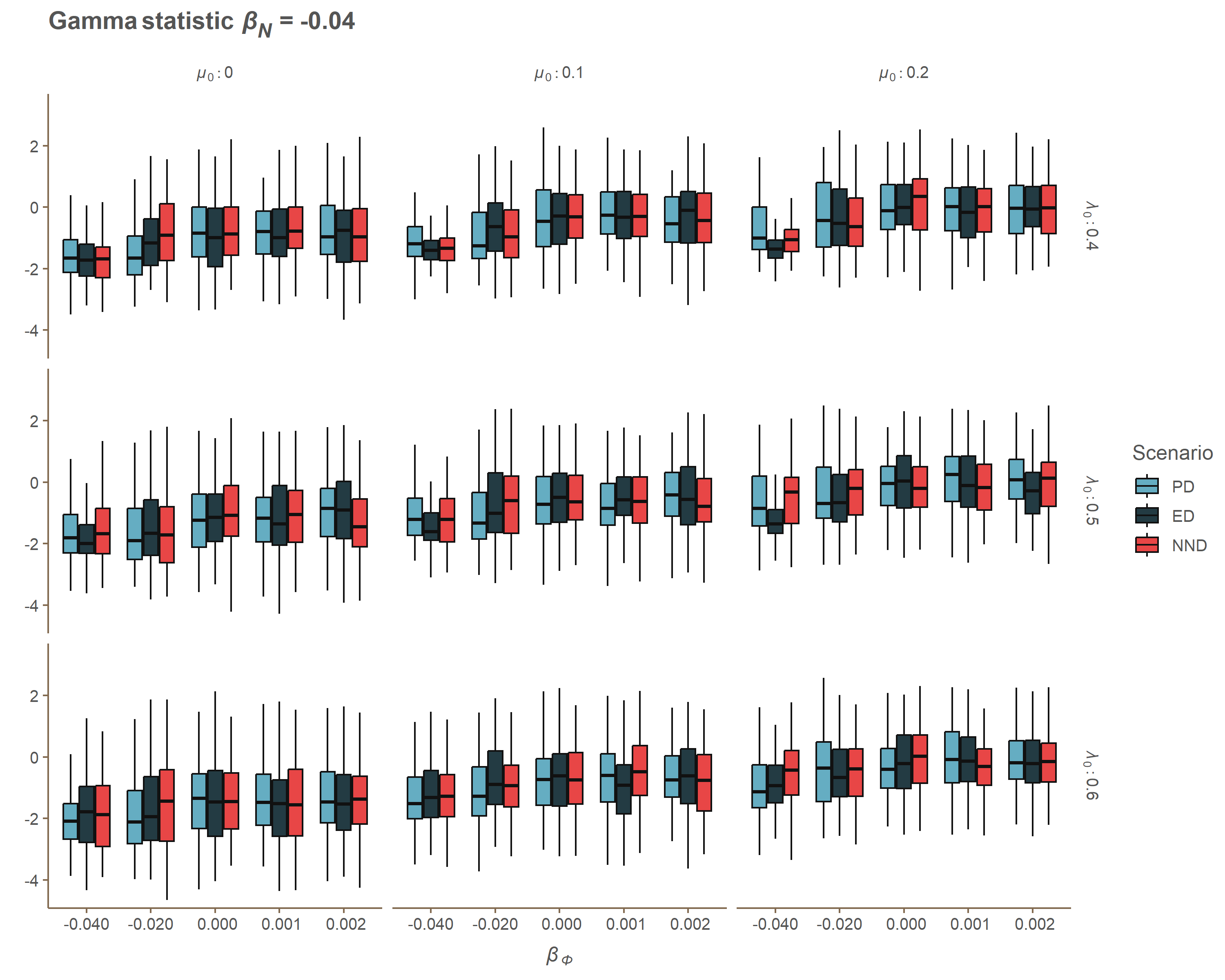

### Appendix A beta_n_J_One.gif

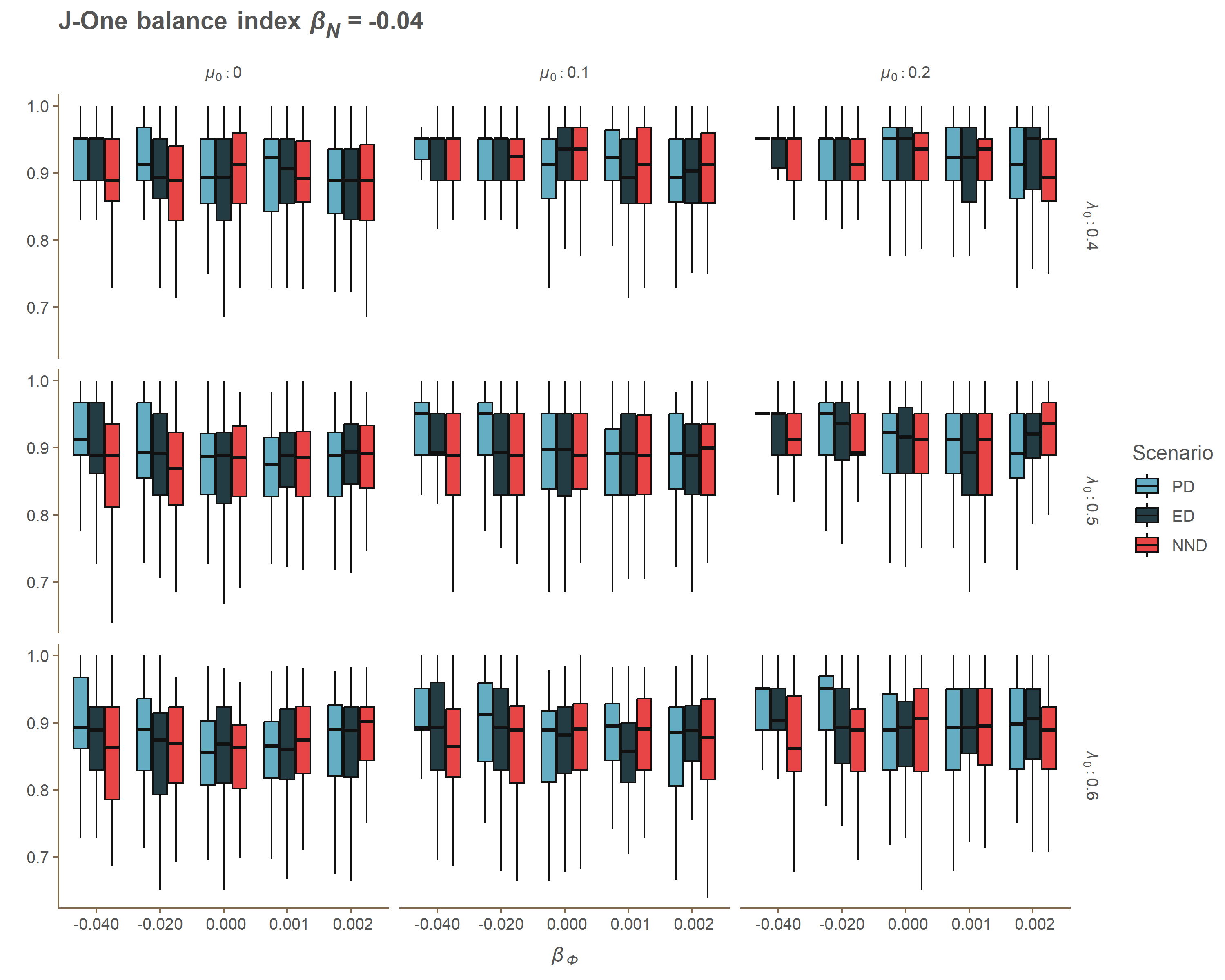

### Appendix A beta_n_MBL.gif

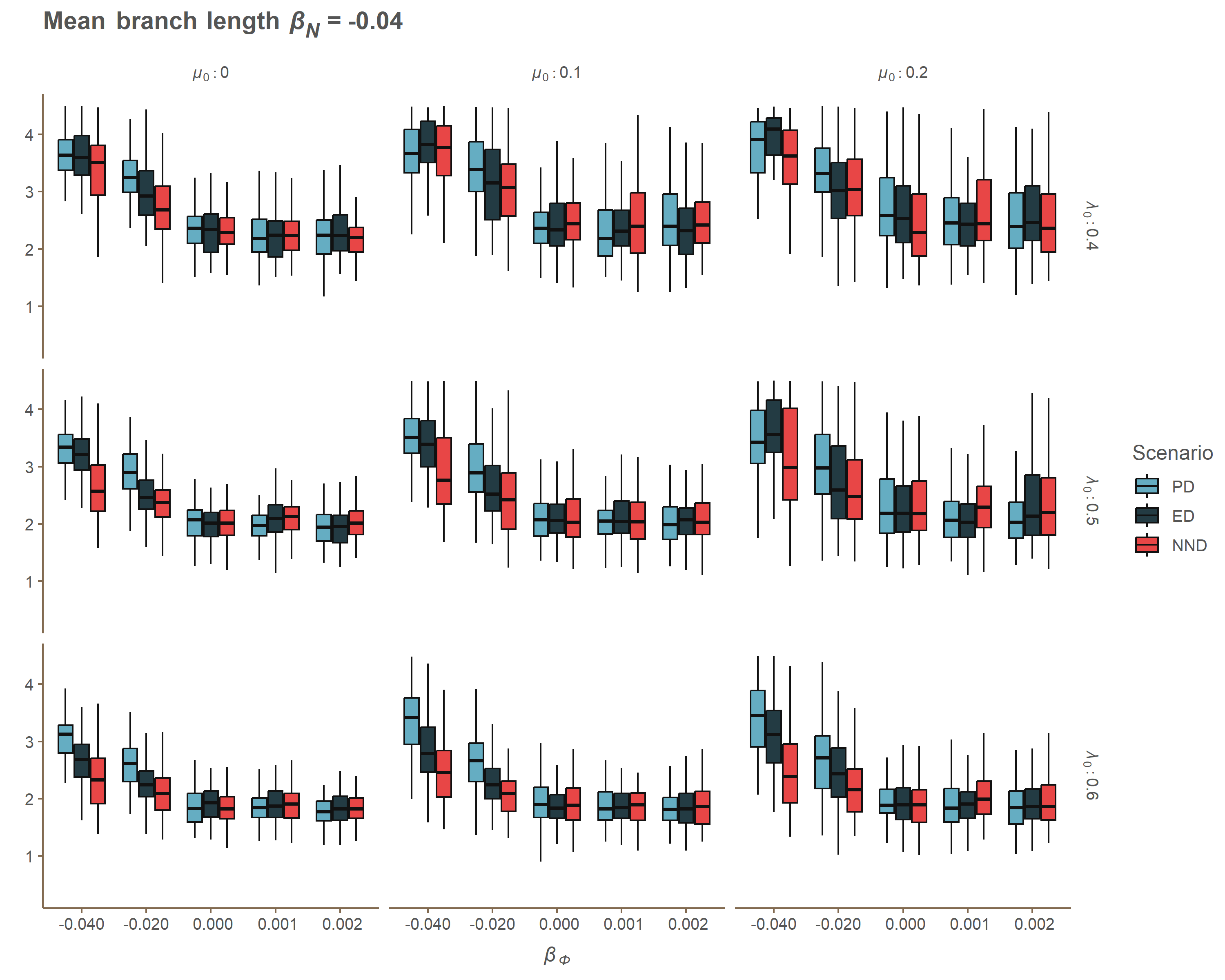

### Appendix A beta_n_MNTD.gif

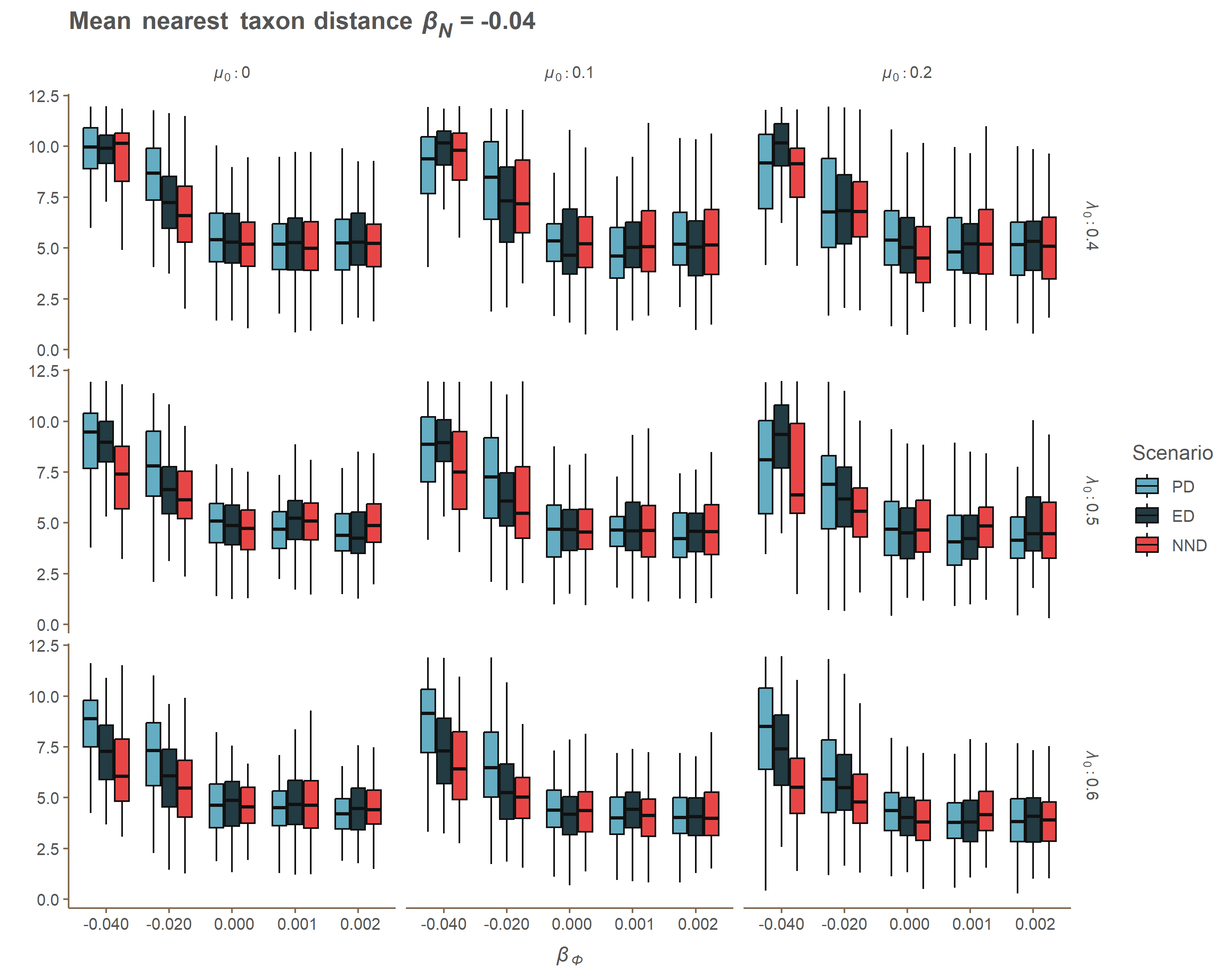

### Appendix A beta_n_MPD.gif

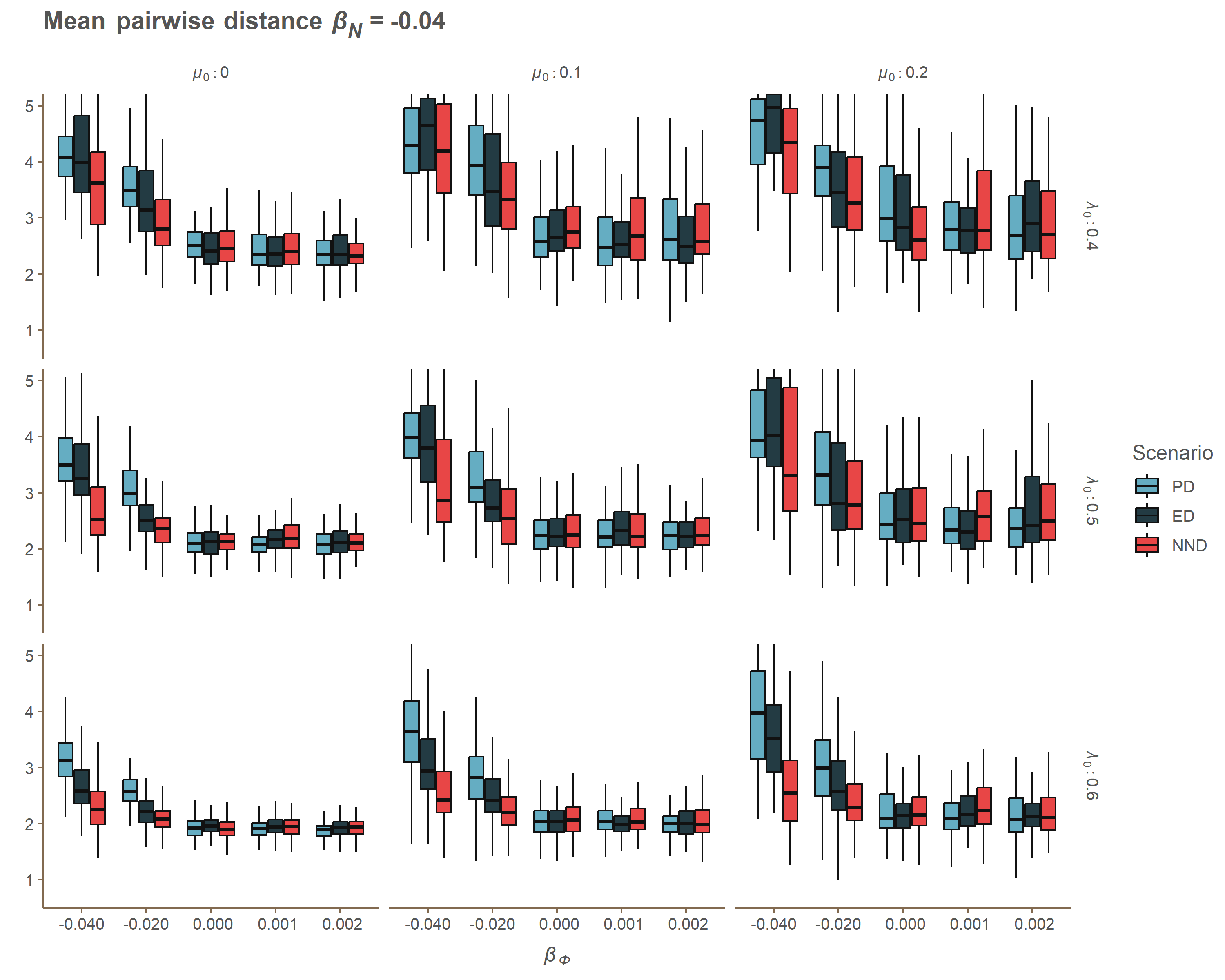

### Appendix A beta_n_PD.gif

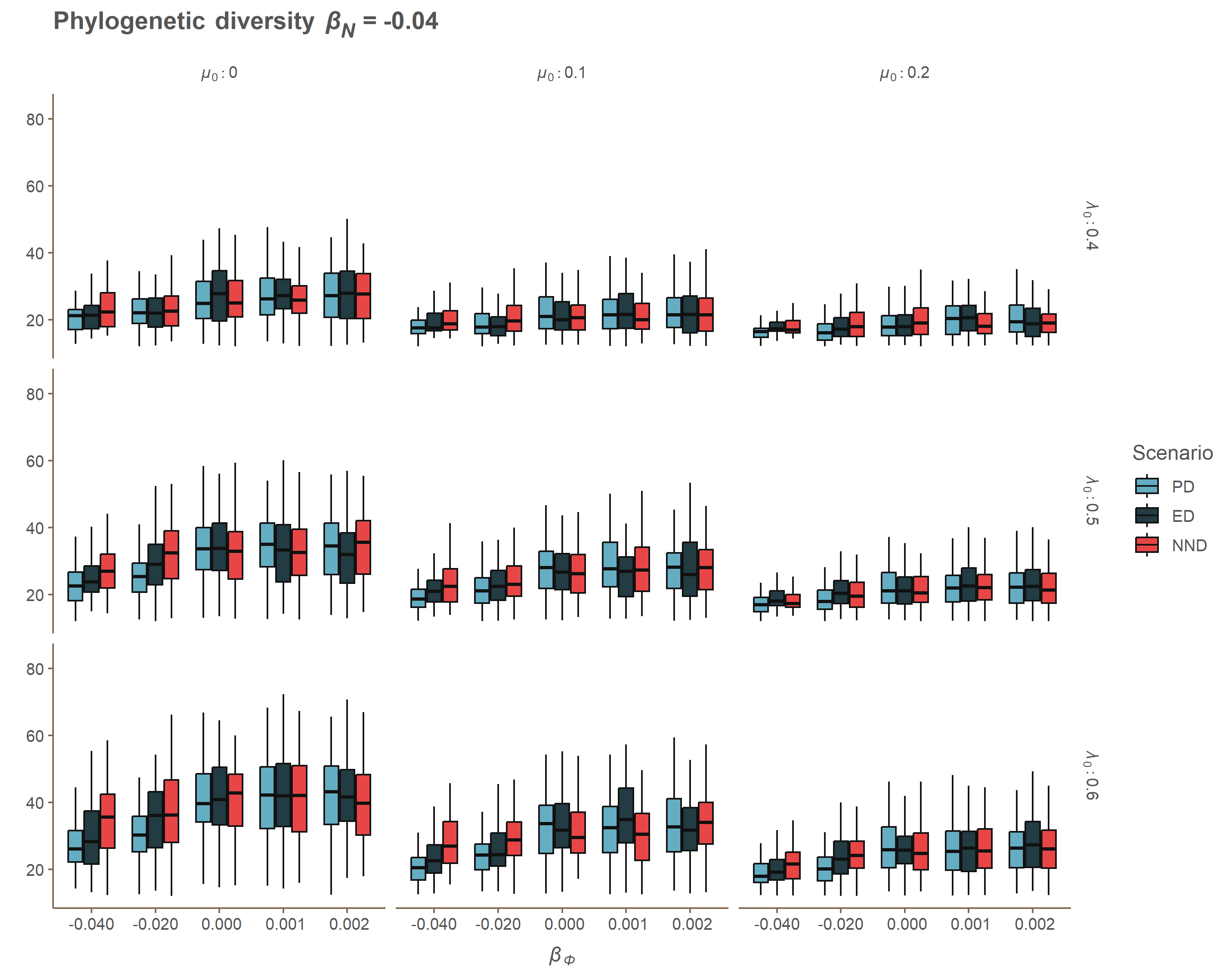

### Appendix A beta_n_SR2.gif

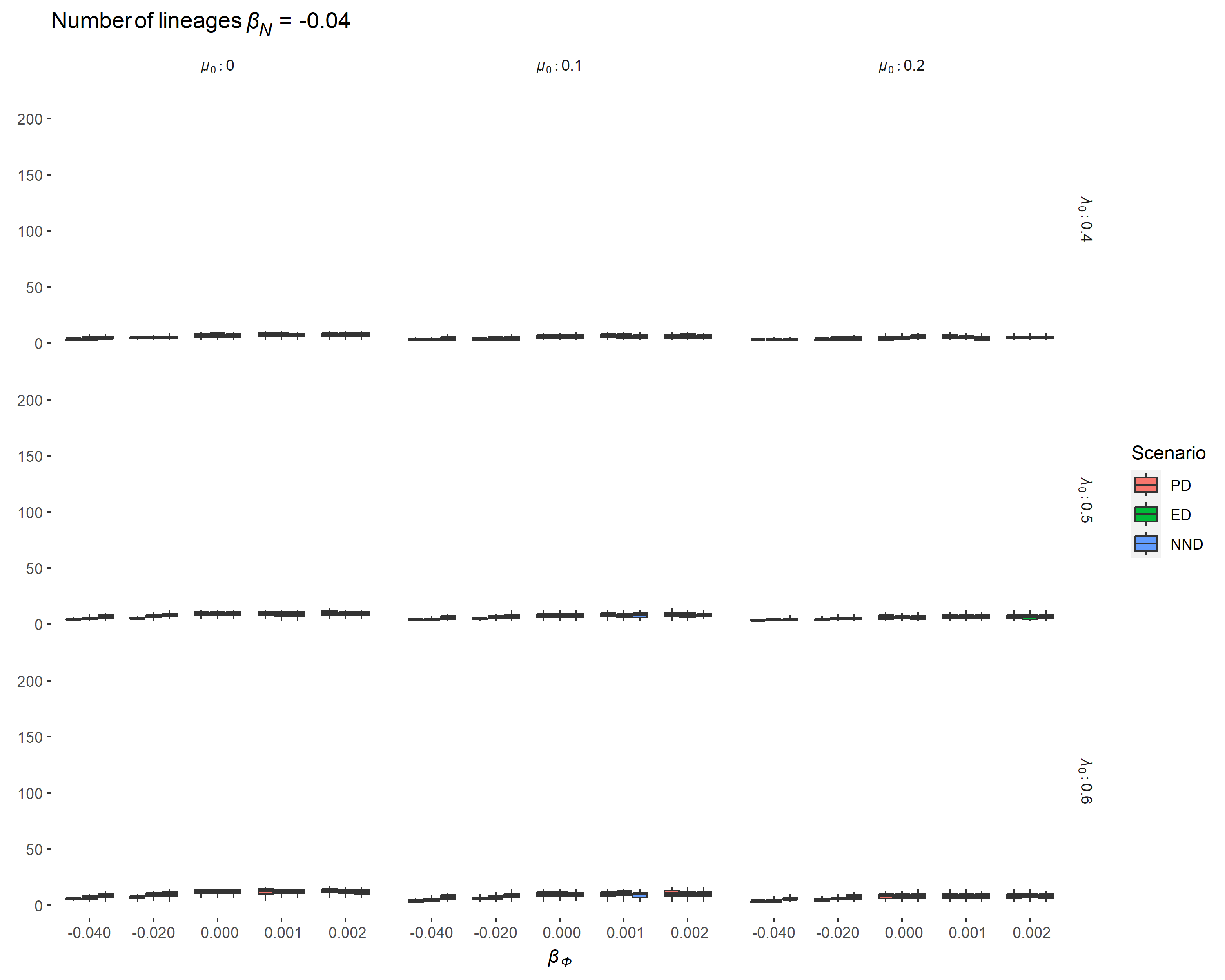

### Appendix A beta_n_SR.gif

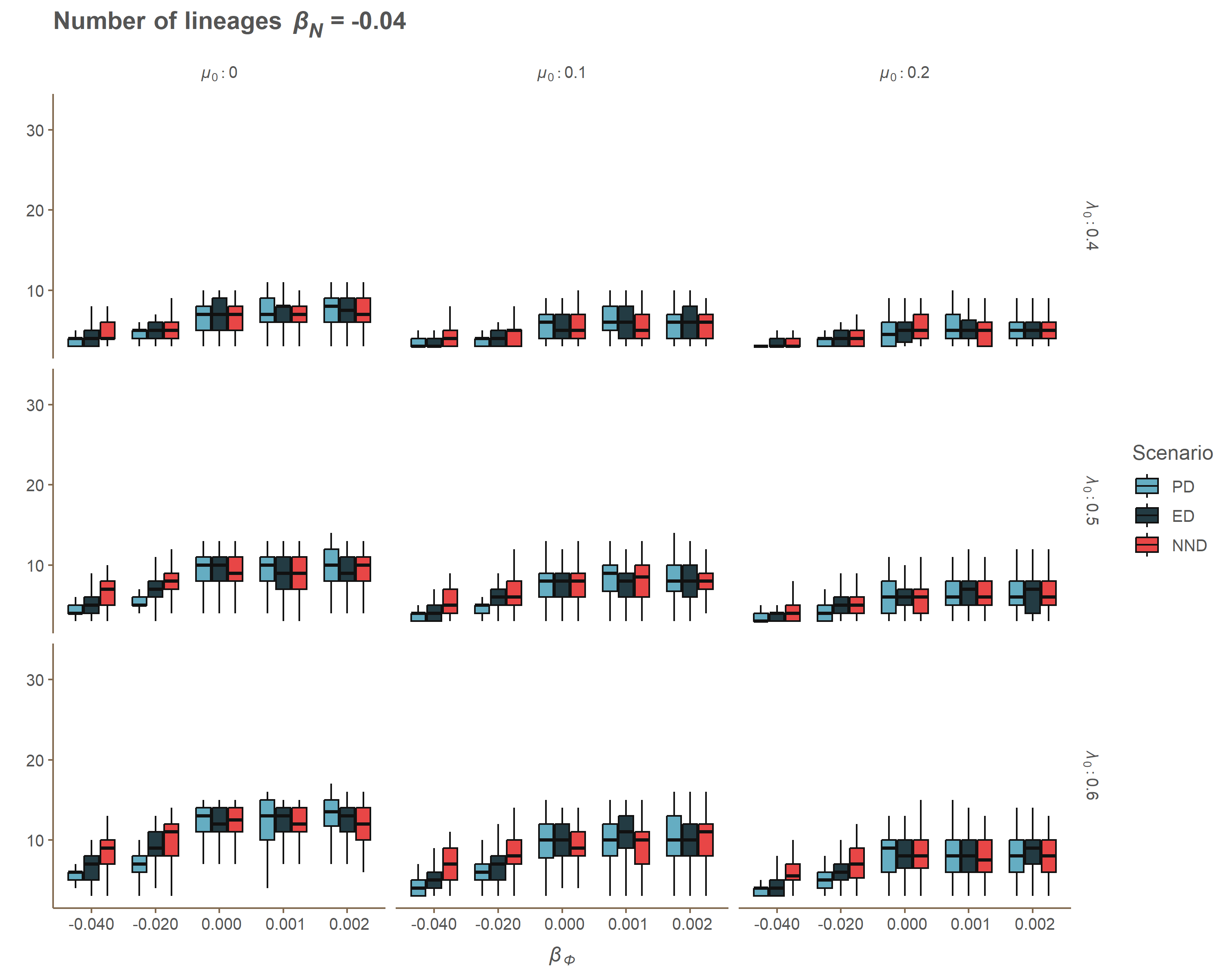

### Appendix A beta_phi_ERE.gif

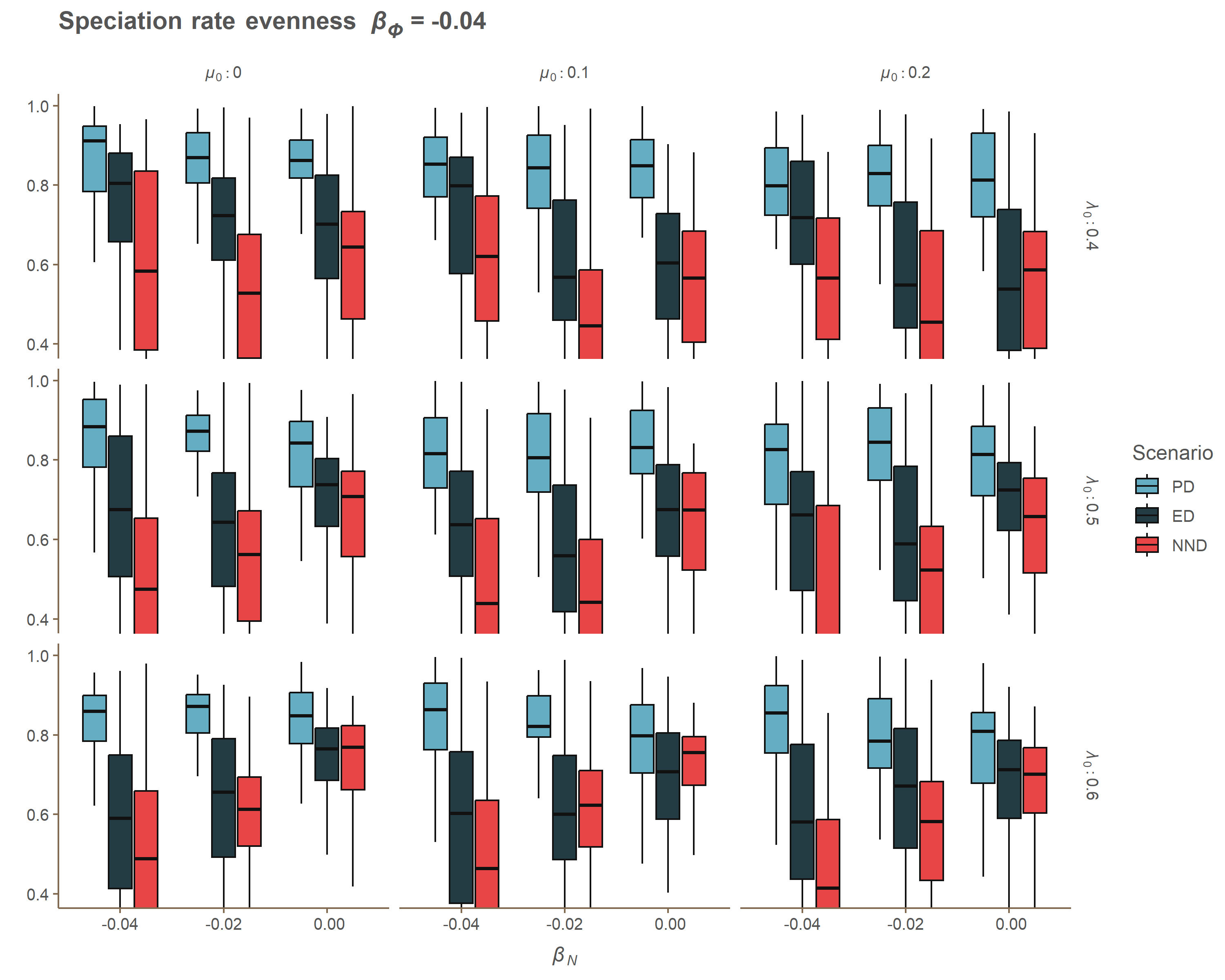

### Appendix A beta_phi_Gamma.gif

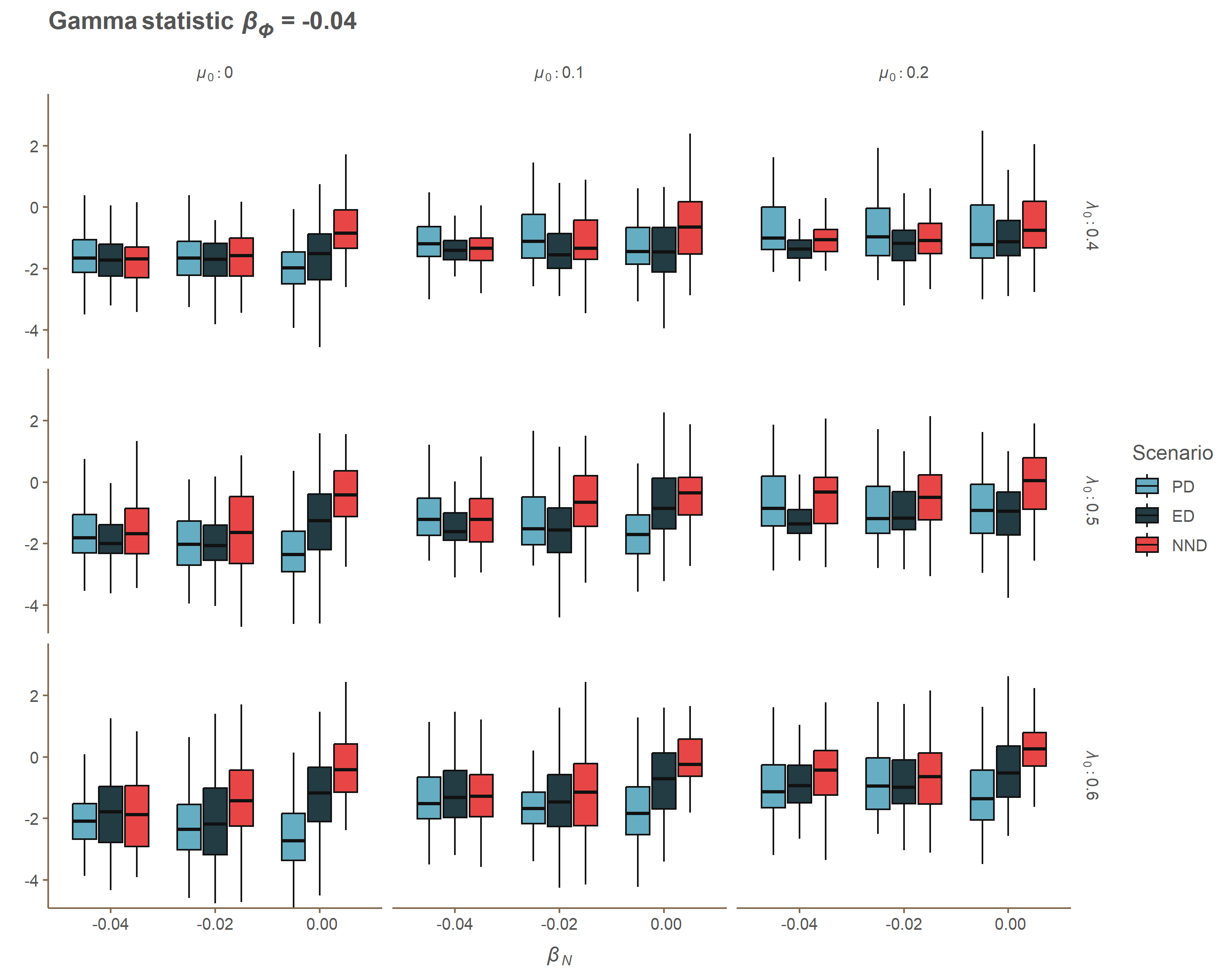

### Appendix A beta_phi_J_One.gif

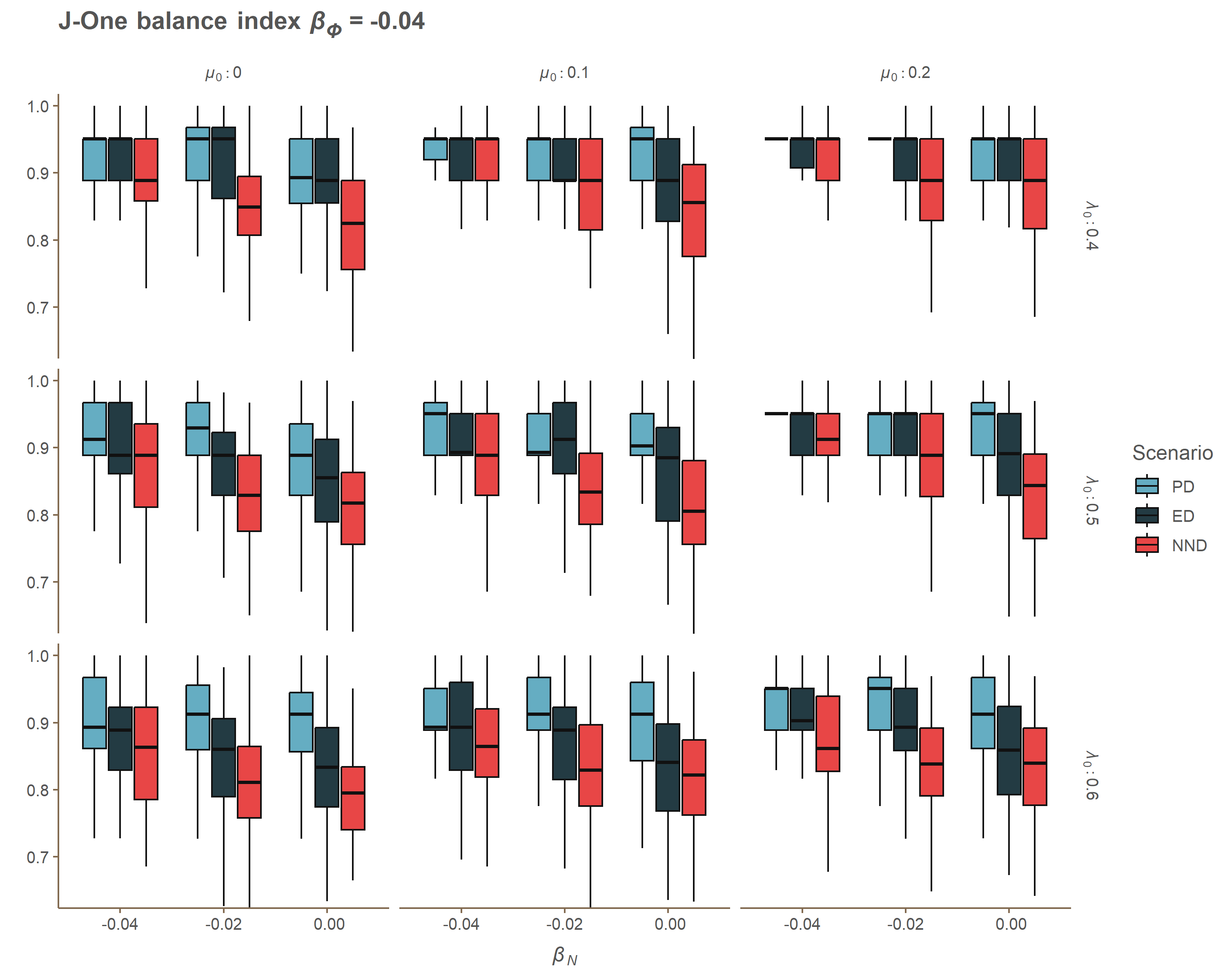

### Appendix A beta_phi_MBL.gif

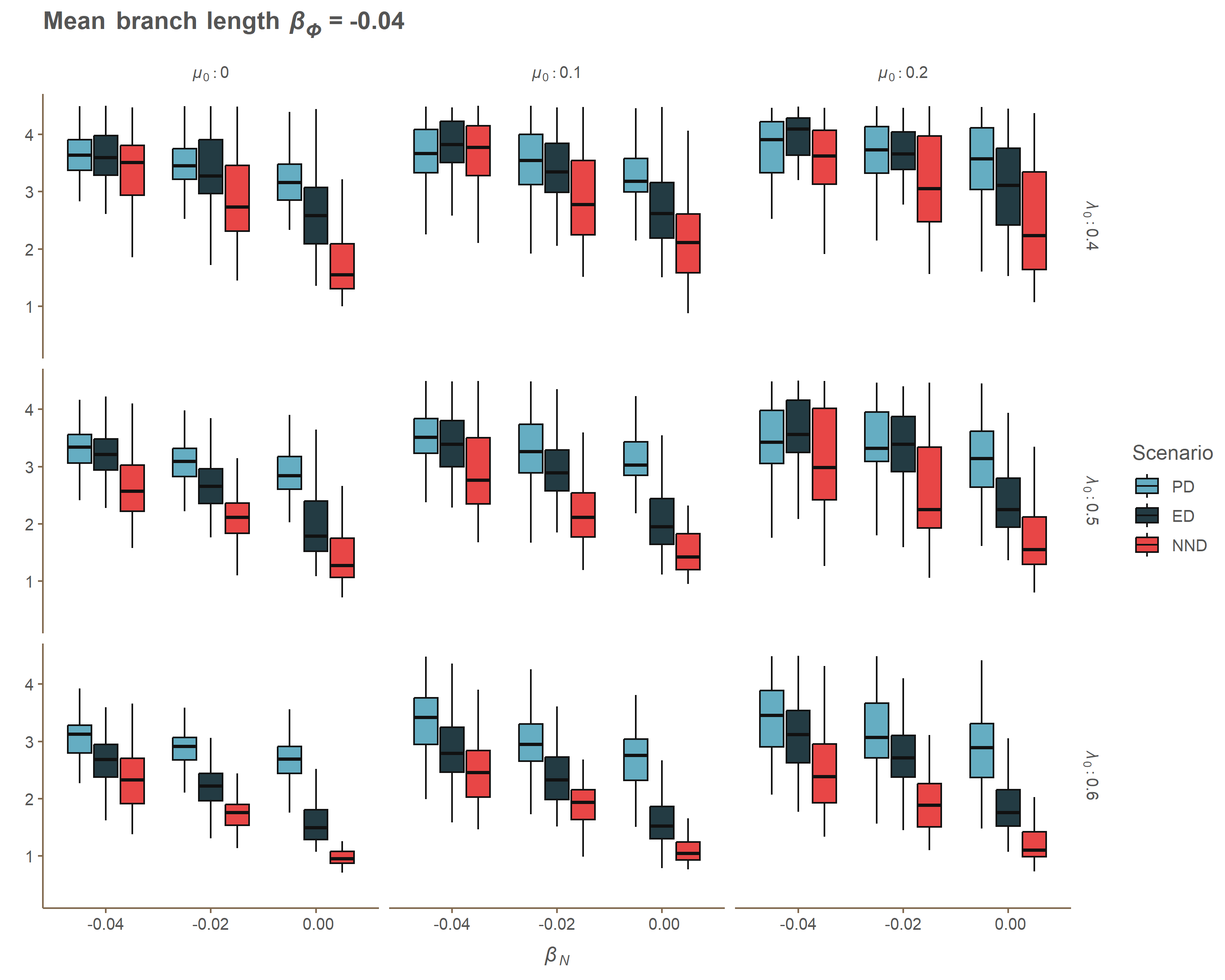

### Appendix A beta_phi_MNTD.gif

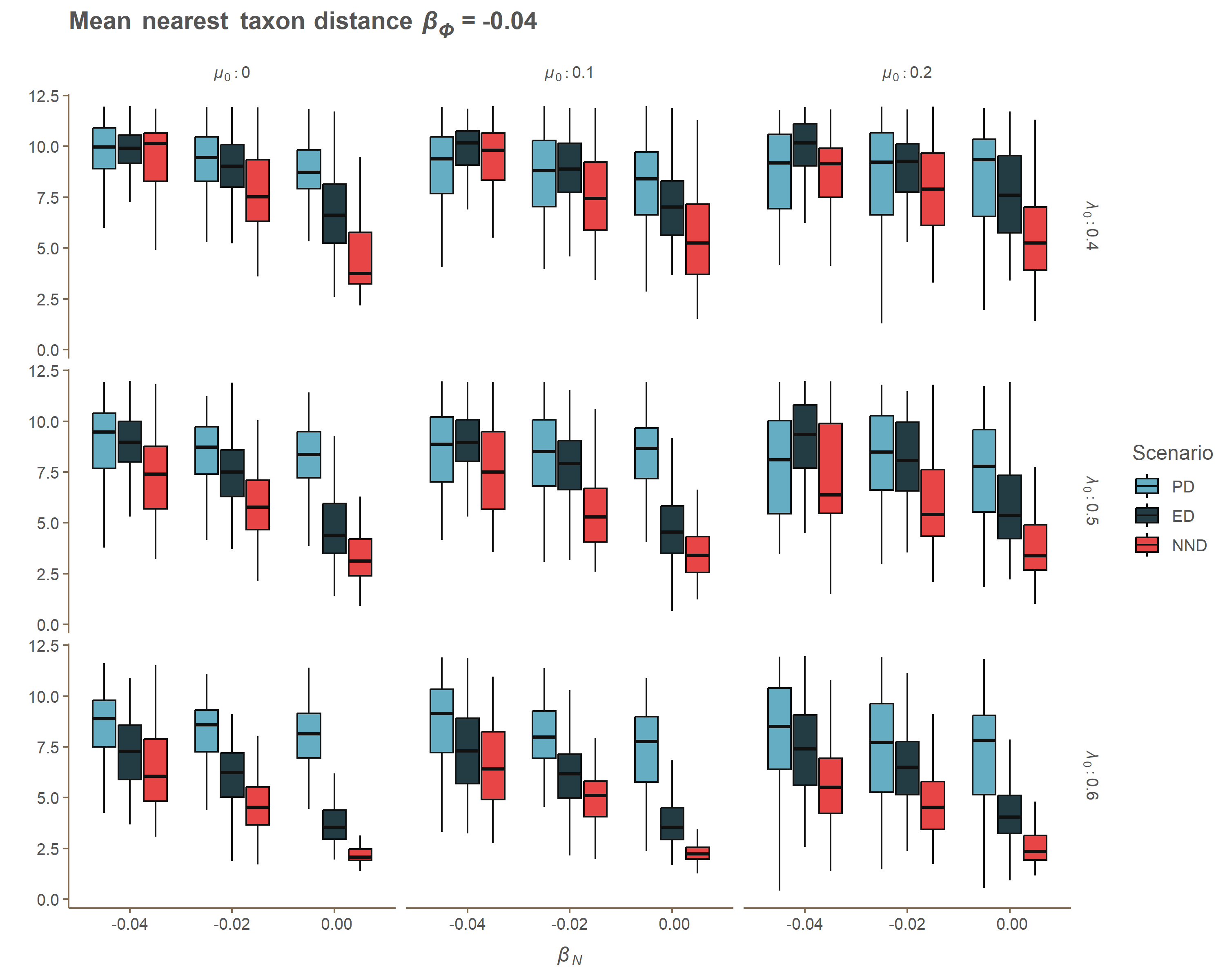

### Appendix A beta_phi_MPD.gif

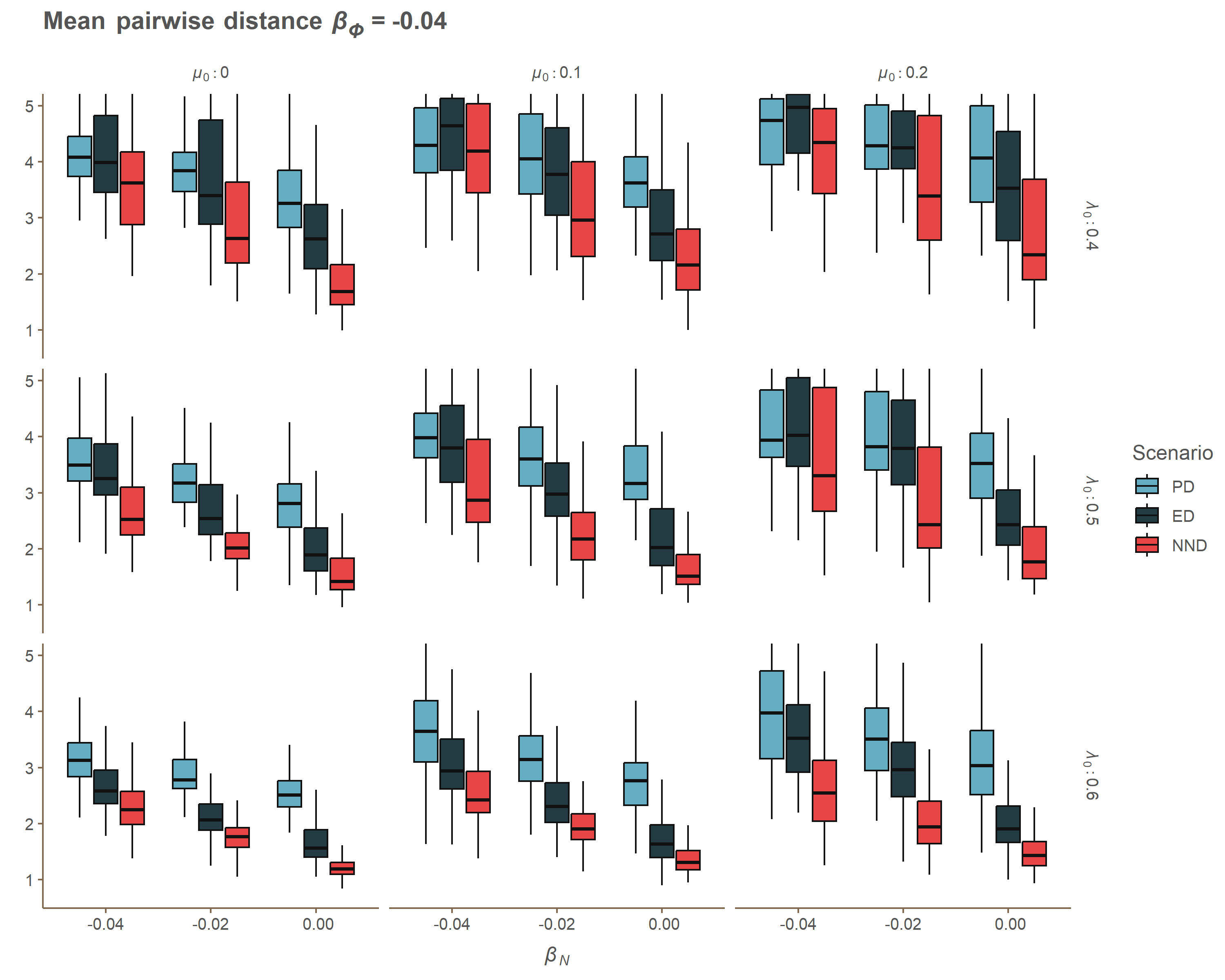

### Appendix A beta_phi_PD.gif

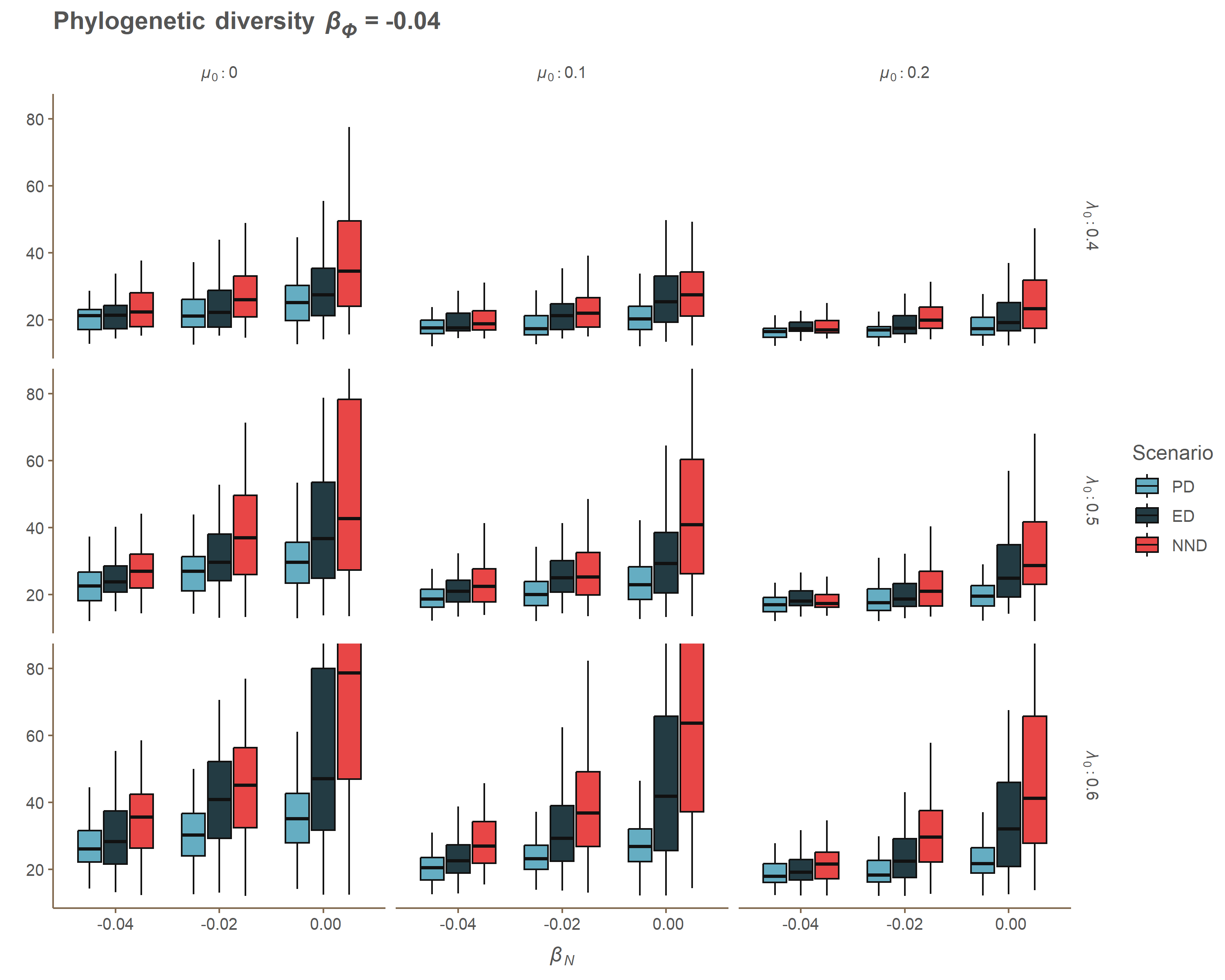

### Appendix A beta_phi_SR2.gif

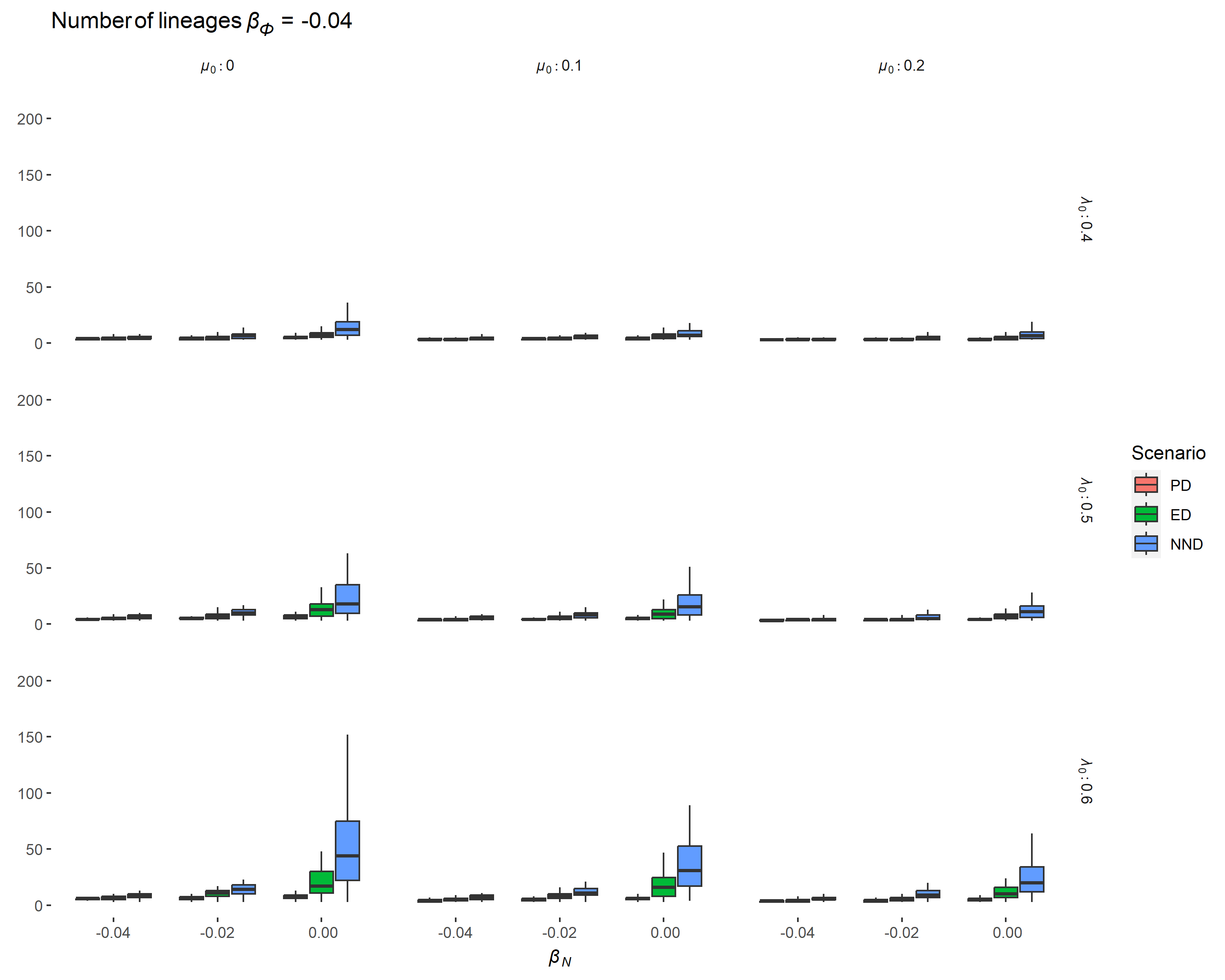

### Appendix A beta_phi_SR.gif

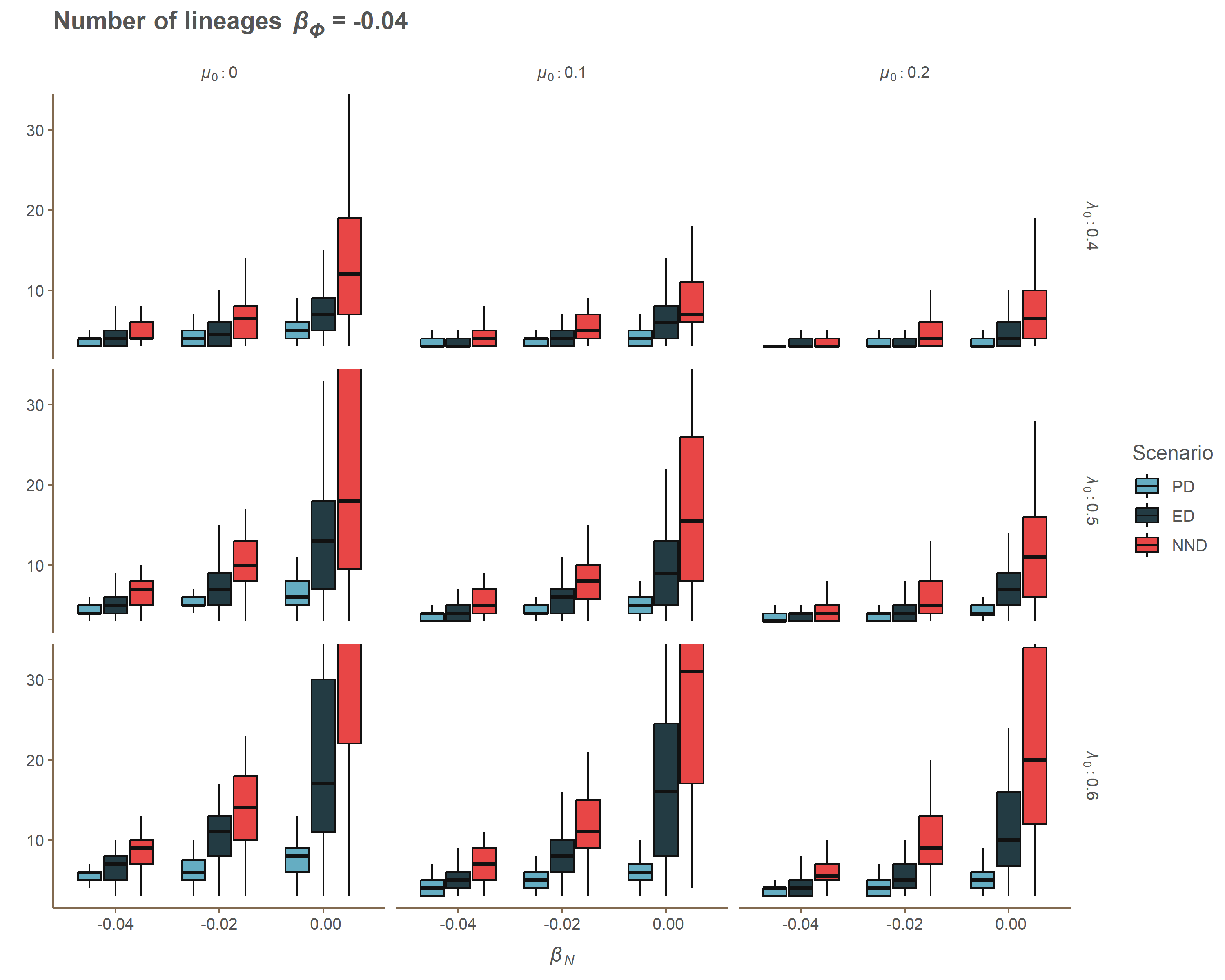

### Appendix A lambda_ERE.gif

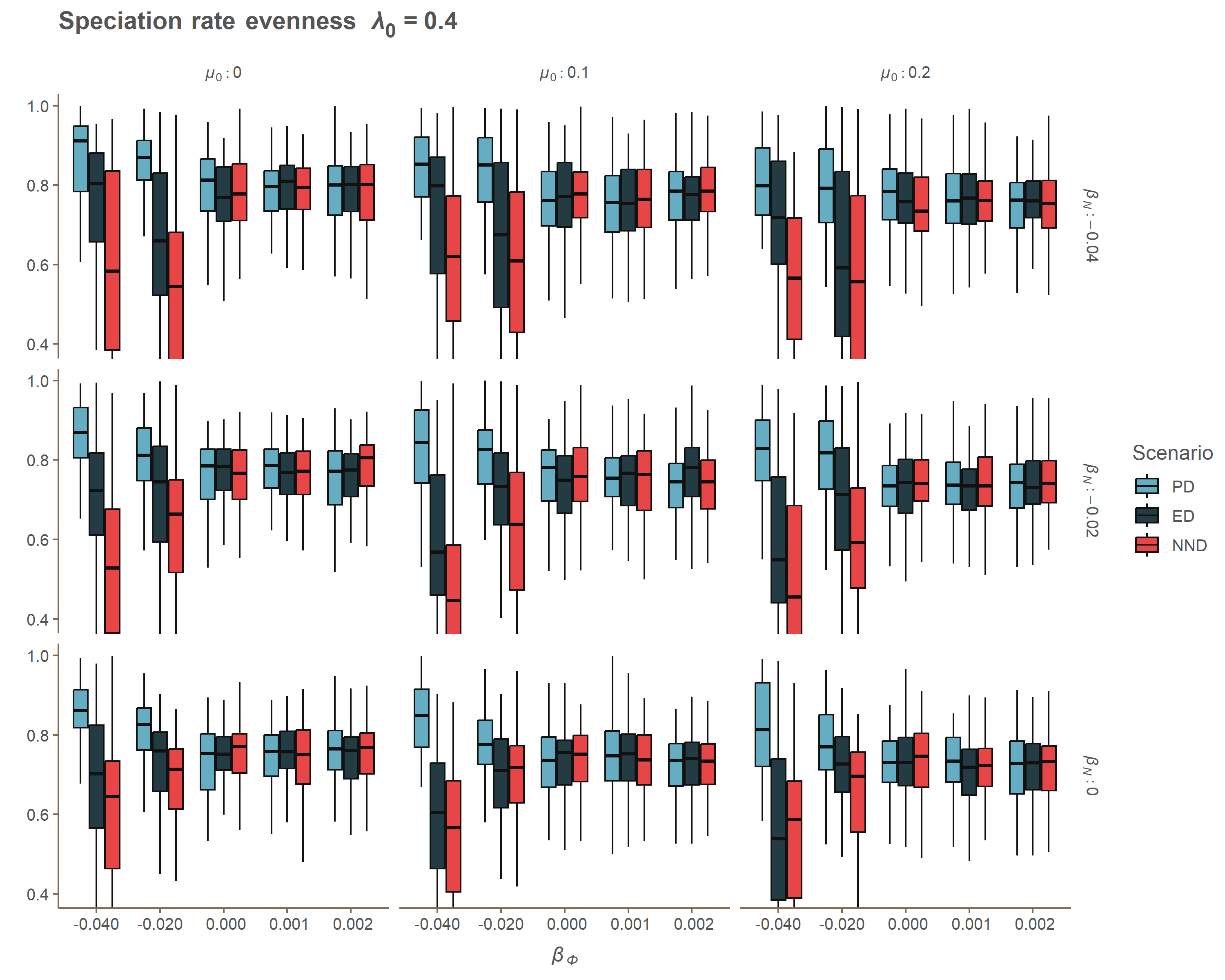

### Appendix A lambda_Gamma.gif

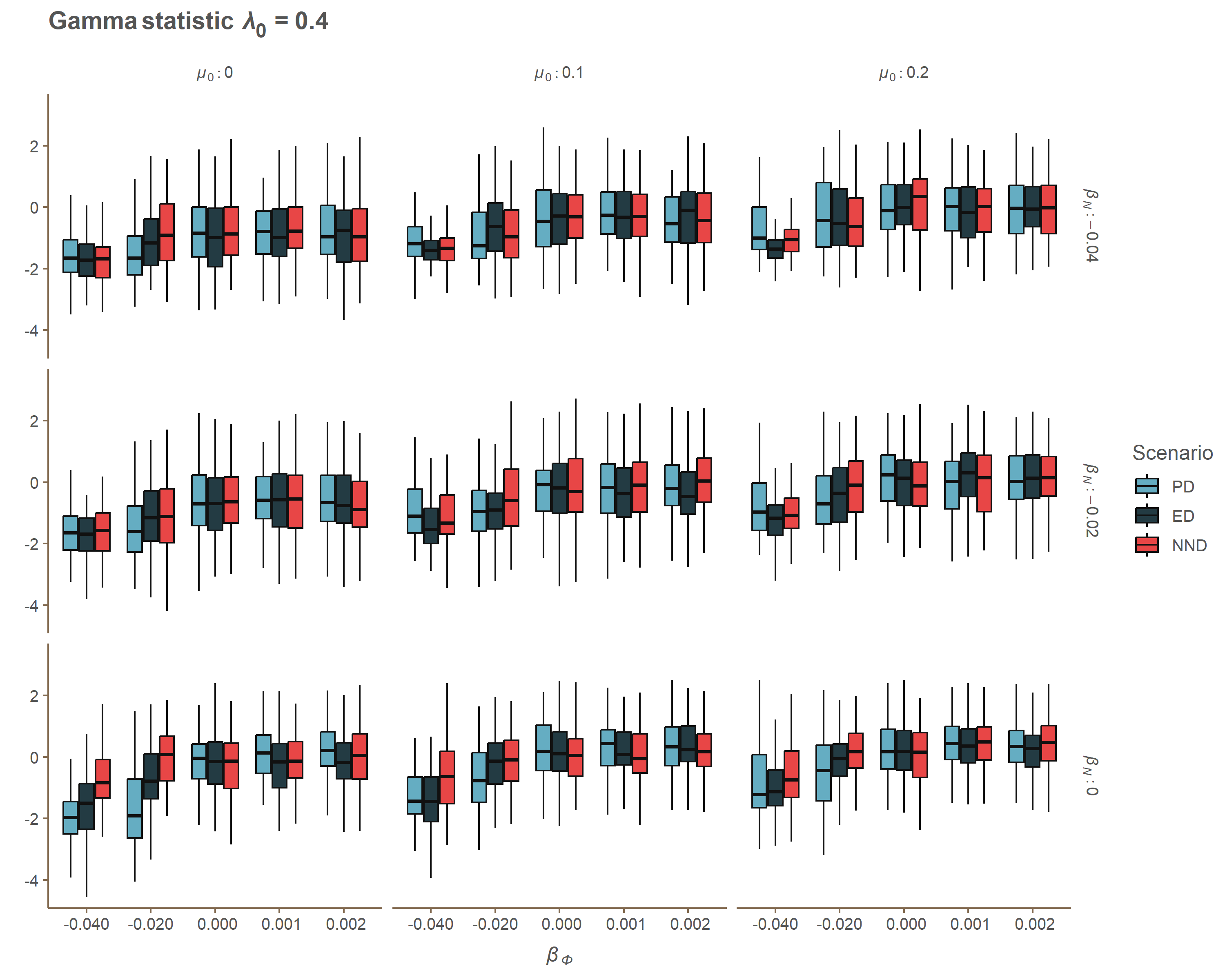

### Appendix A lambda_J_One.gif

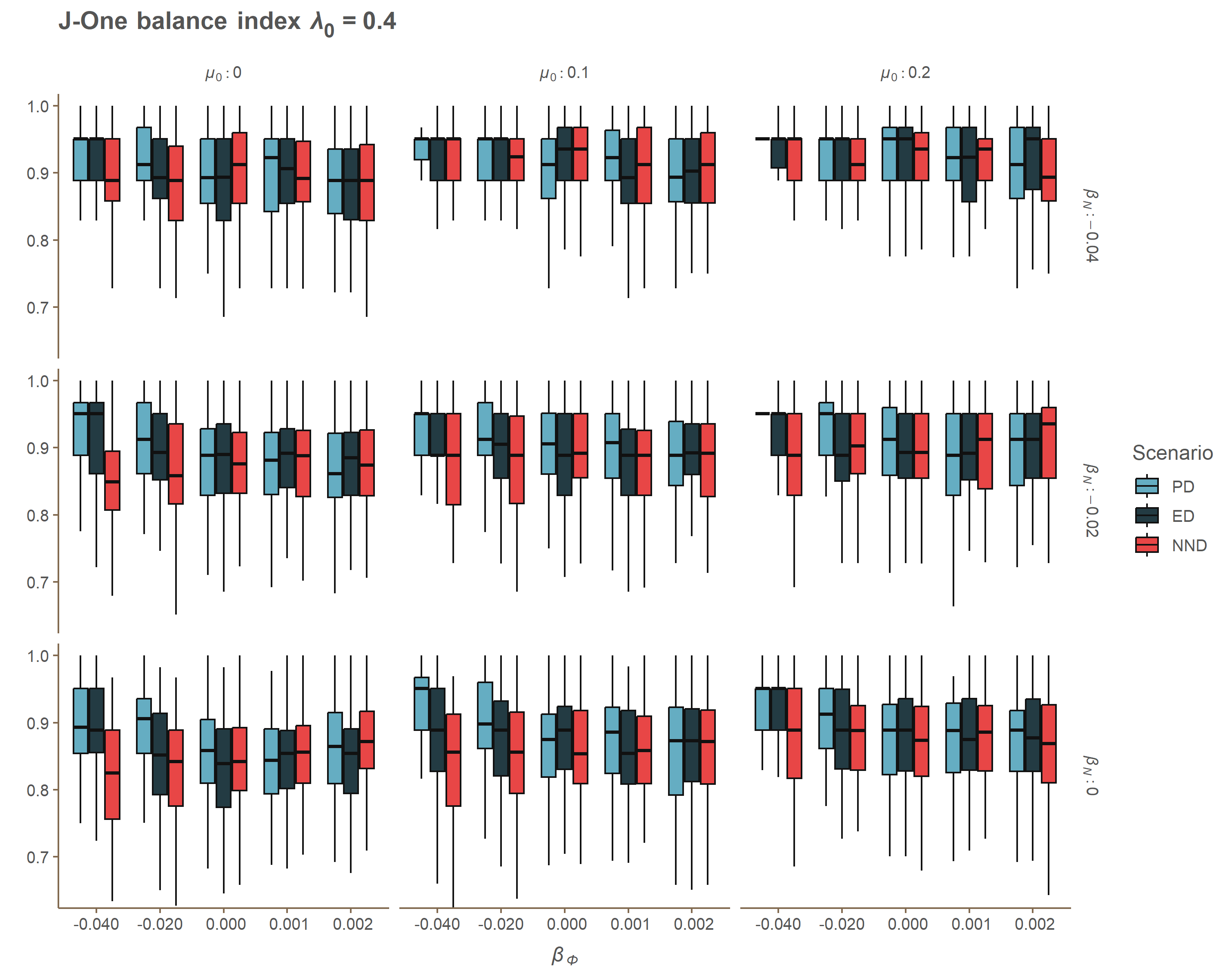

### Appendix A lambda_MBL.gif

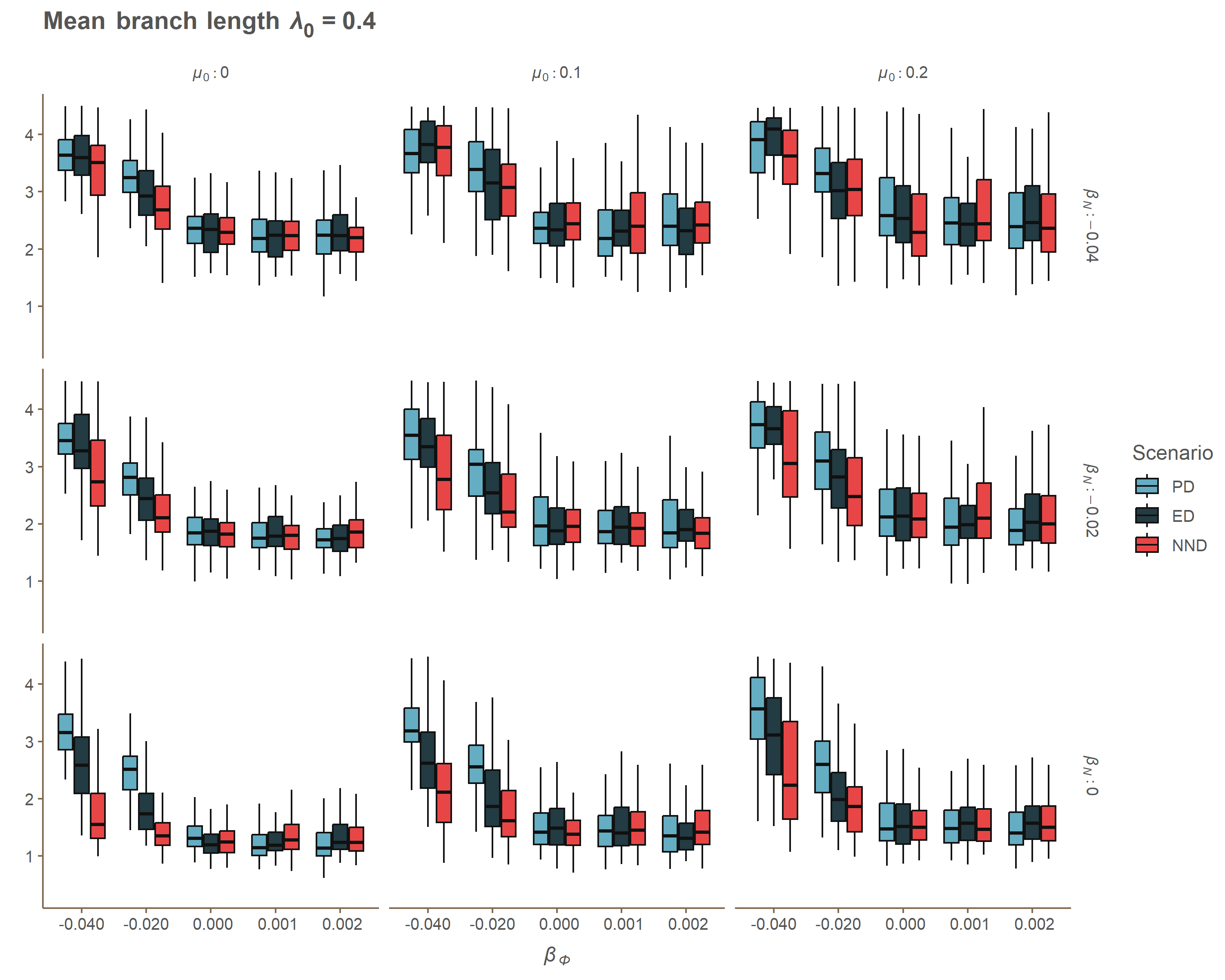

### Appendix A lambda_MNTD.gif

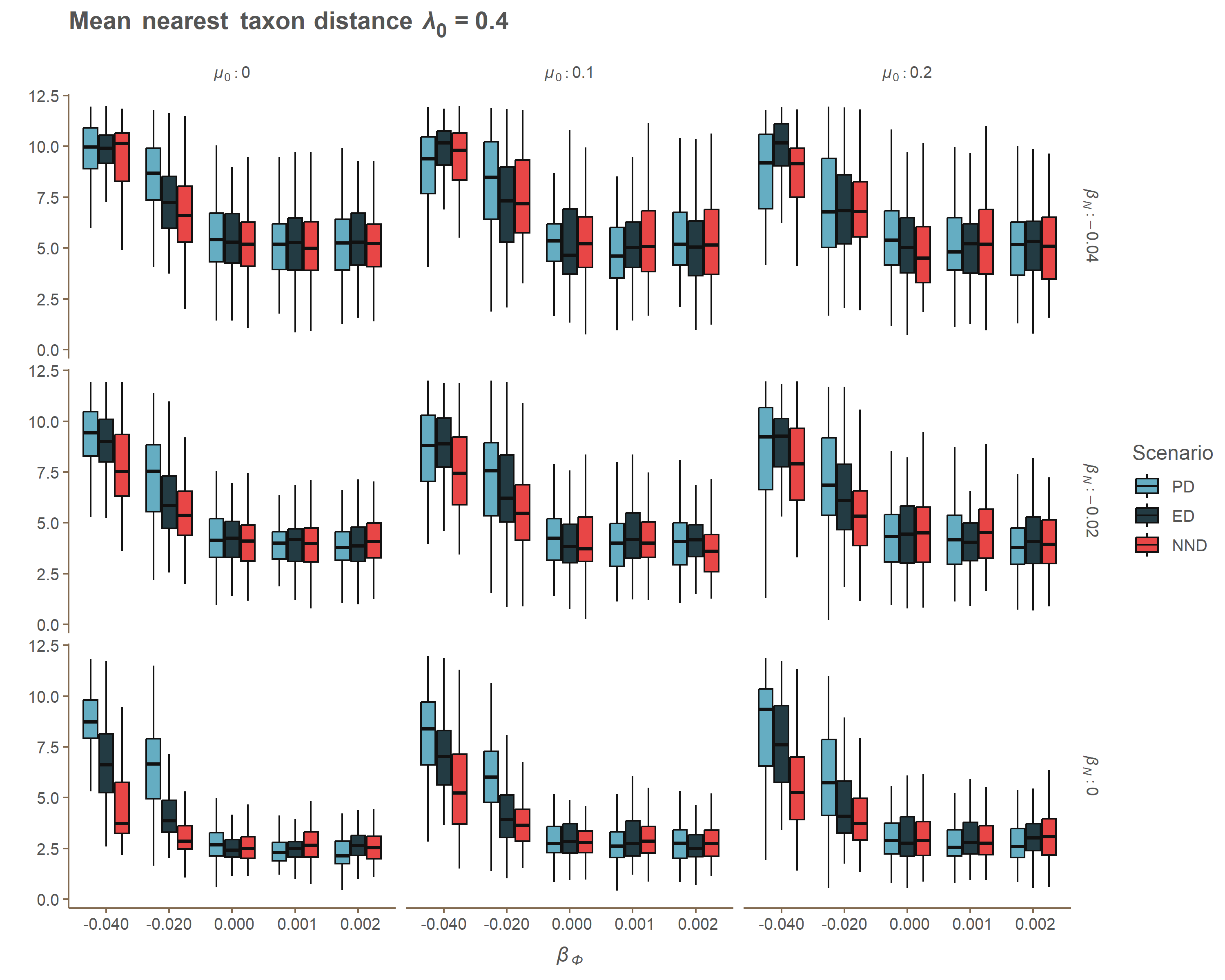

### Appendix A lambda_MPD.gif

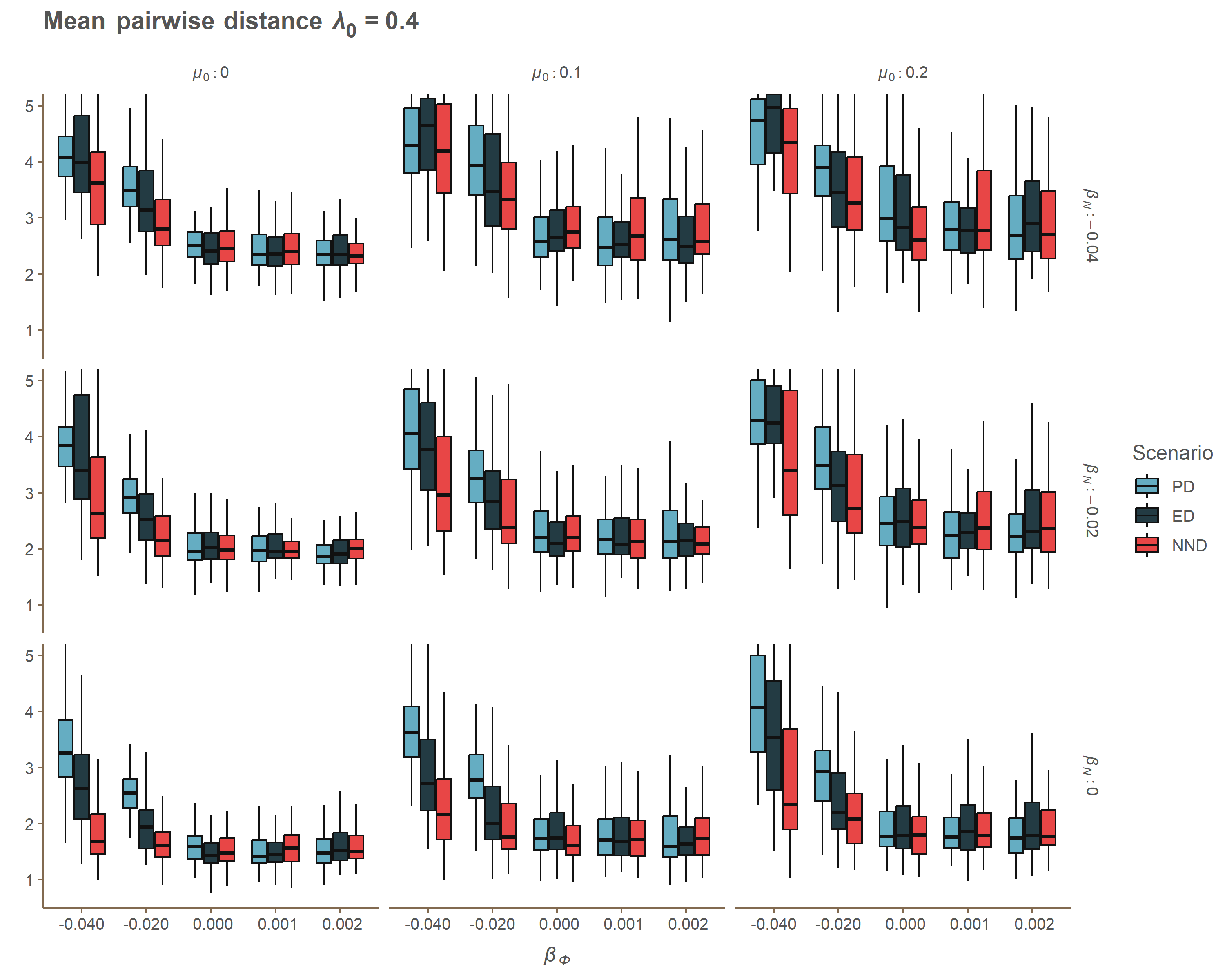

### Appendix A lambda_PD.gif

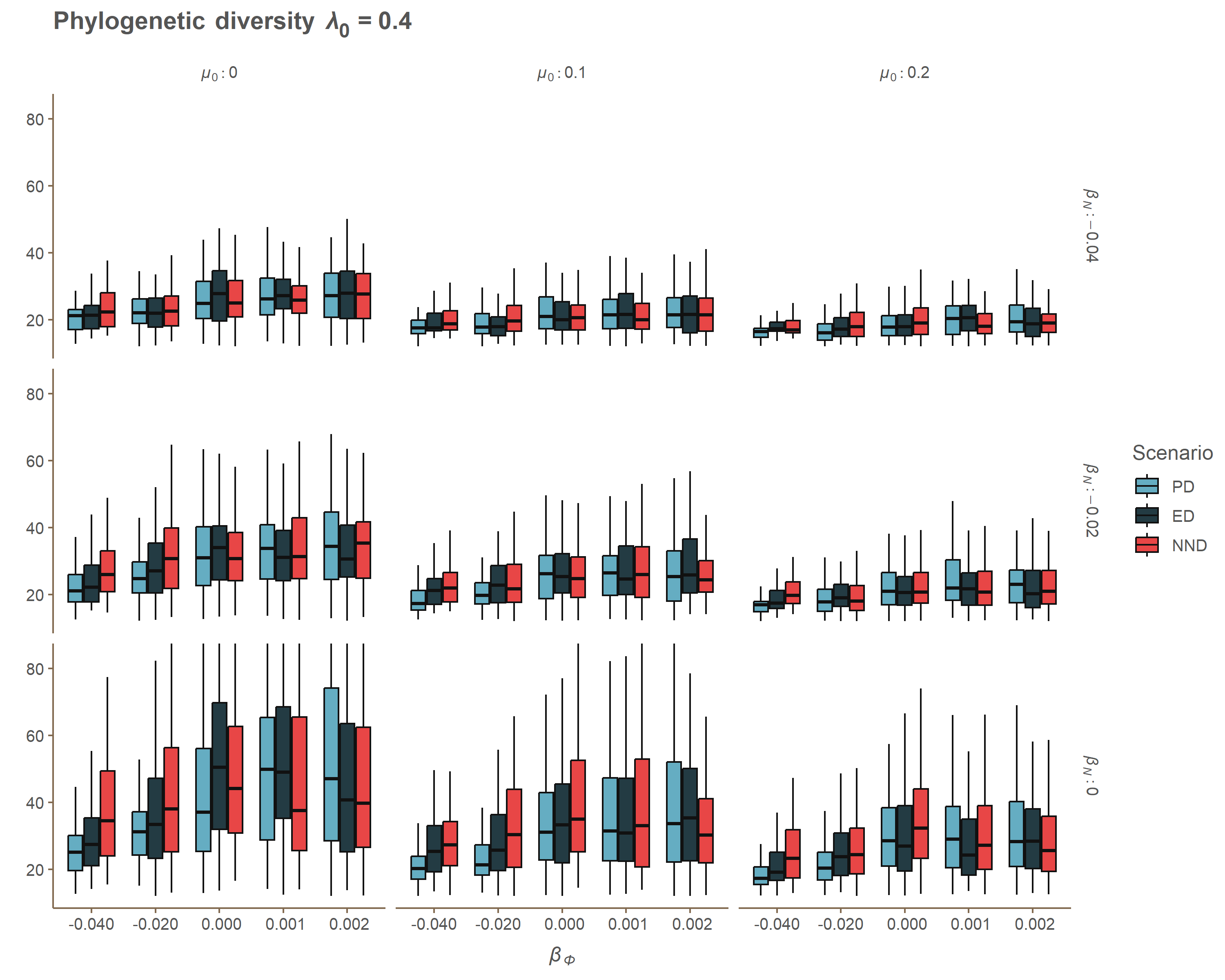

### Appendix A lambda_SR2.gif

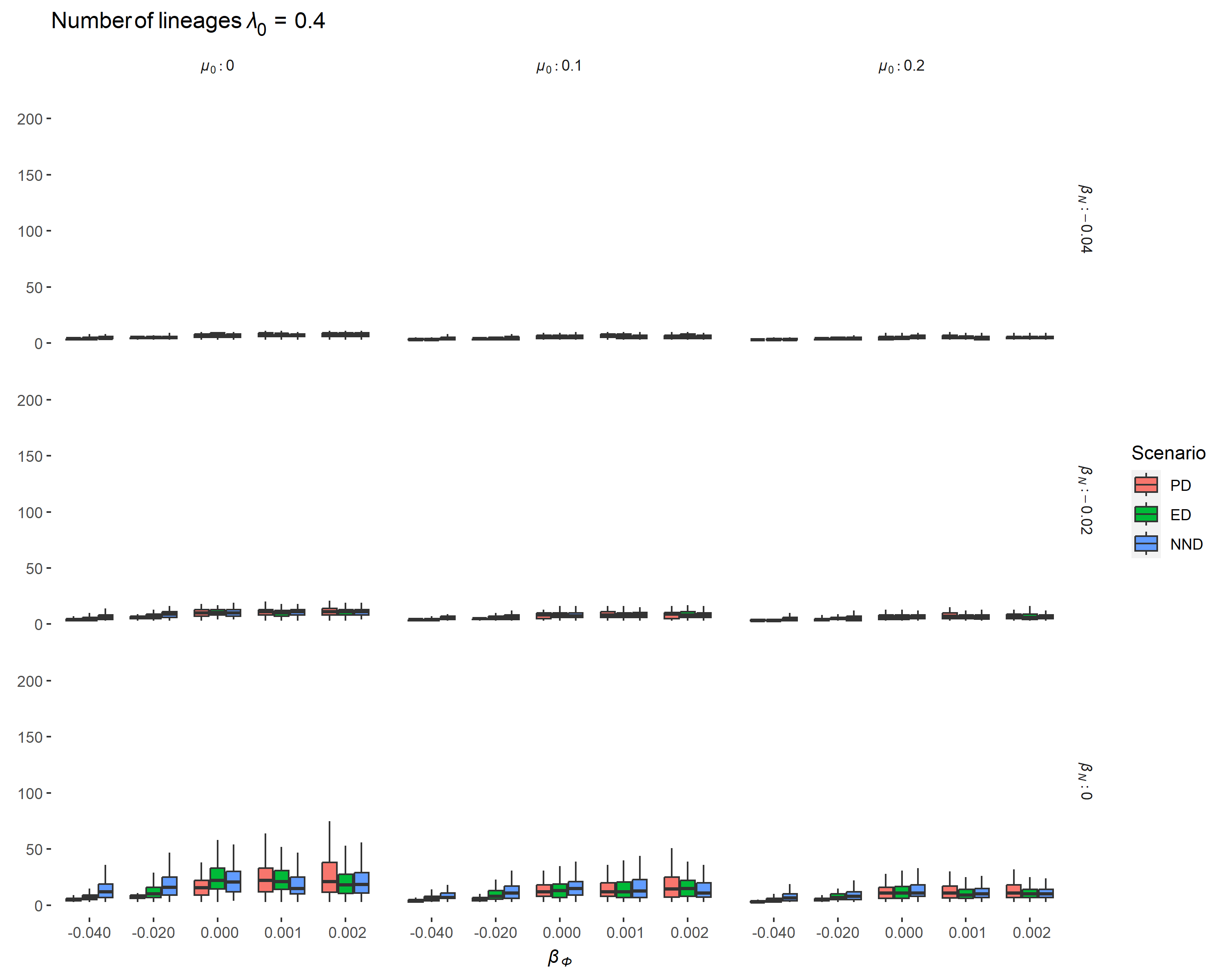

### Appendix A lambda_SR.gif

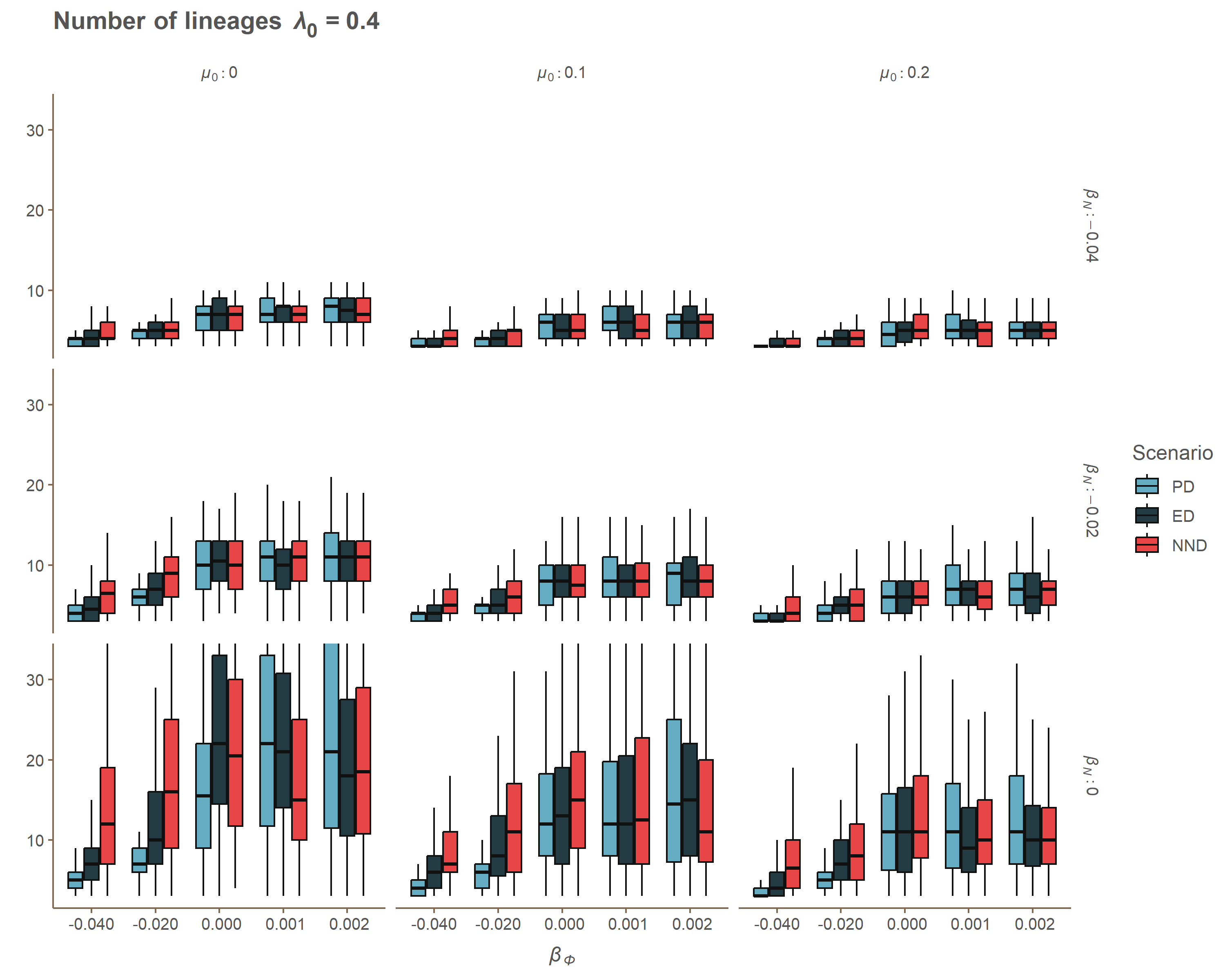

### Appendix A mu_ERE.gif

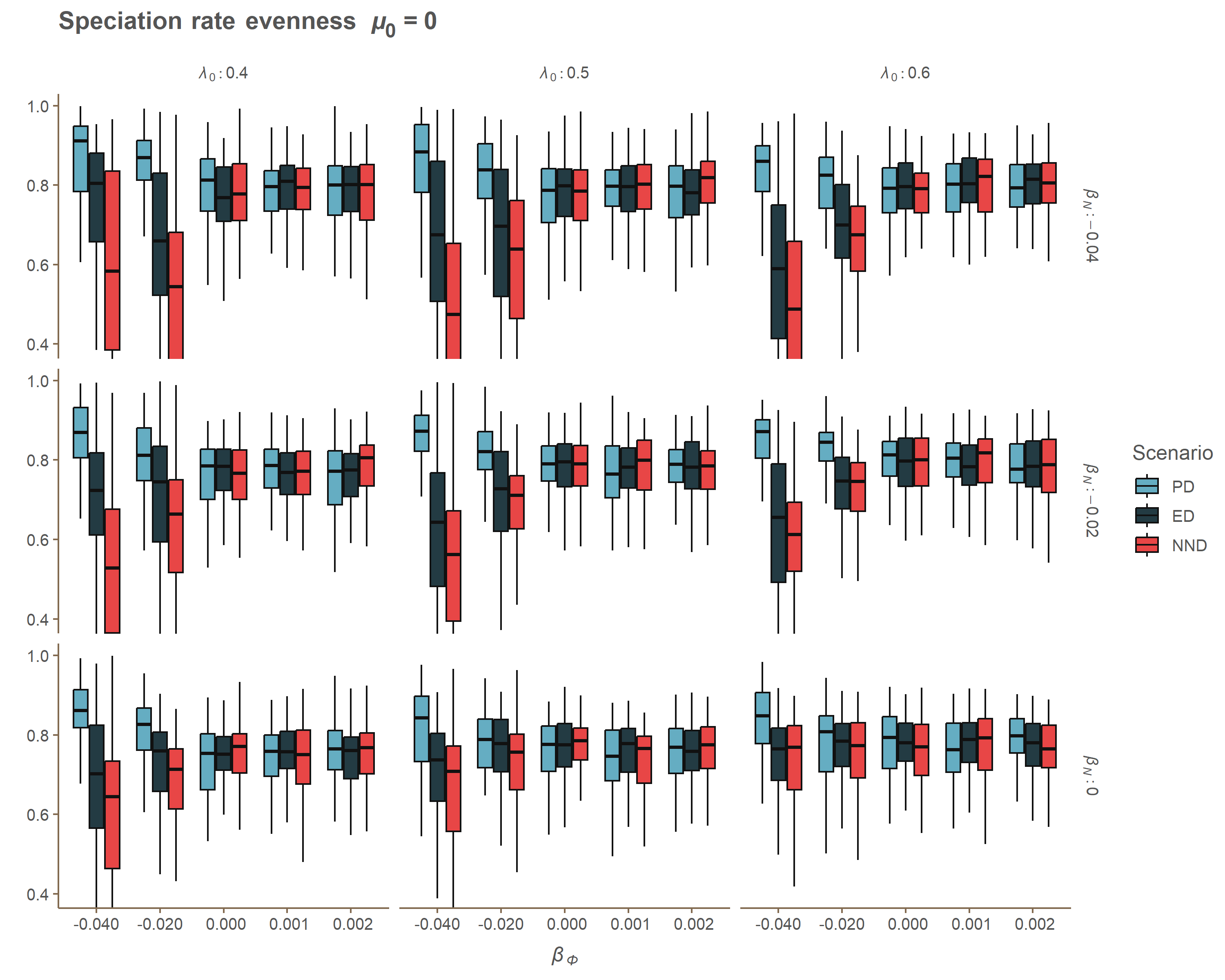

### Appendix A mu_Gamma.gif

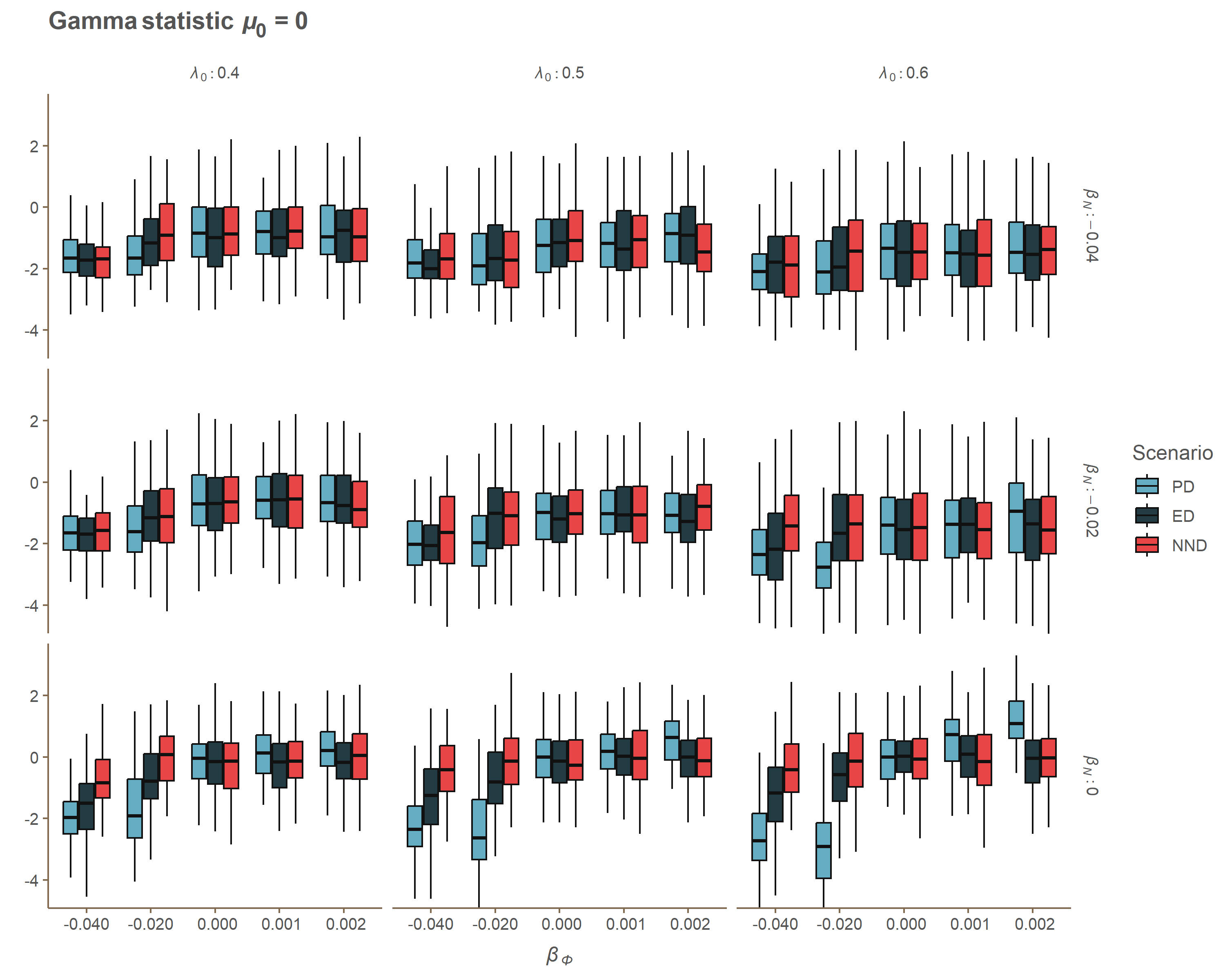

### Appendix A mu_J_One.gif

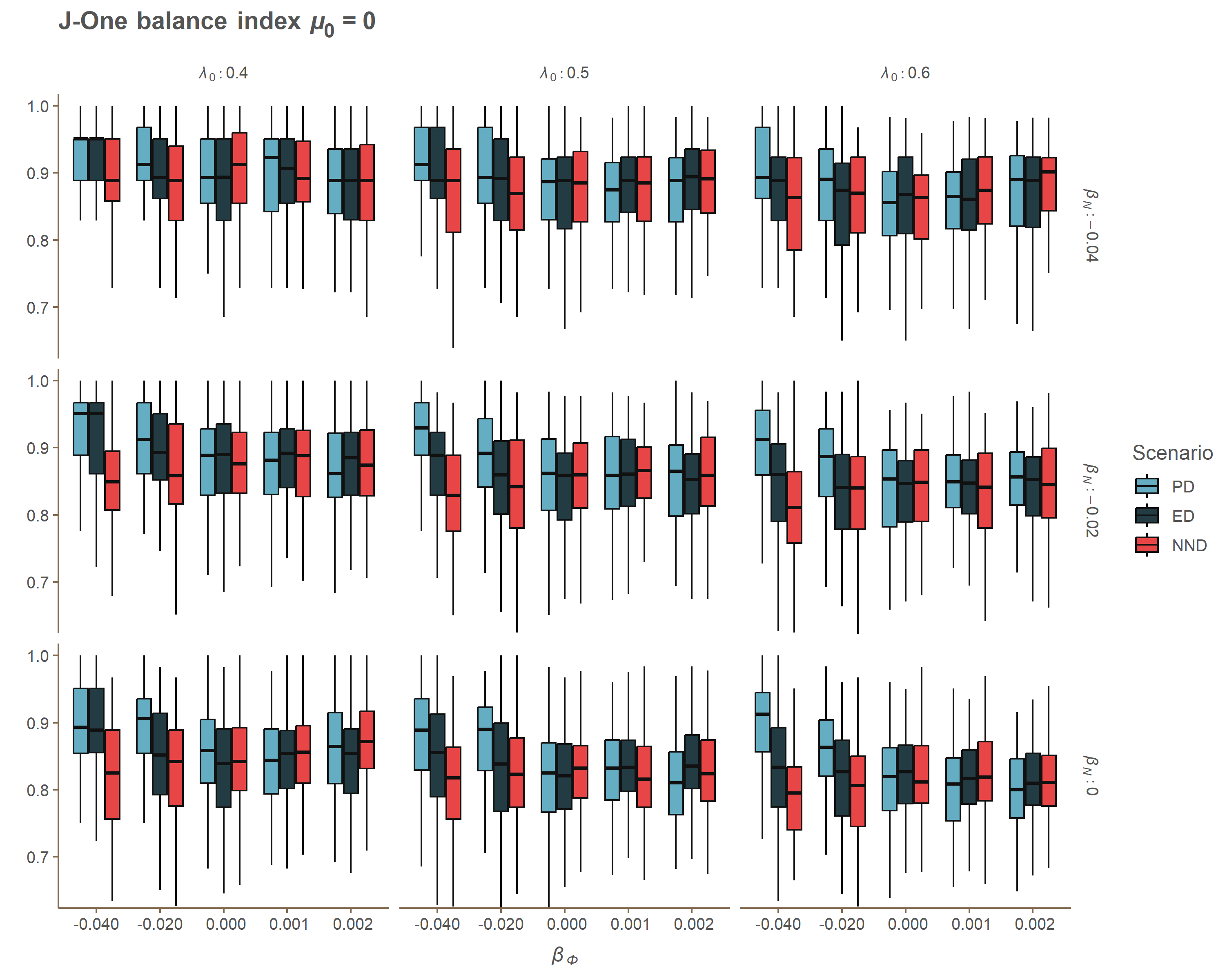

### Appendix A mu_MBL.gif

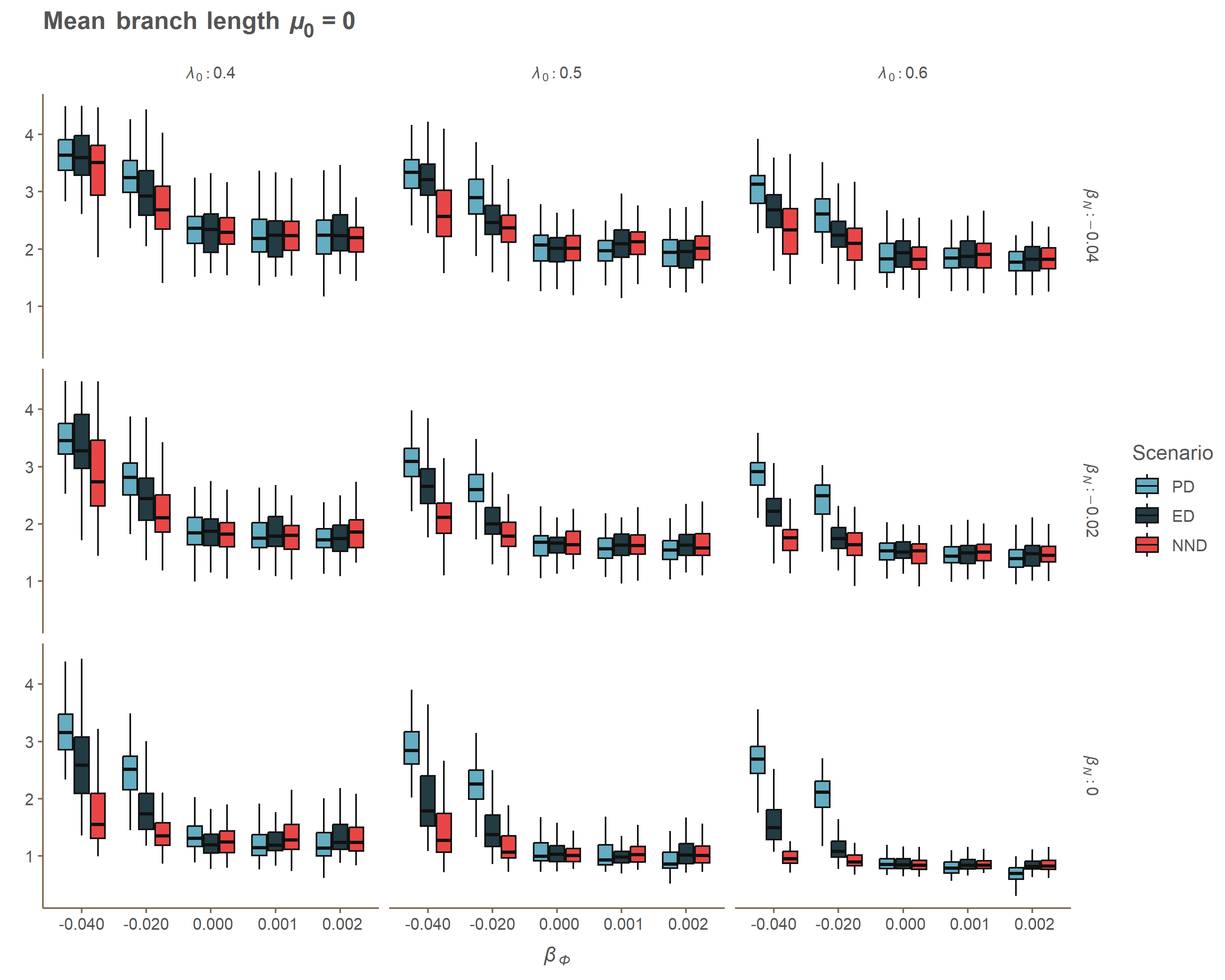

### Appendix A mu_MNTD.gif

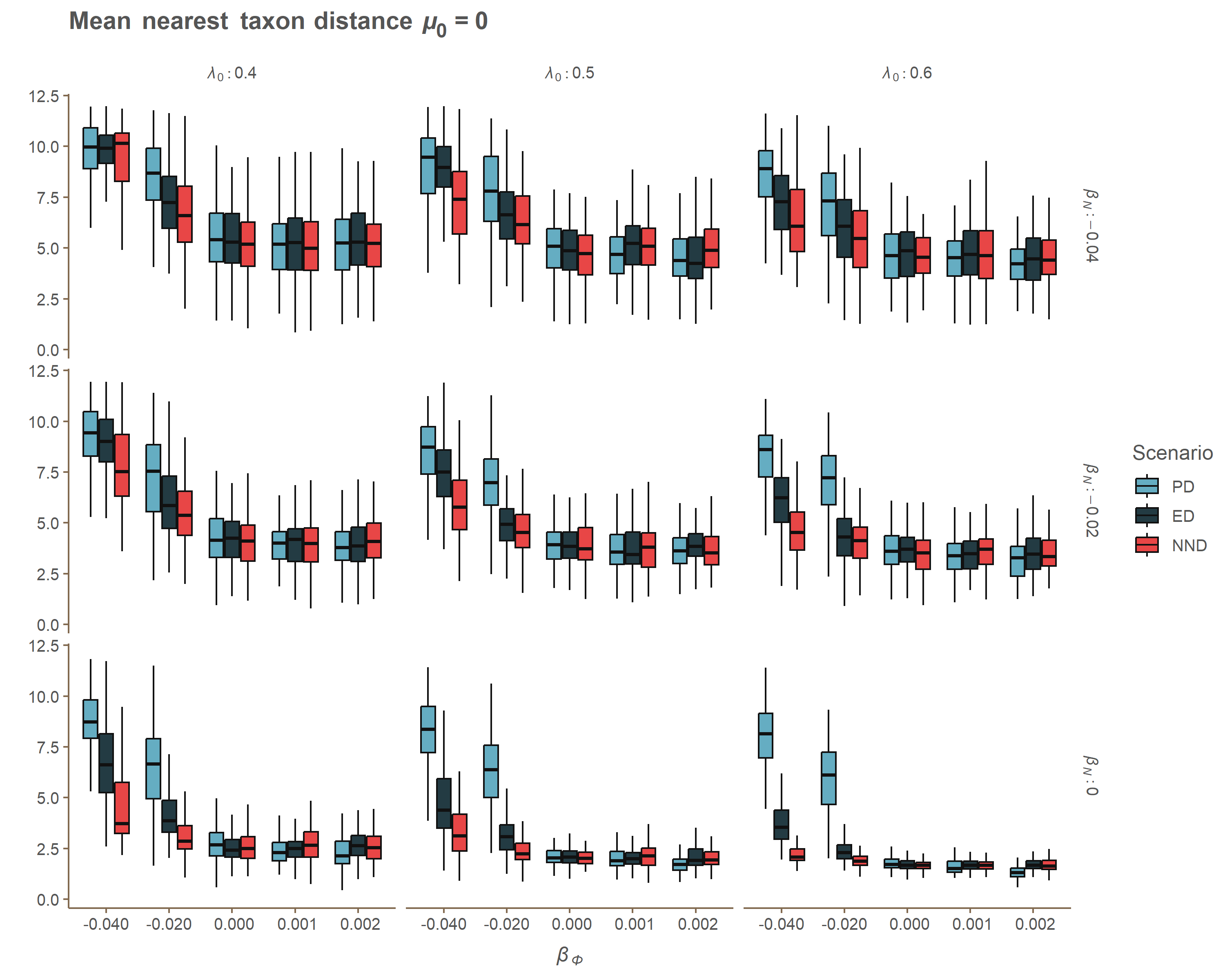

### Appendix A mu_MPD.gif

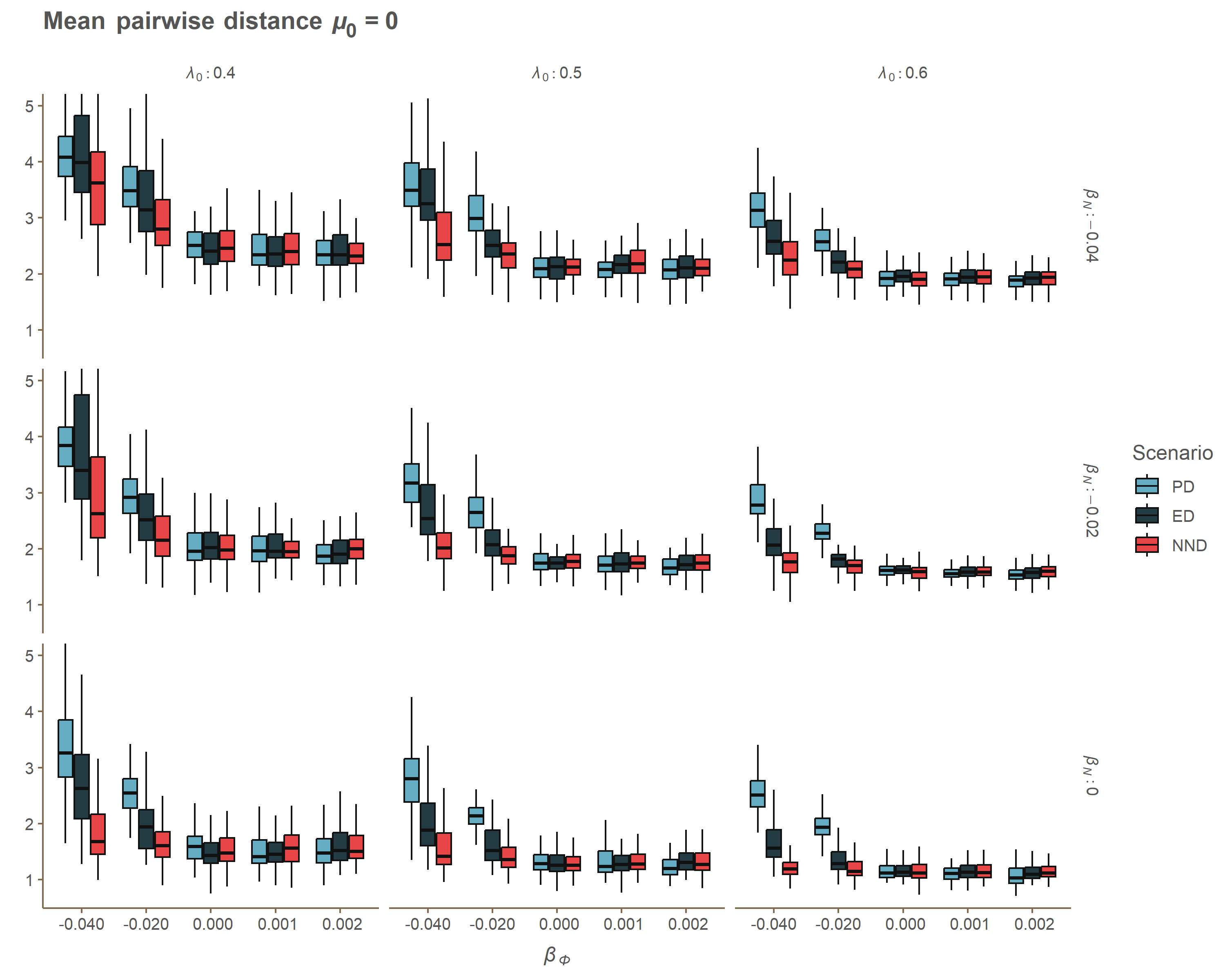

### Appendix A mu_PD.gif

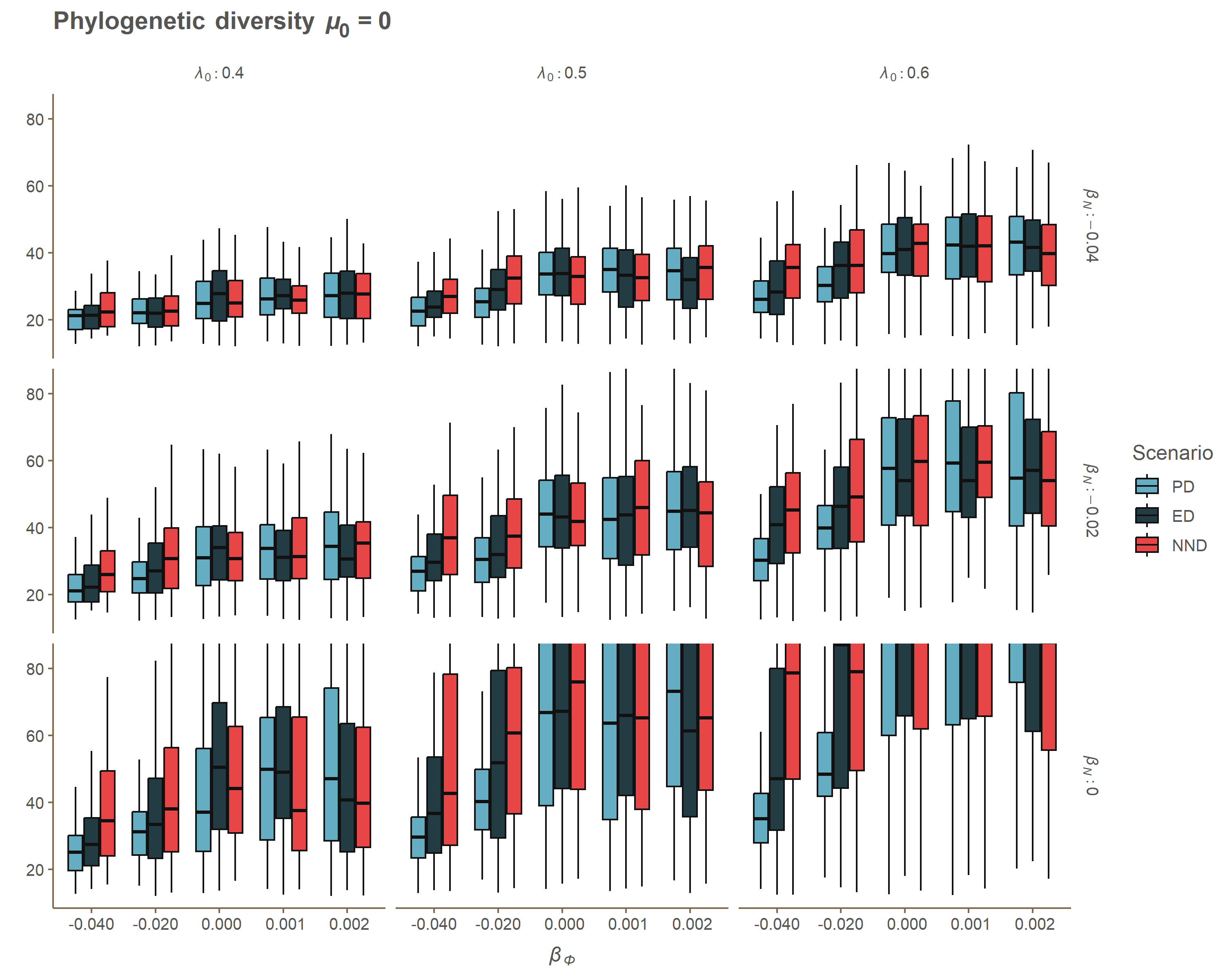

### Appendix A mu_SR2.gif

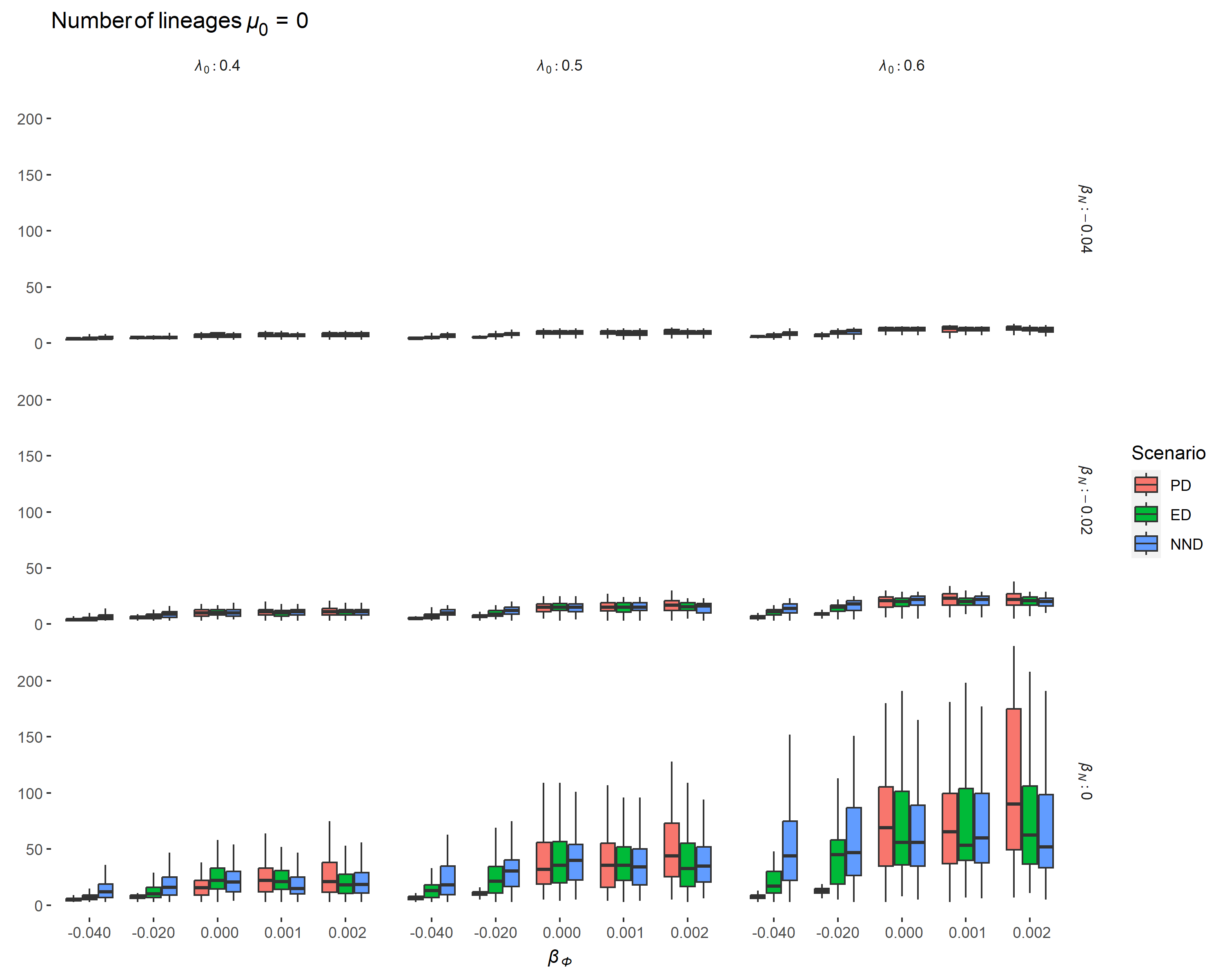

### Appendix A mu_SR.gif

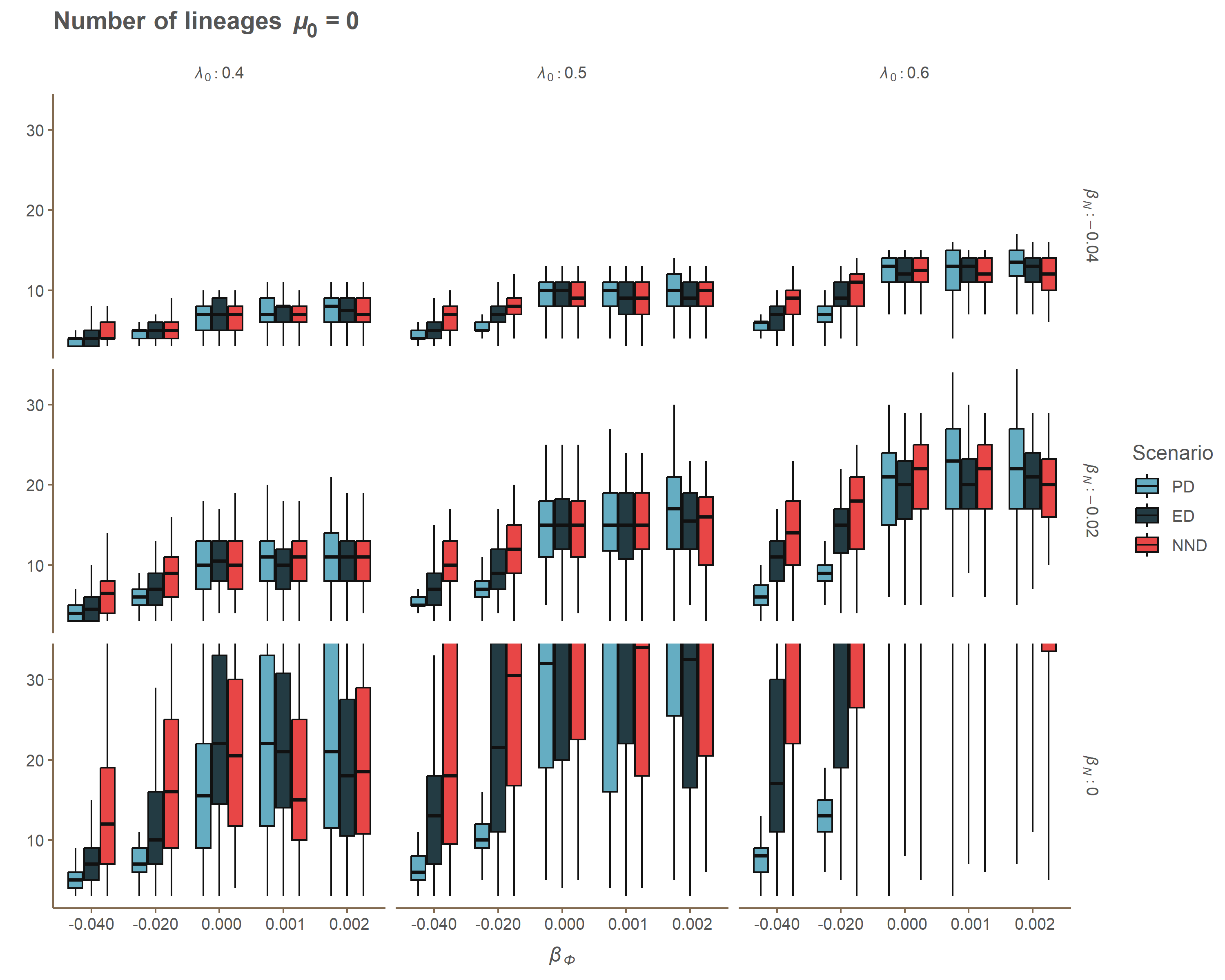

### Appendix B heatmap_ED.png

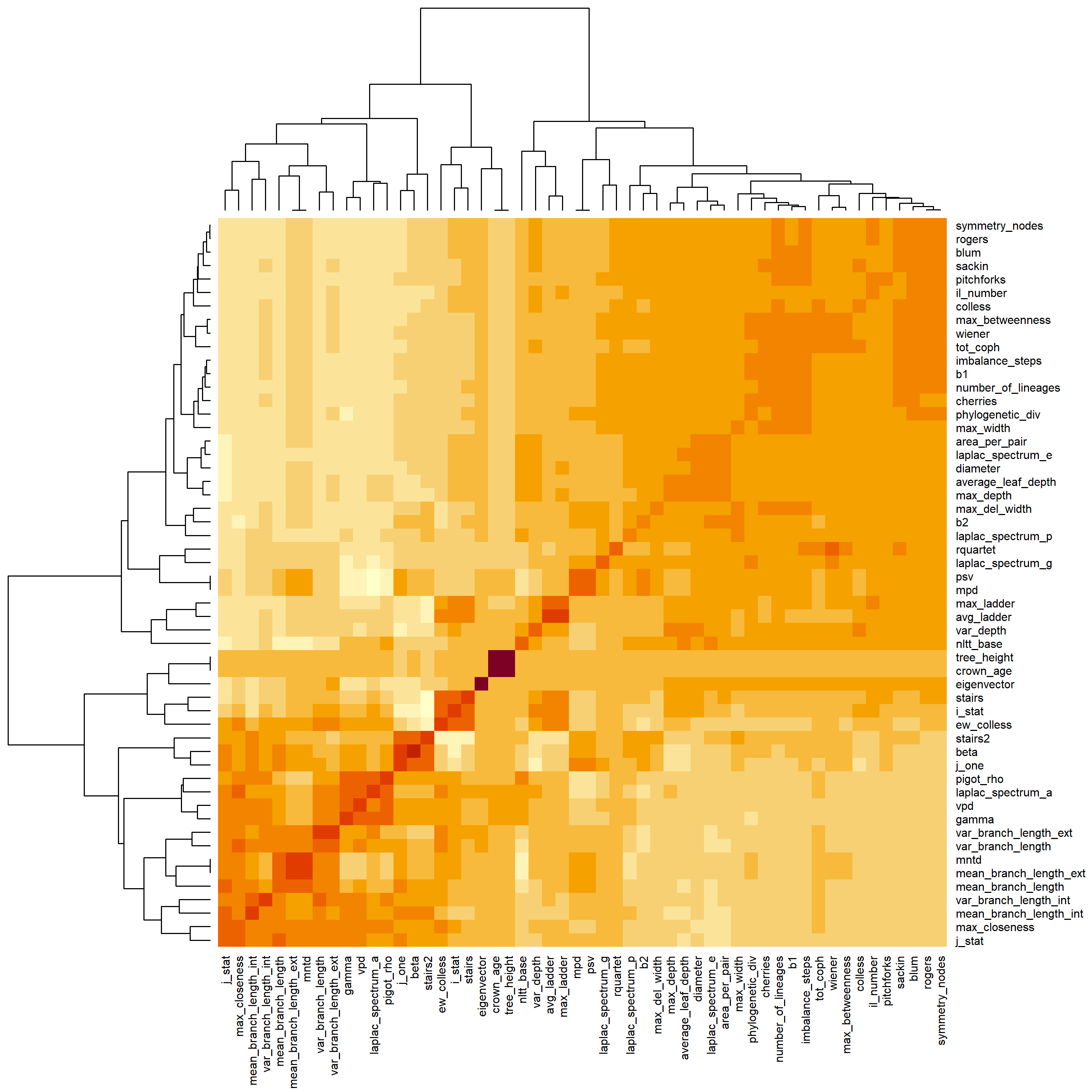

### Appendix B heatmap_NND.png

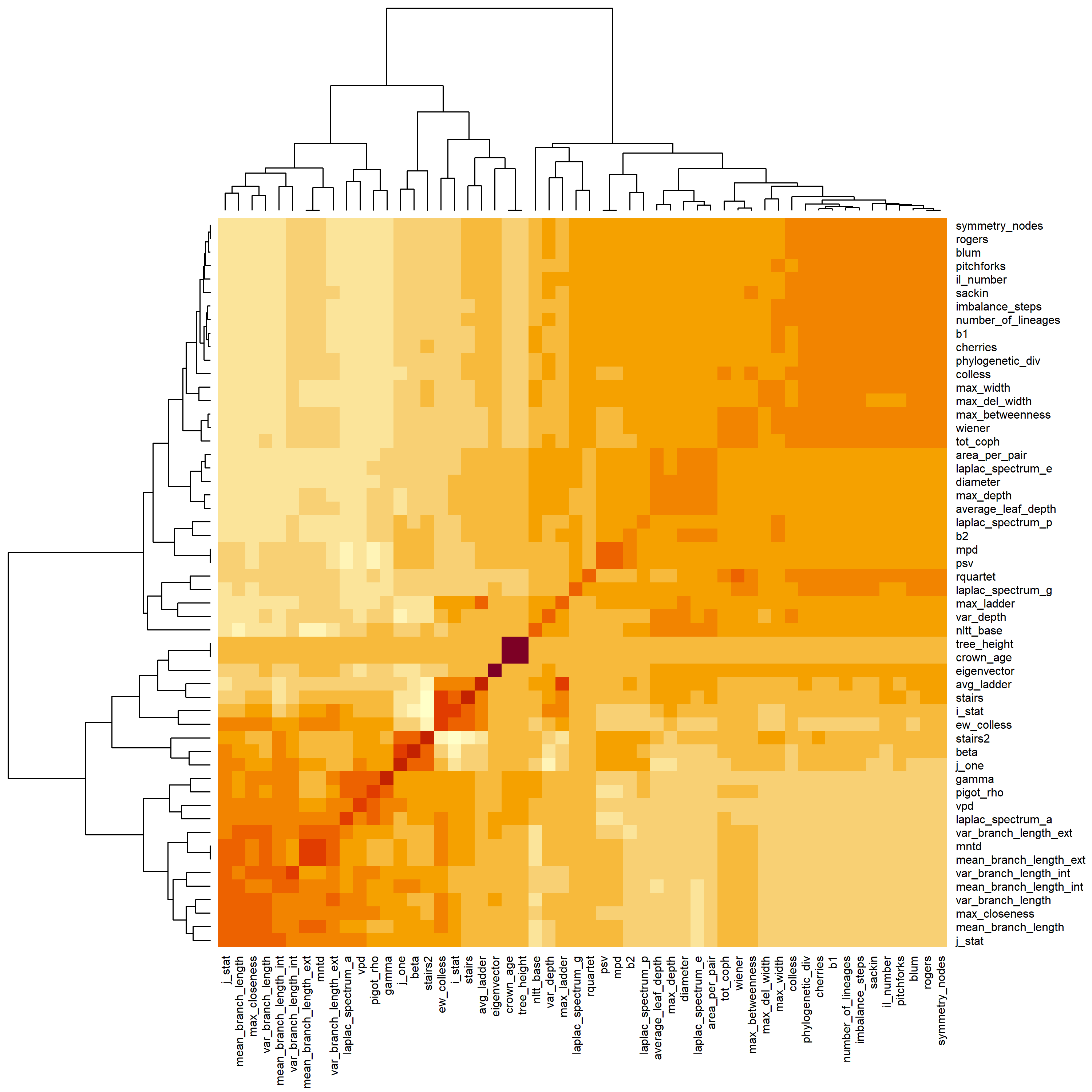

### Appendix B heatmap_PD.png

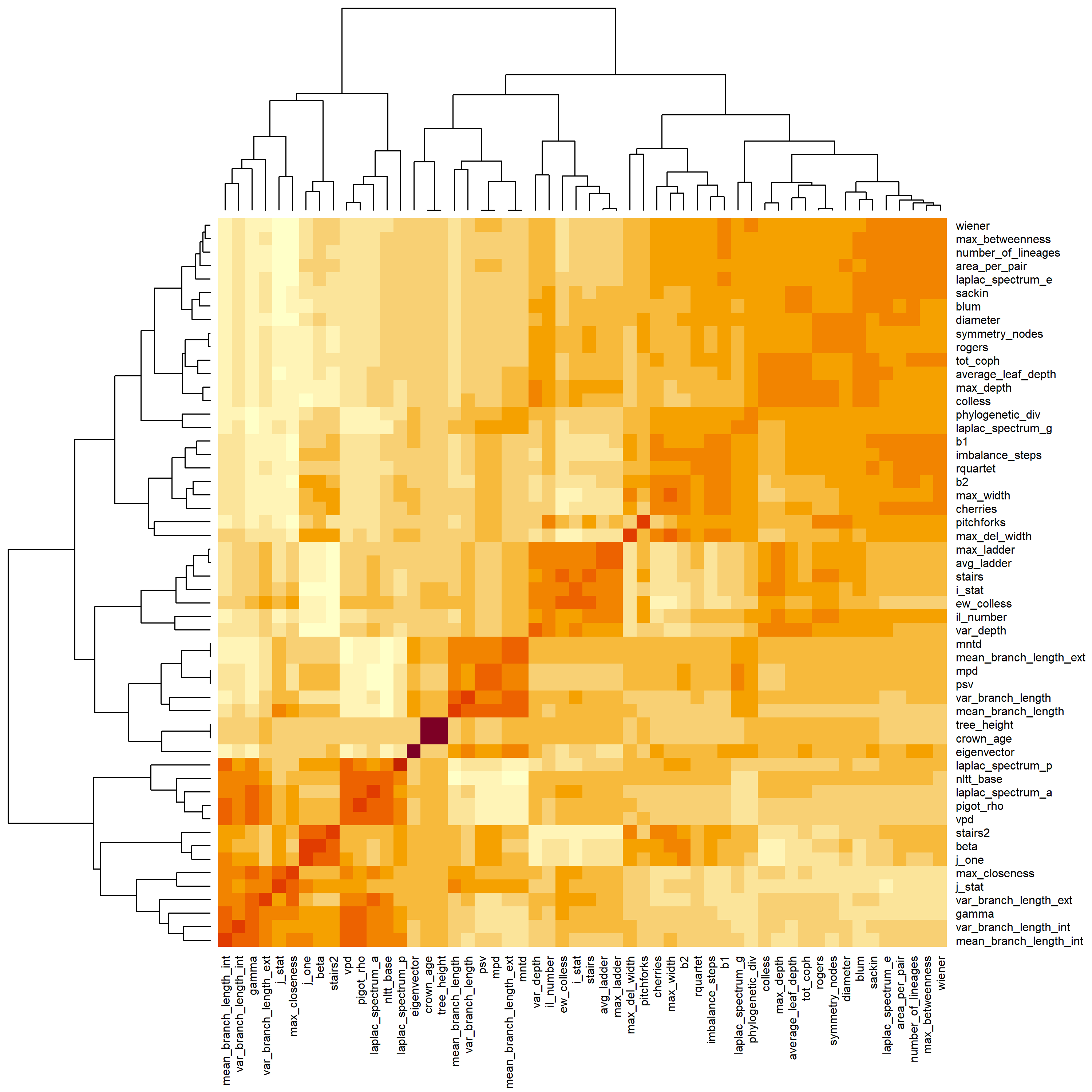

### Appendix D 0.4_0.1_0.png

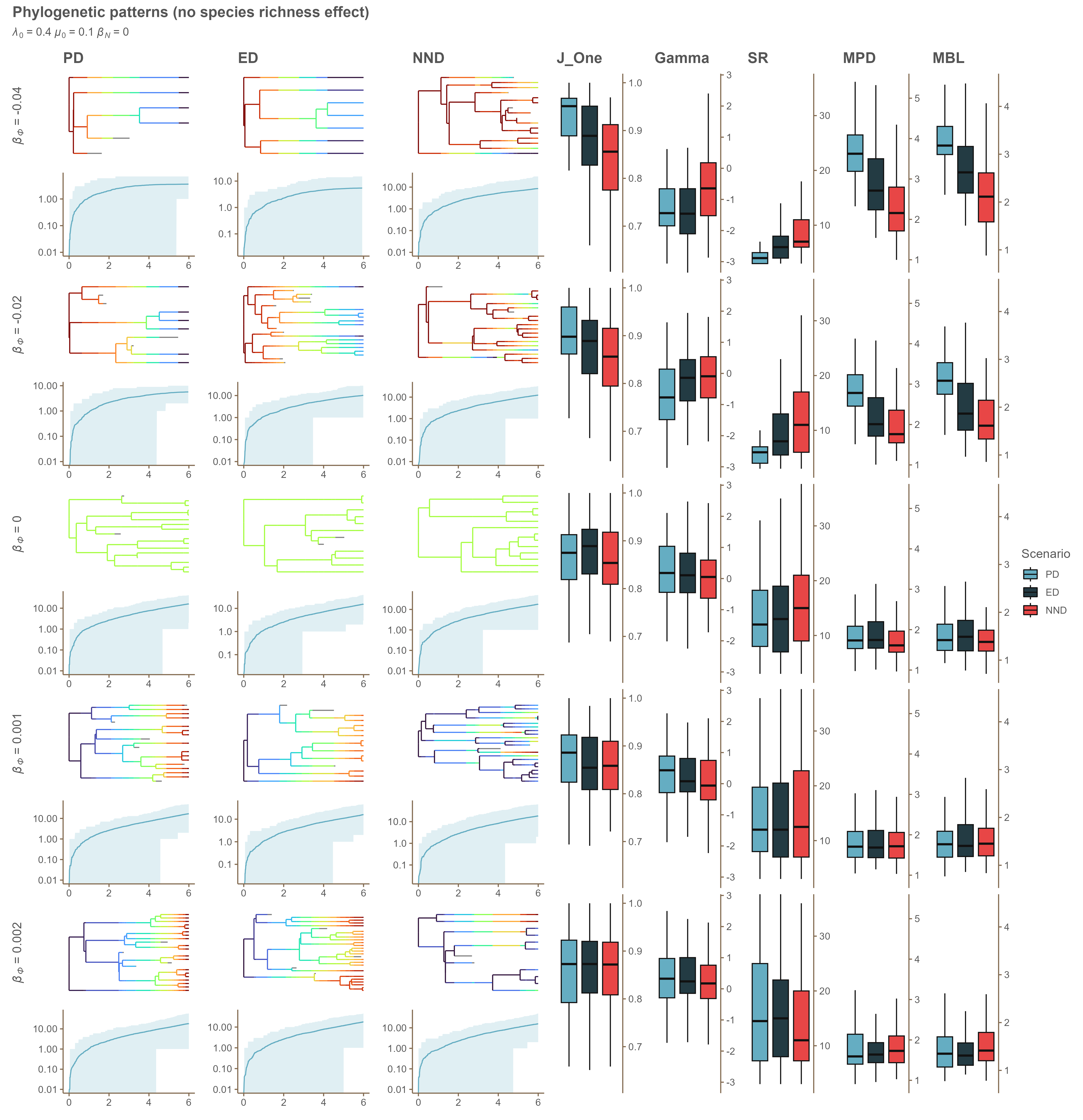

### Appendix D 0.4_0.1_-0.02.png

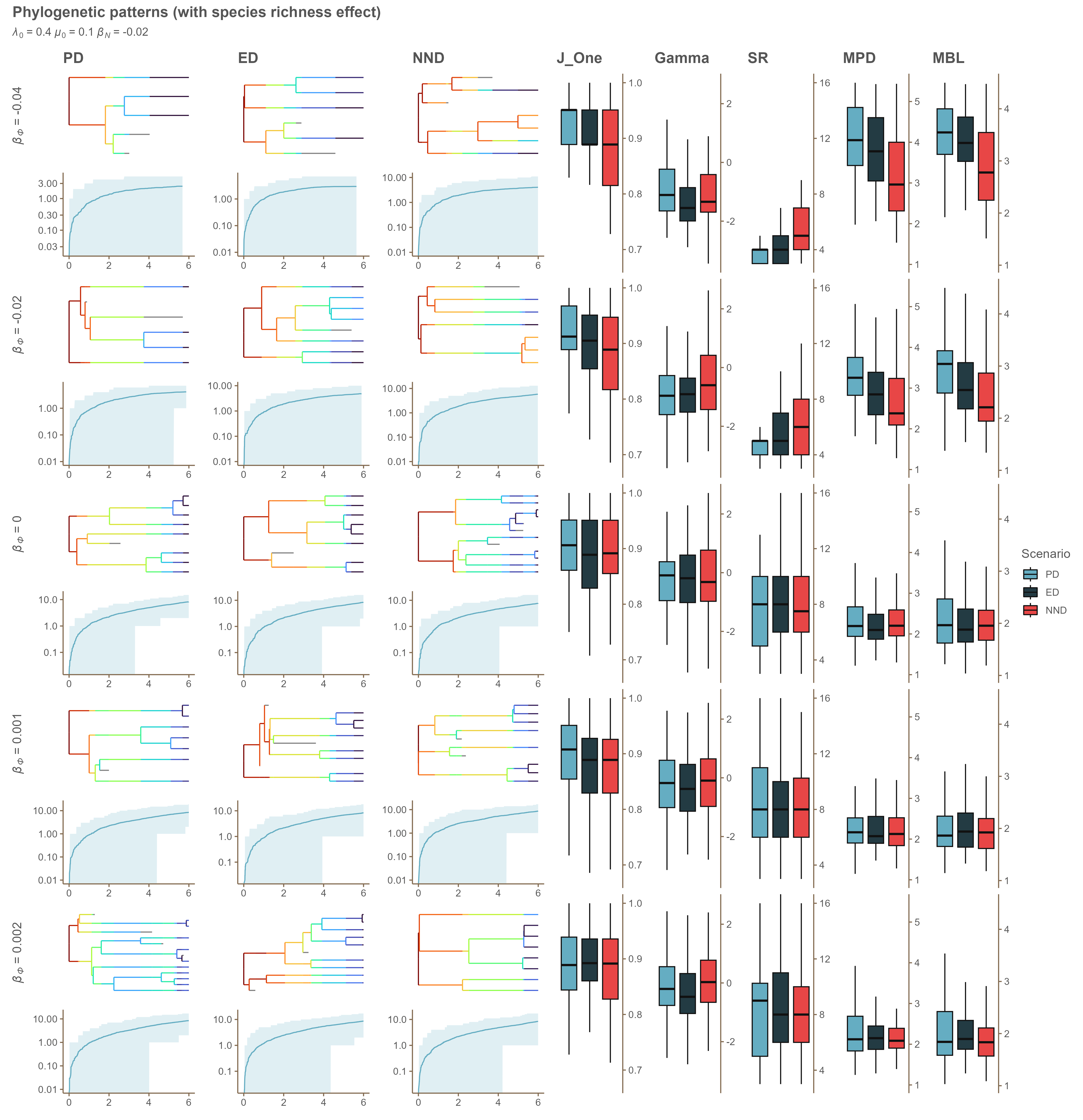

### Appendix D 0.4_0.1_-0.04.png

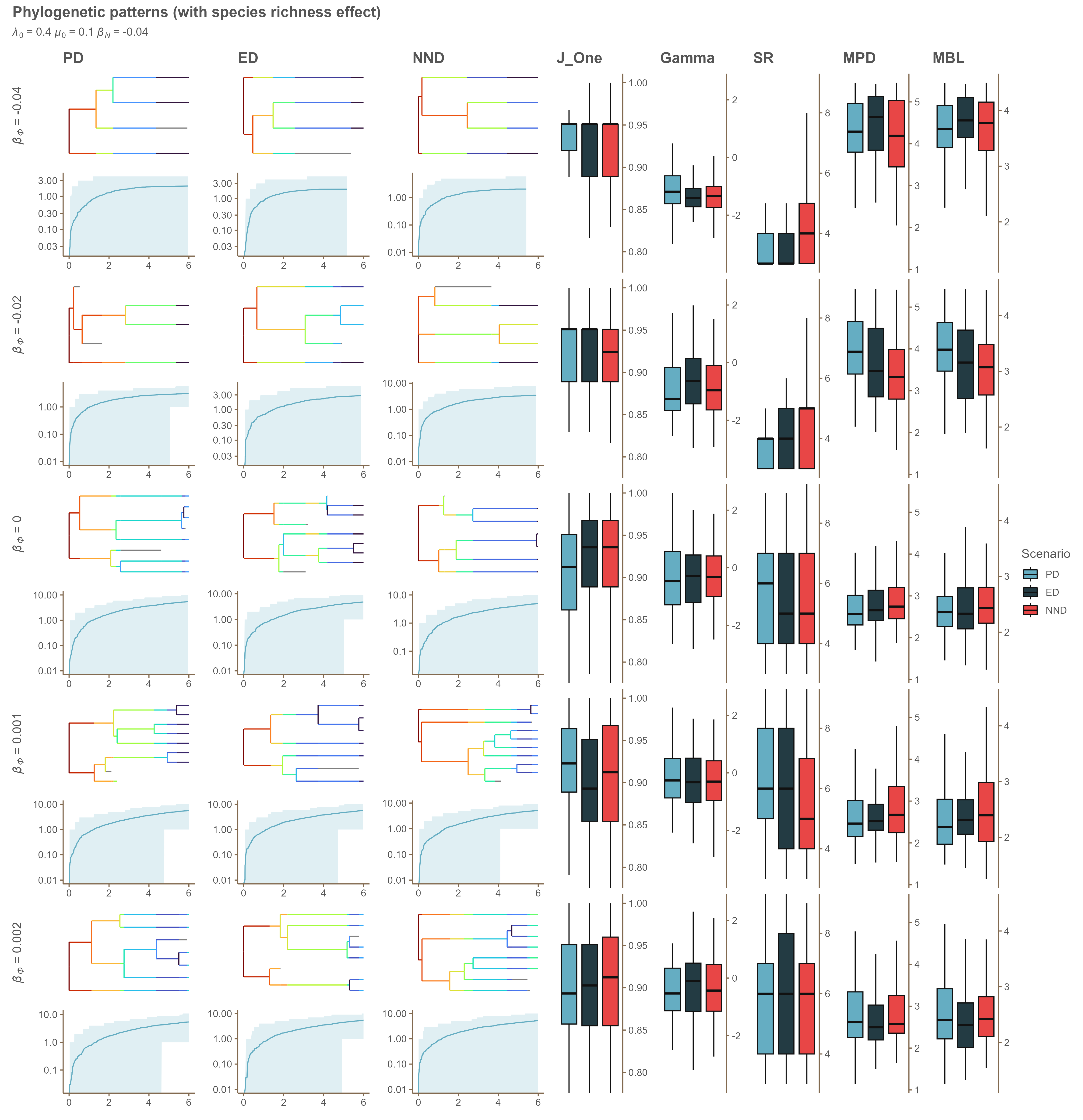

### Appendix D 0.4_0.2_0.png

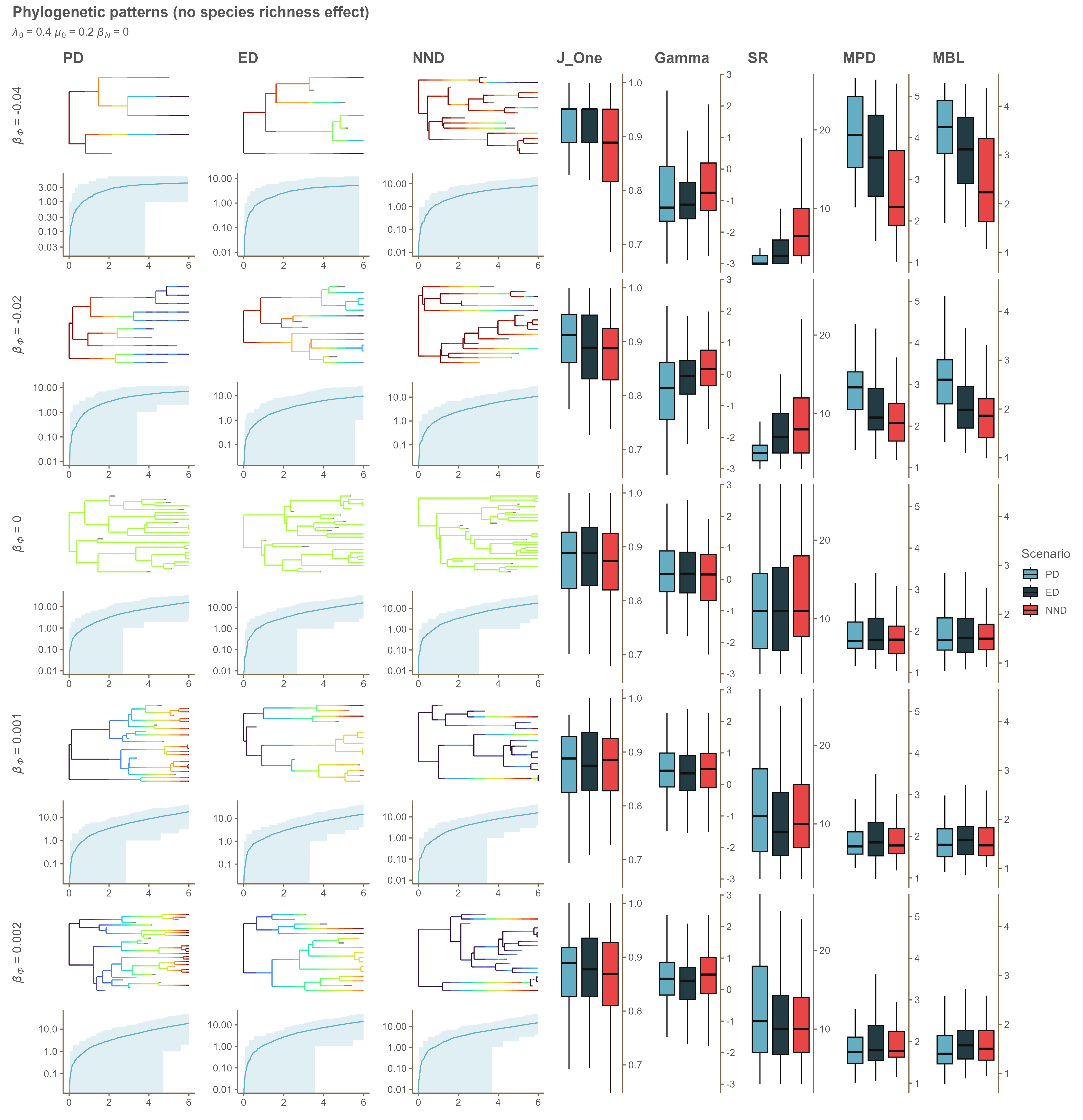

### Appendix D 0.4_0.2_-0.02.png

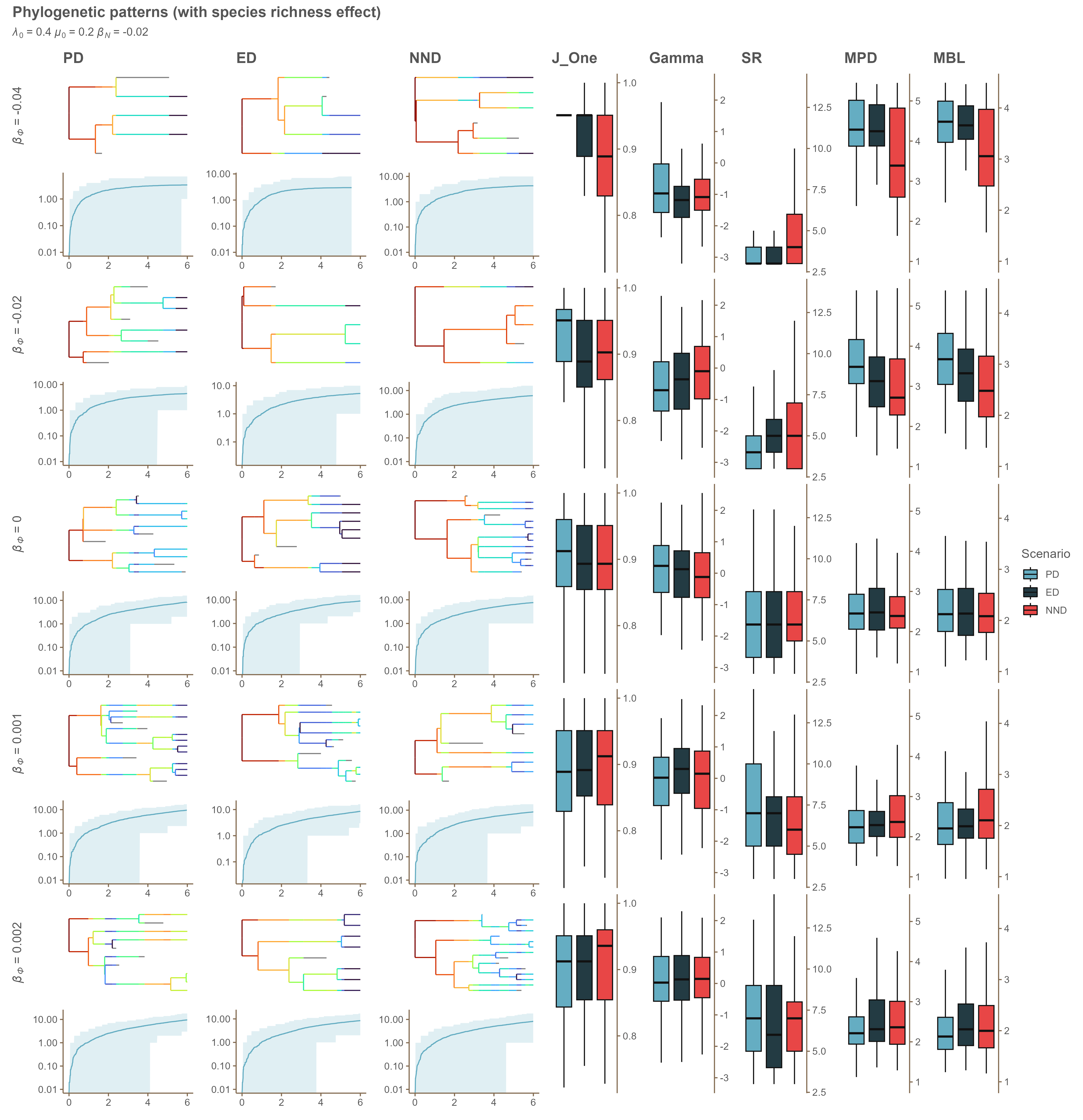

### Appendix D 0.4_0.2_-0.04.png

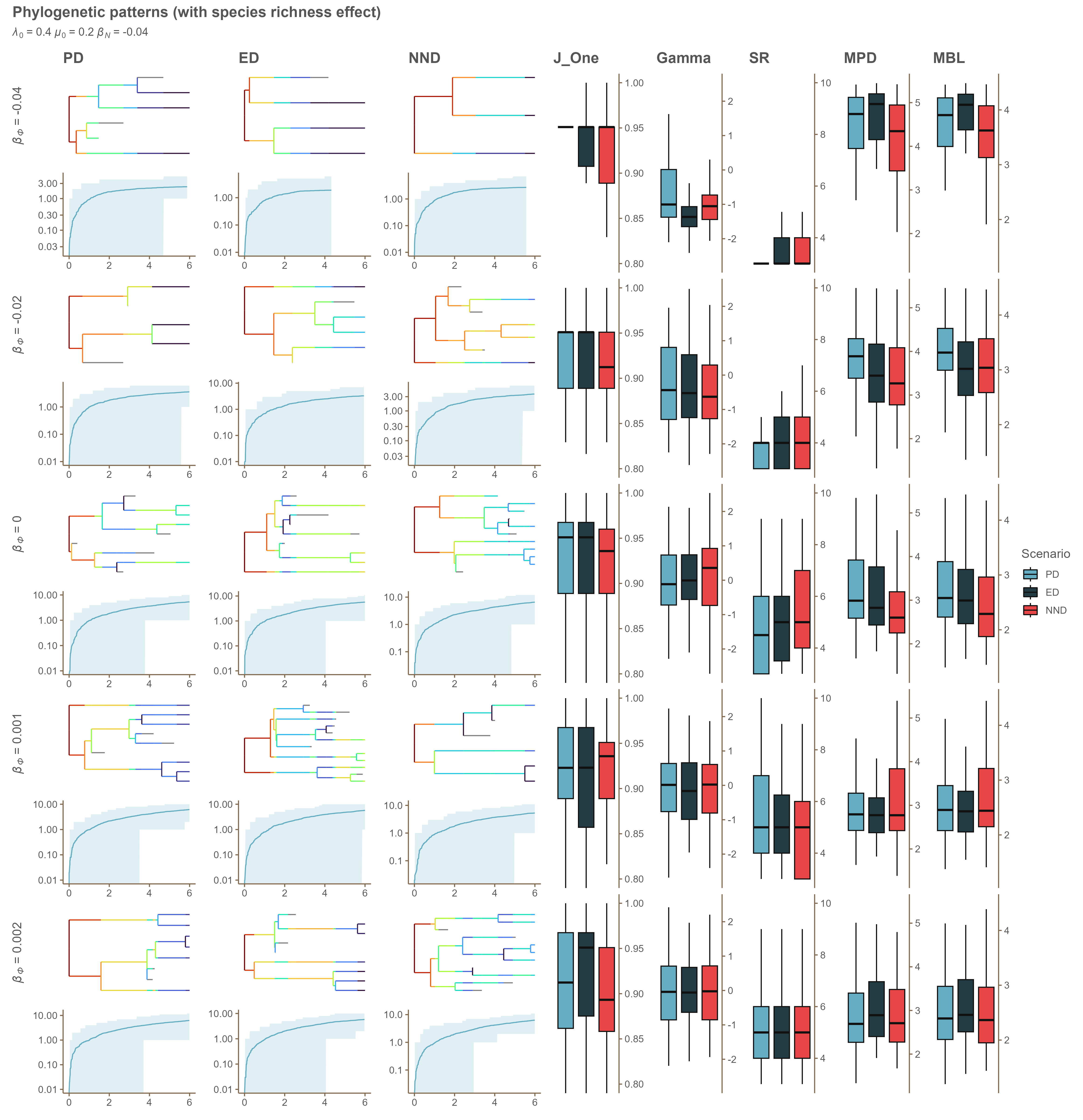

### Appendix D 0.4_0_0.png

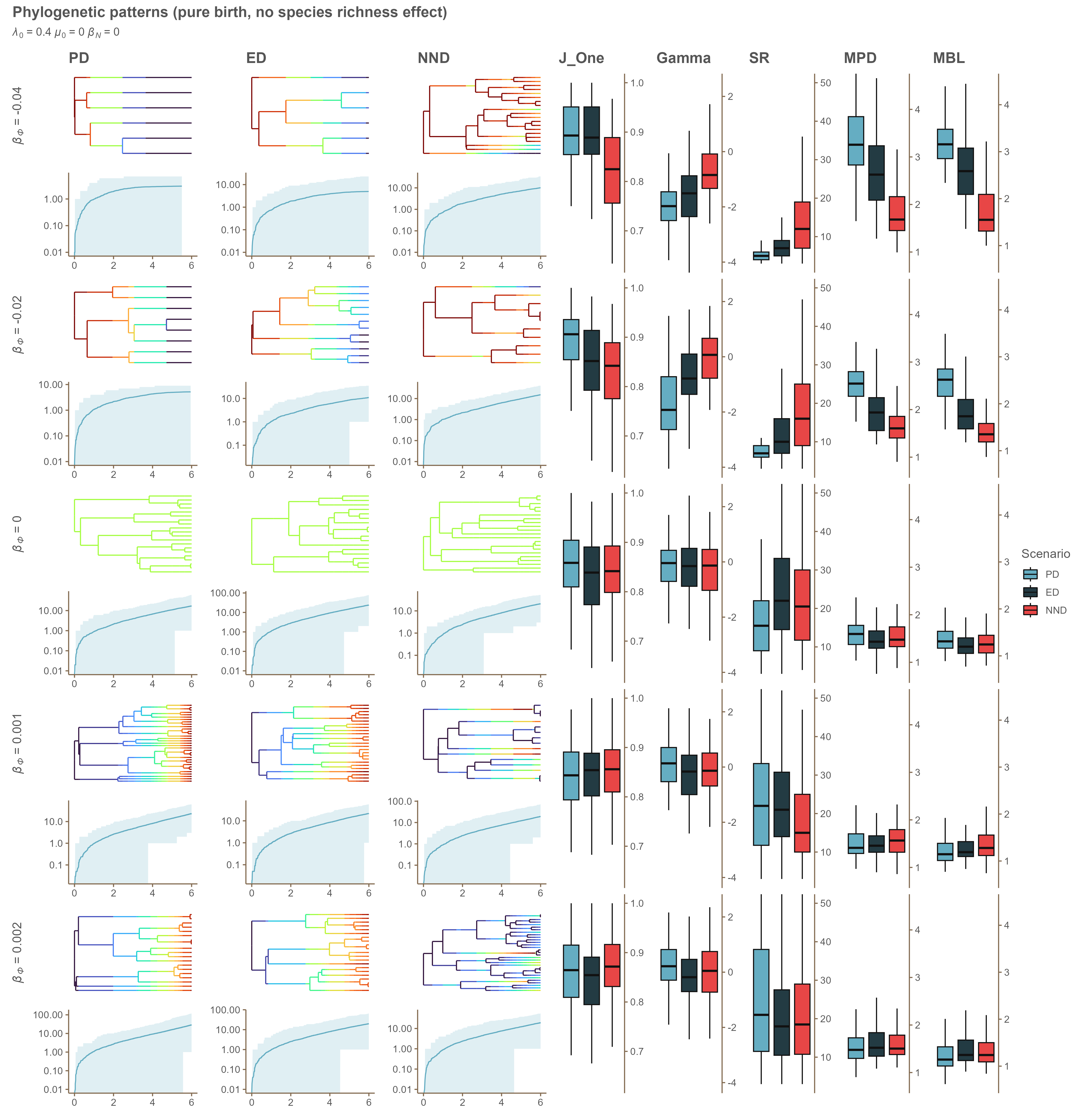

### Appendix D 0.4_0_-0.02.png

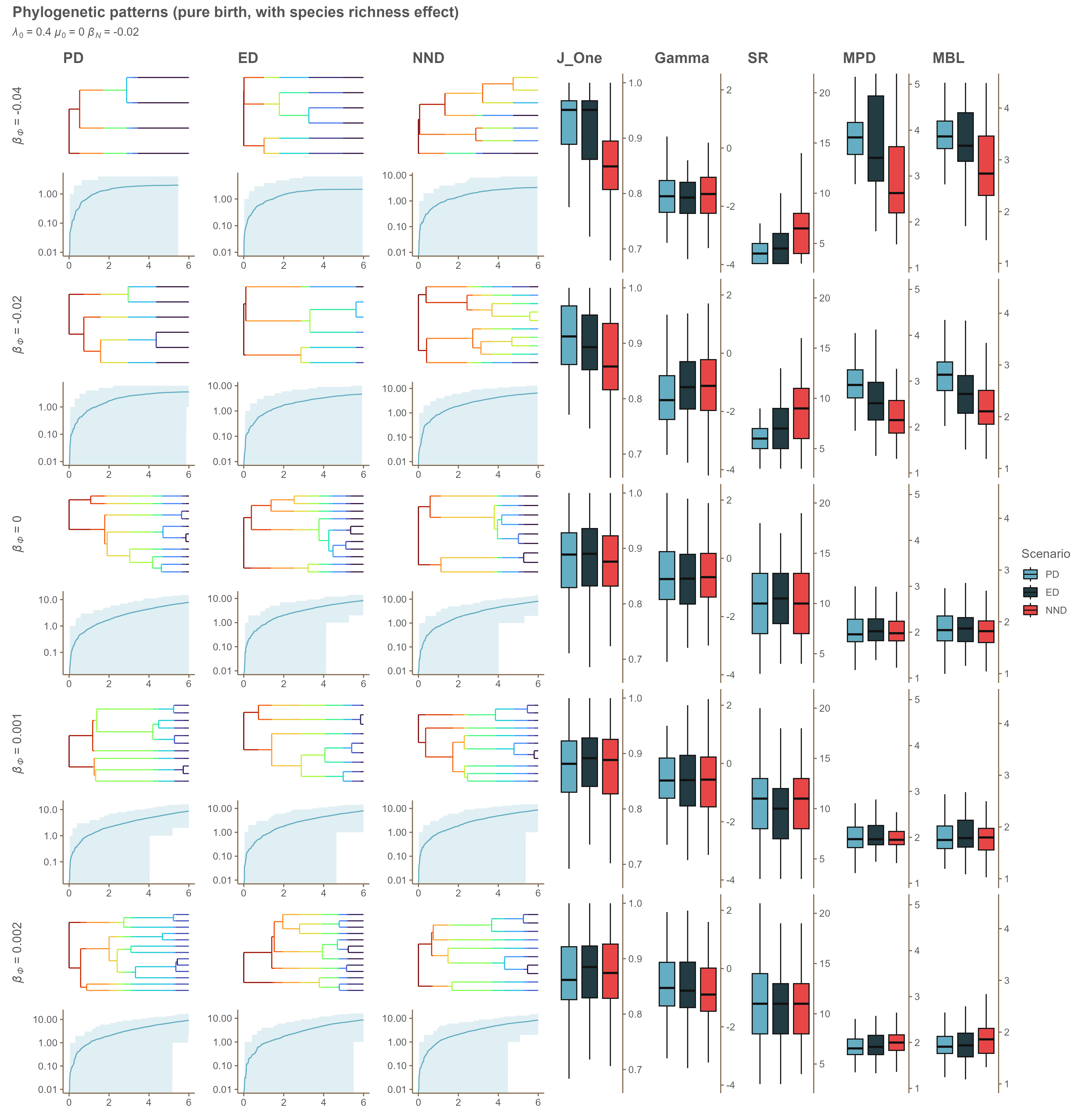

### Appendix D 0.4_0_-0.04.png

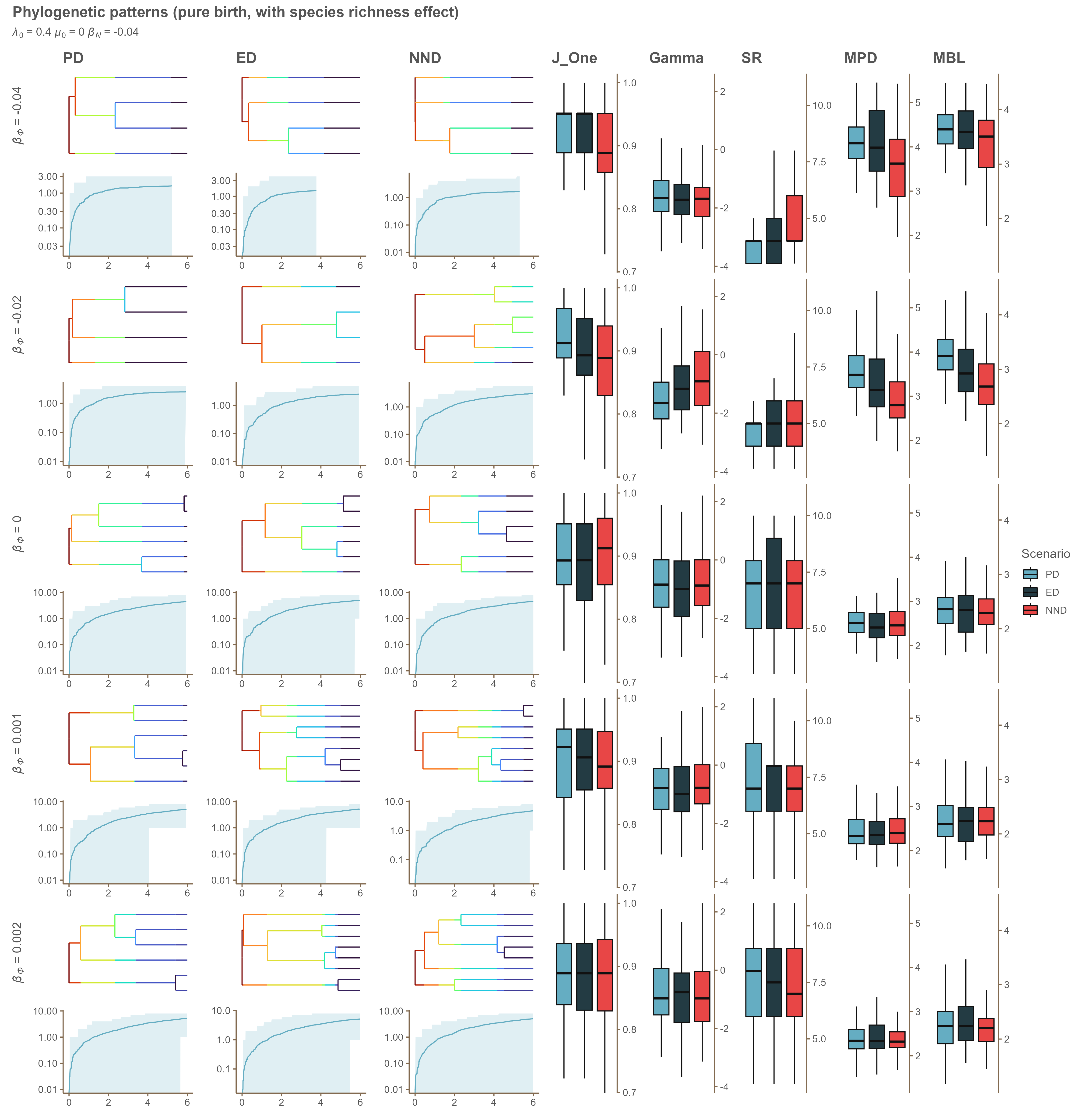

### Appendix D 0.5_0.1_0.png

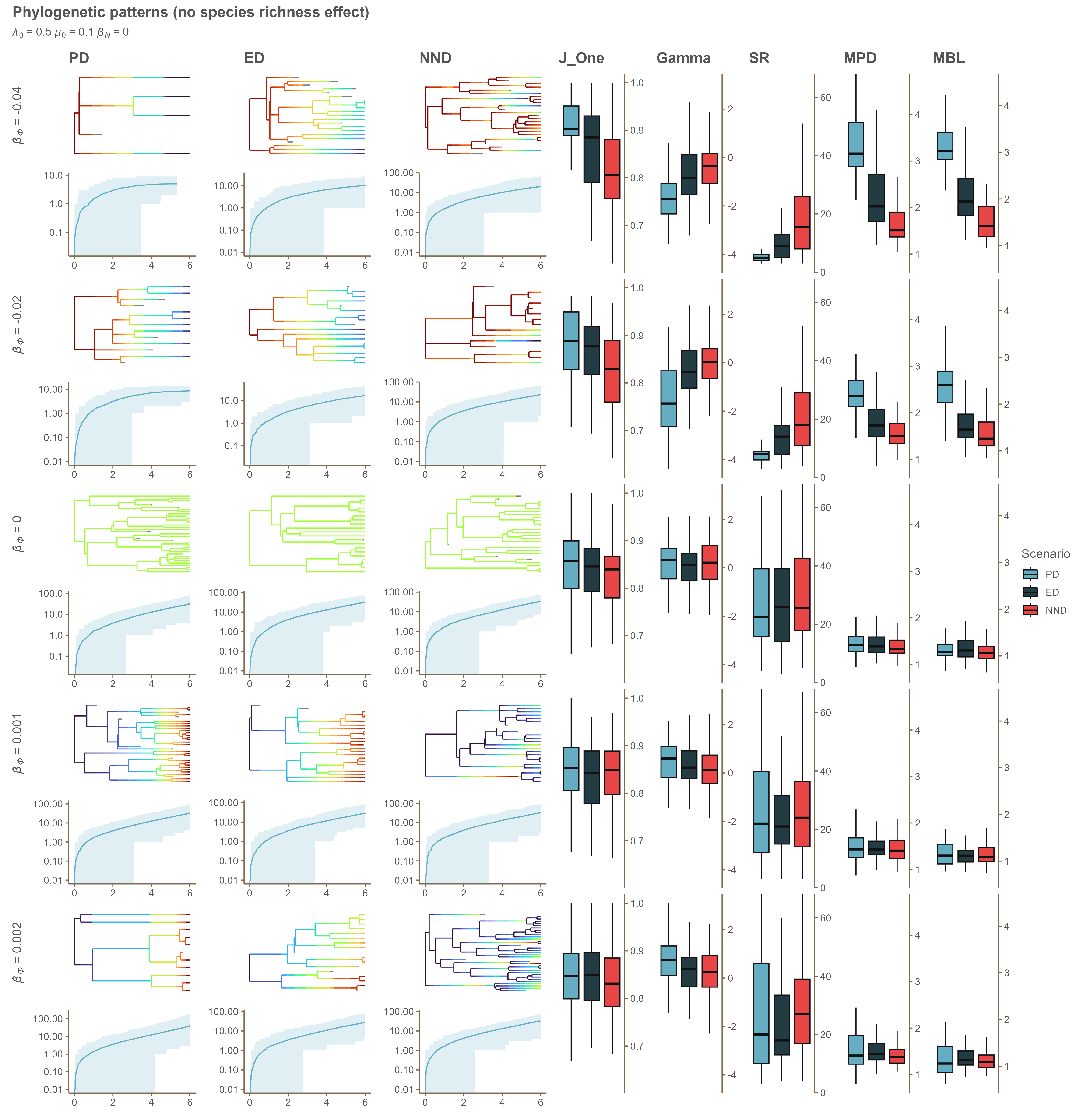

### Appendix D 0.5_0.1_-0.02.png

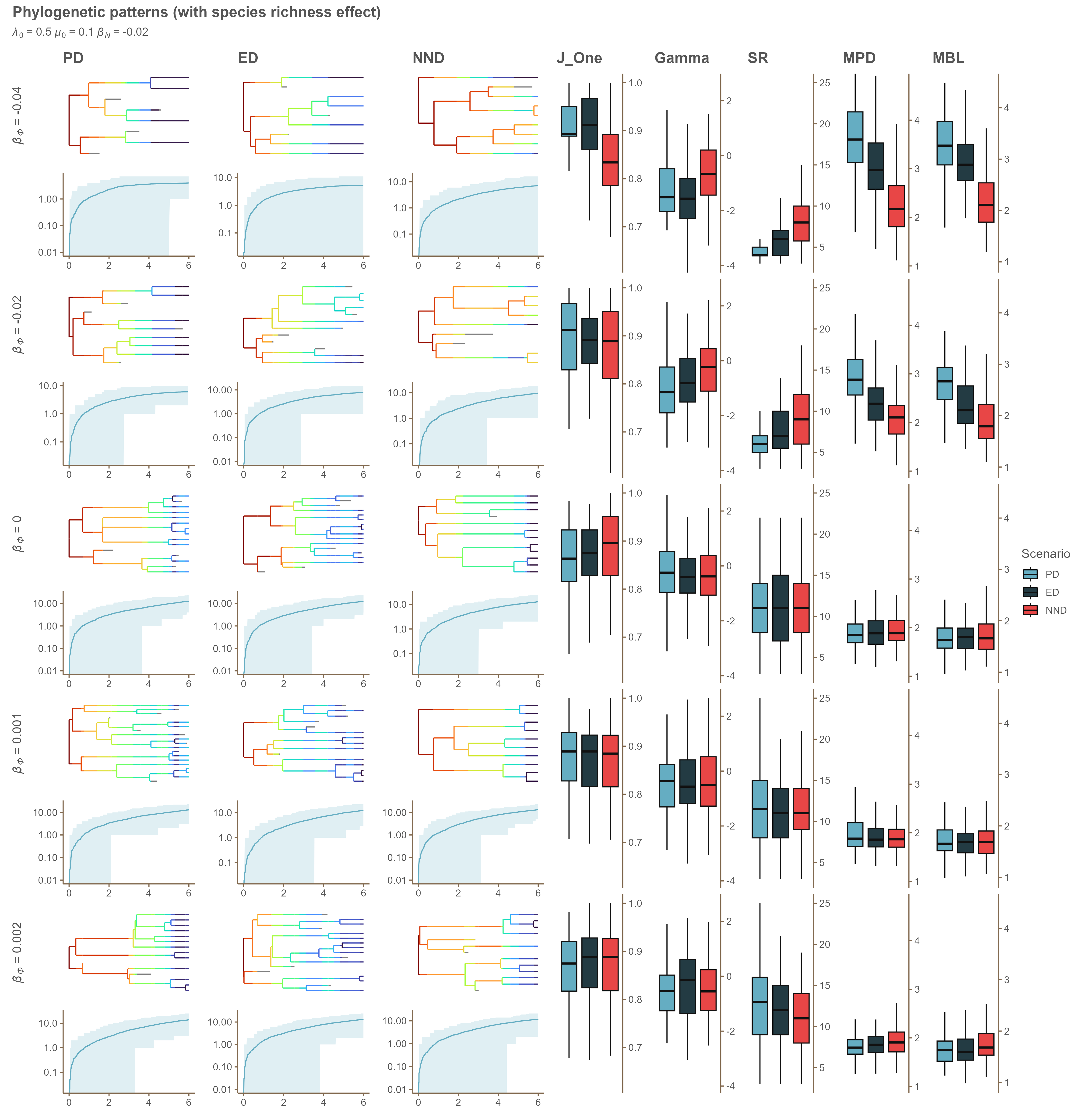

### Appendix D 0.5_0.1_-0.04.png

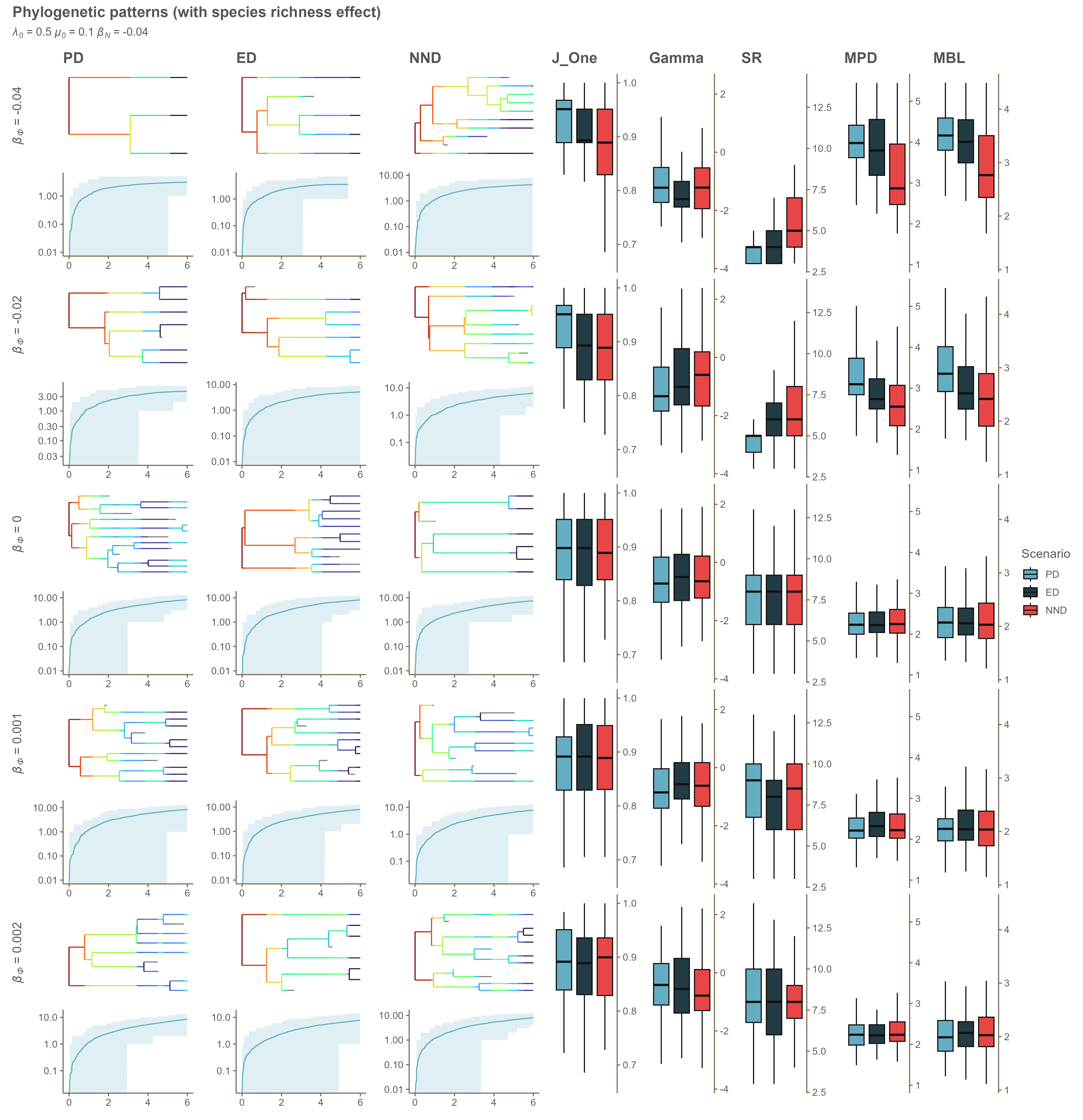

### Appendix D 0.5_0.2_0.png

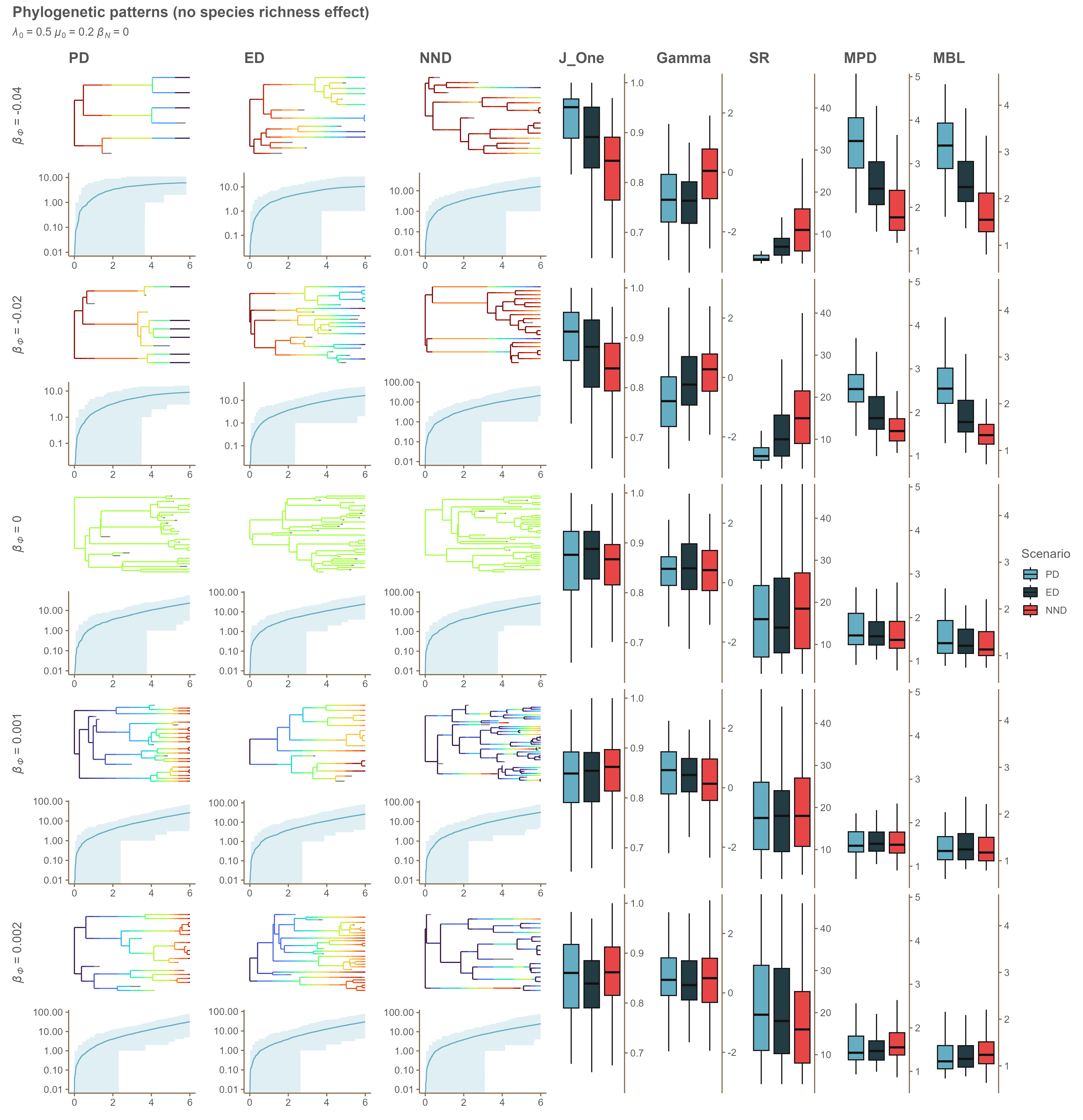

### Appendix D 0.5_0.2_-0.02.png

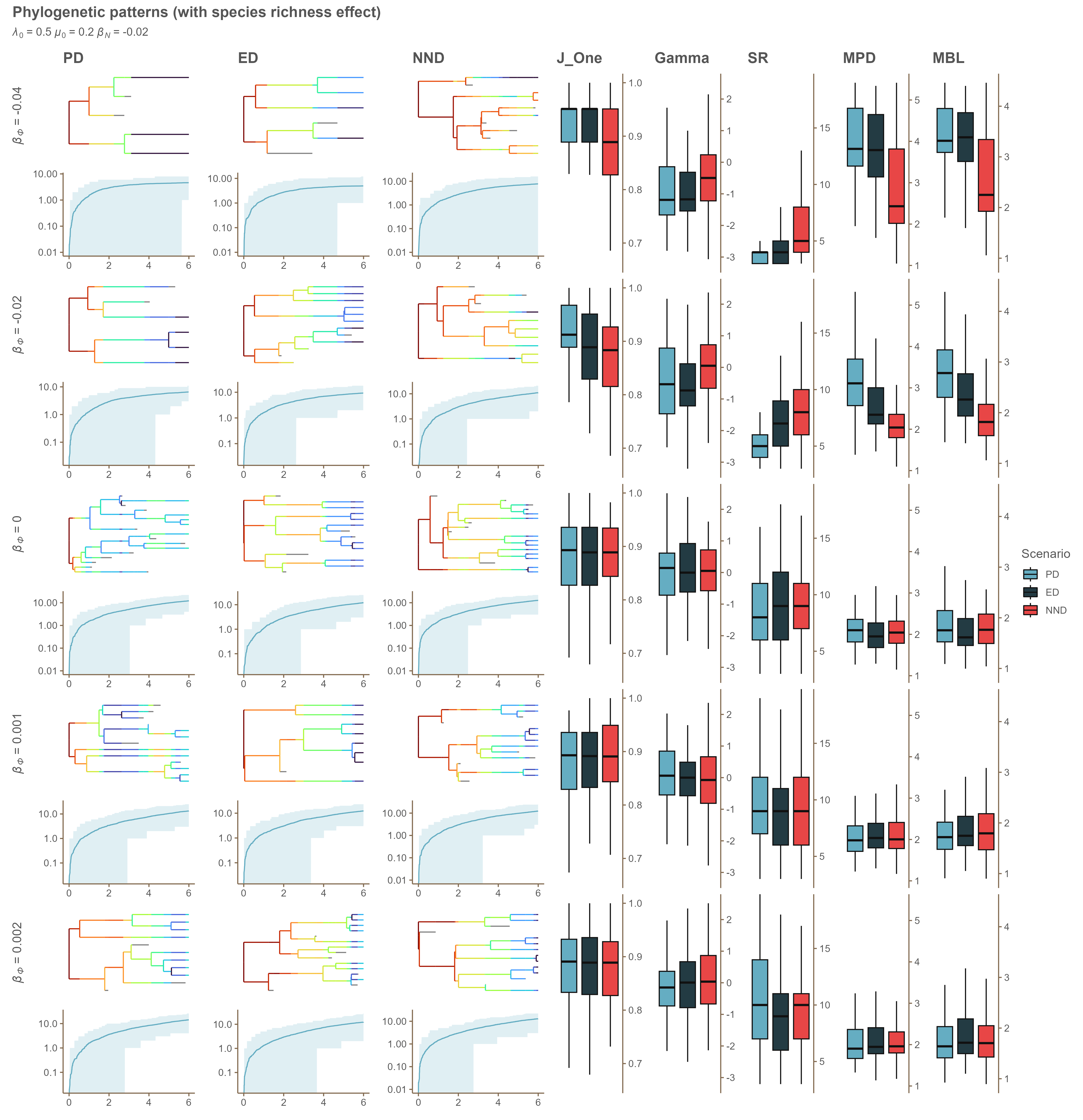

### Appendix D 0.5_0.2_-0.04.png

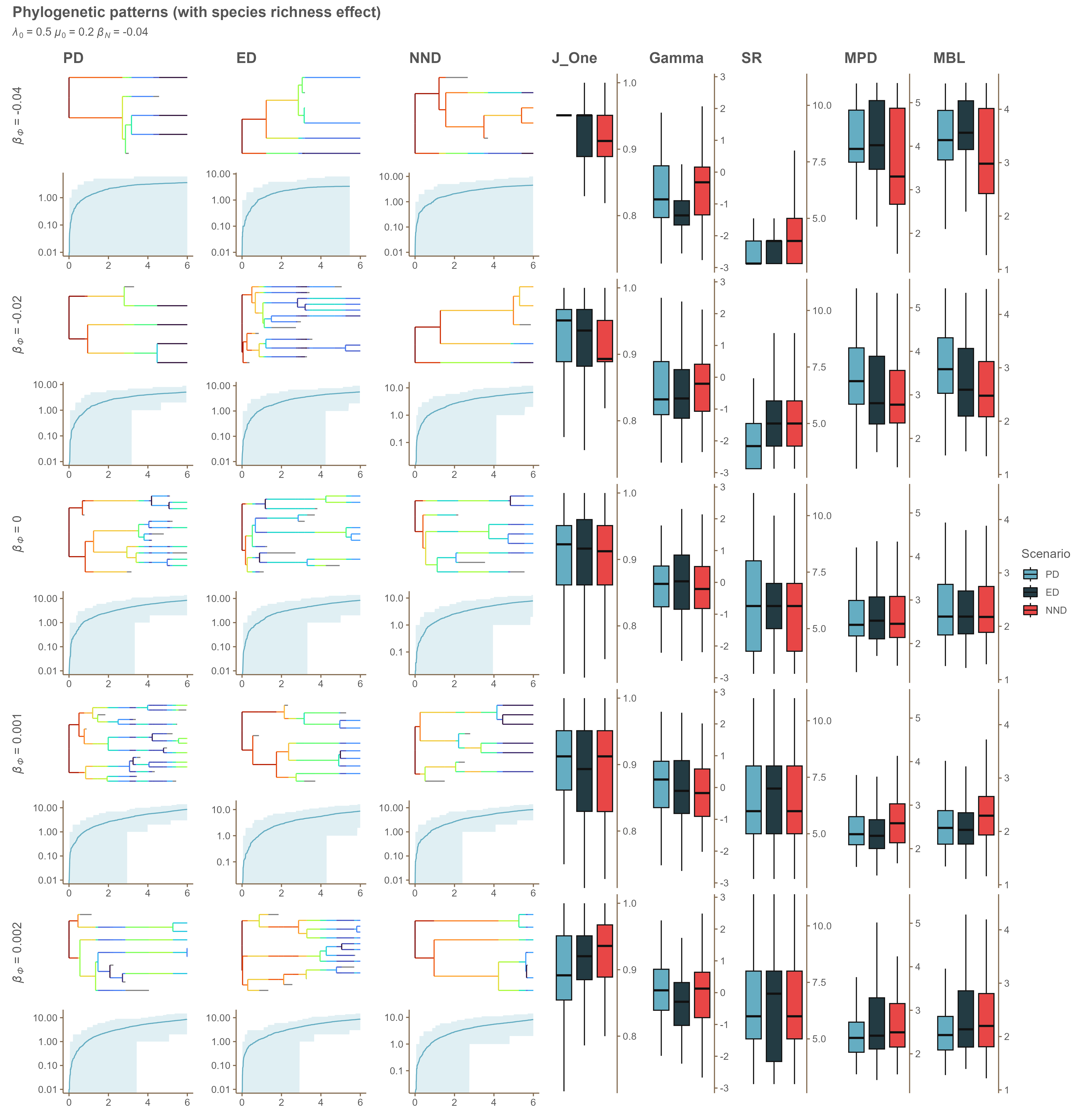

### Appendix D 0.5_0_0.png

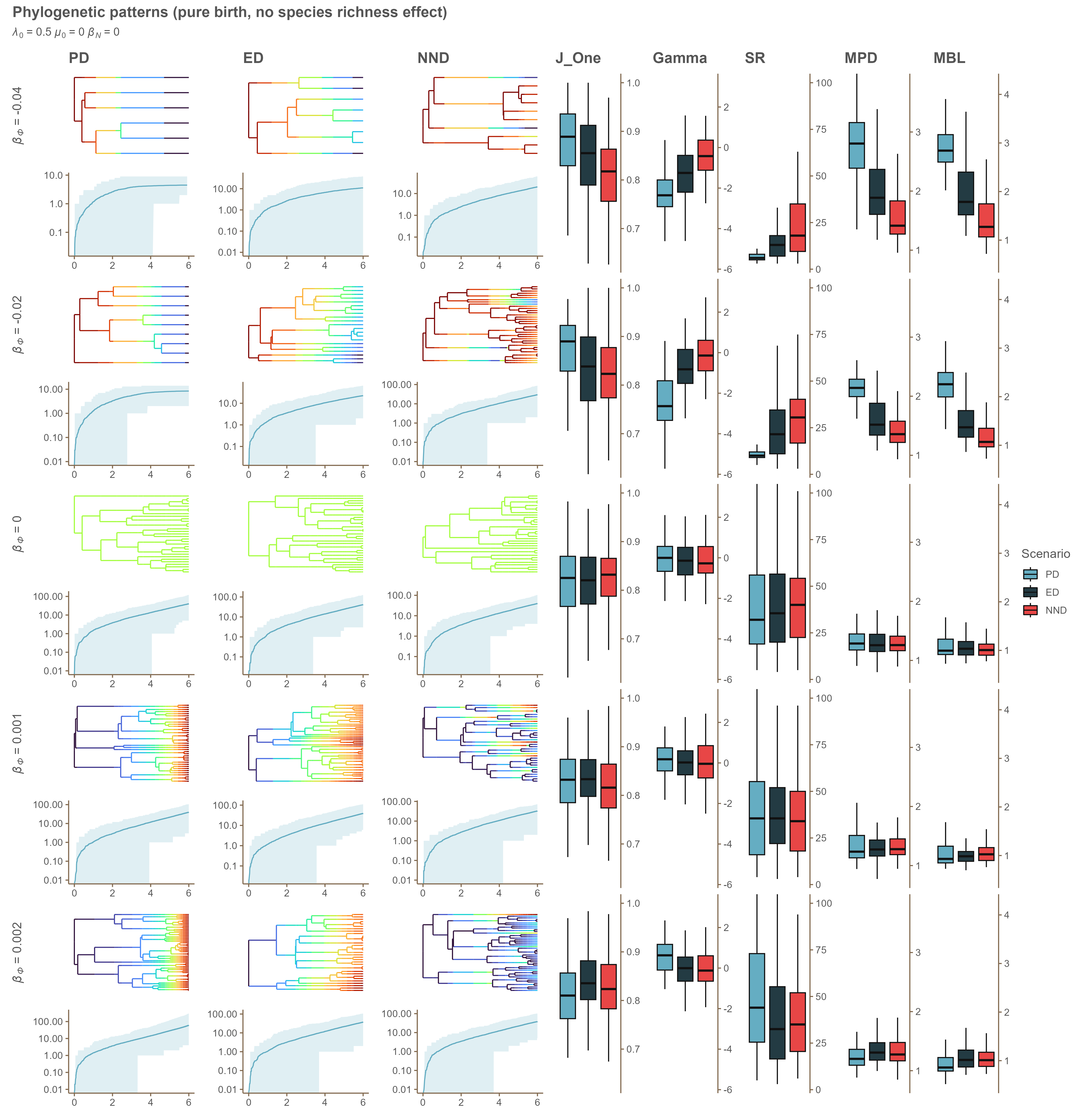

### Appendix D 0.5_0_-0.02.png

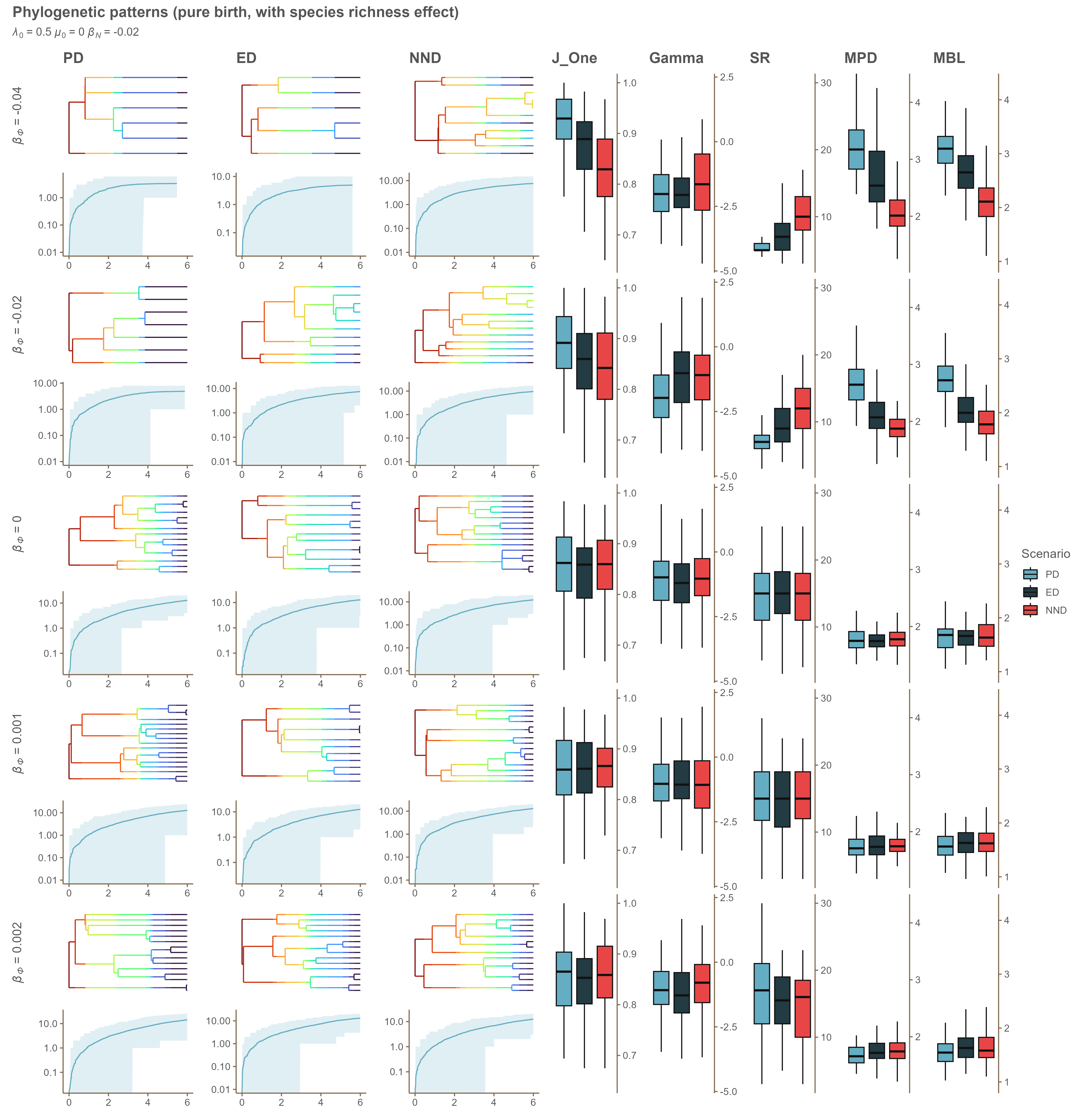

### Appendix D 0.5_0_-0.04.png

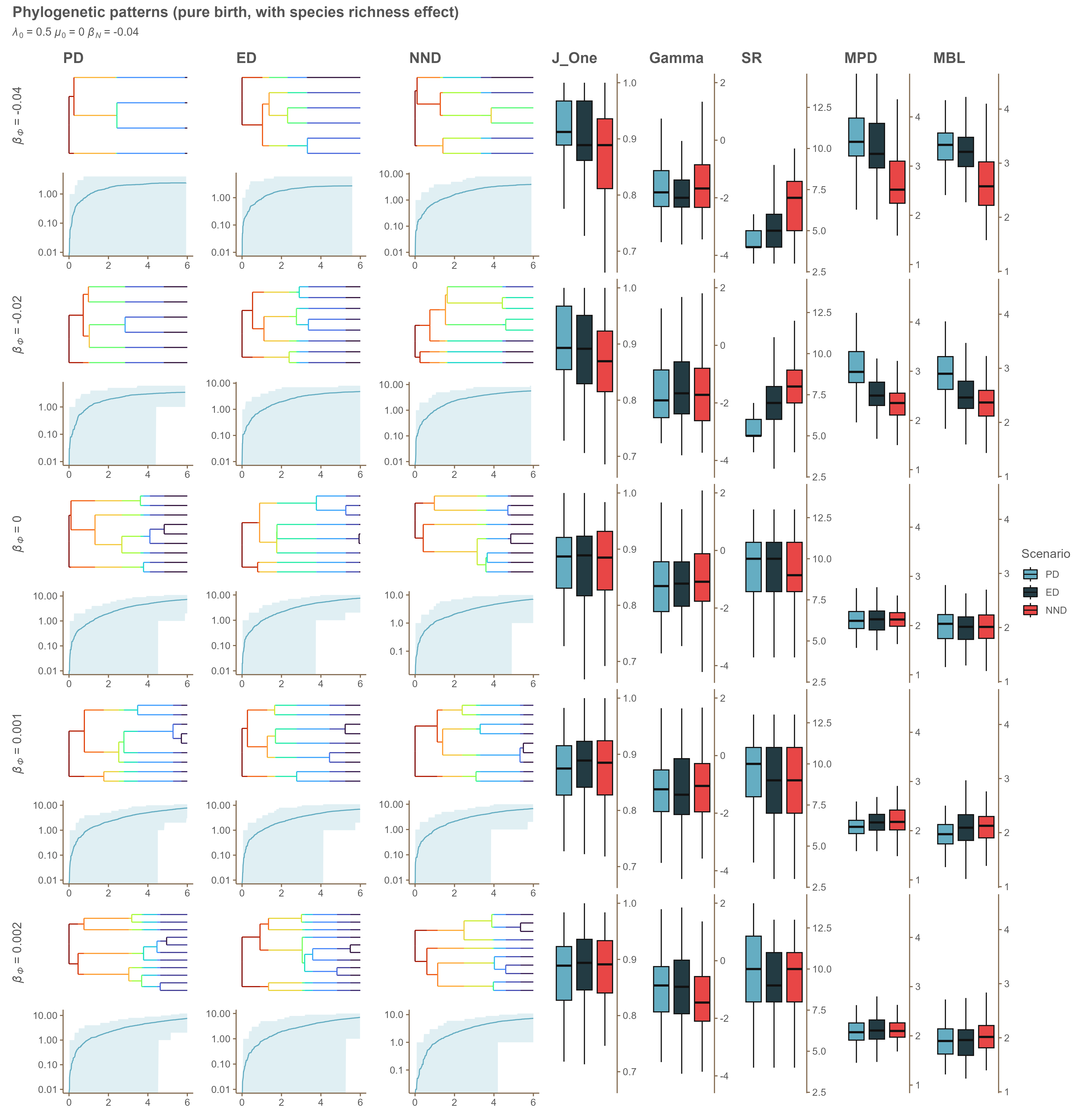

### Appendix D 0.6_0.1_0.png

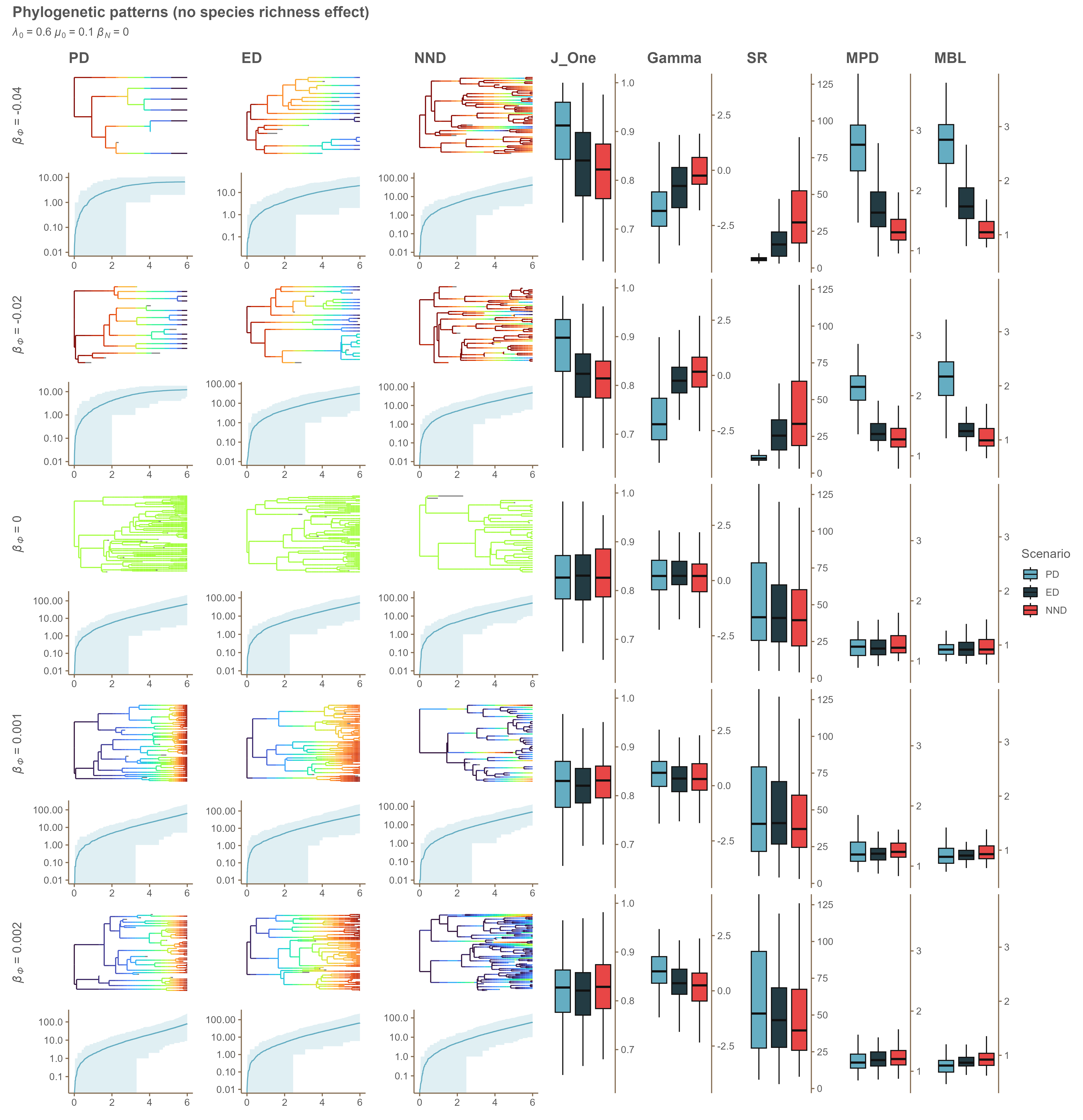

### Appendix D 0.6_0.1_-0.02.png

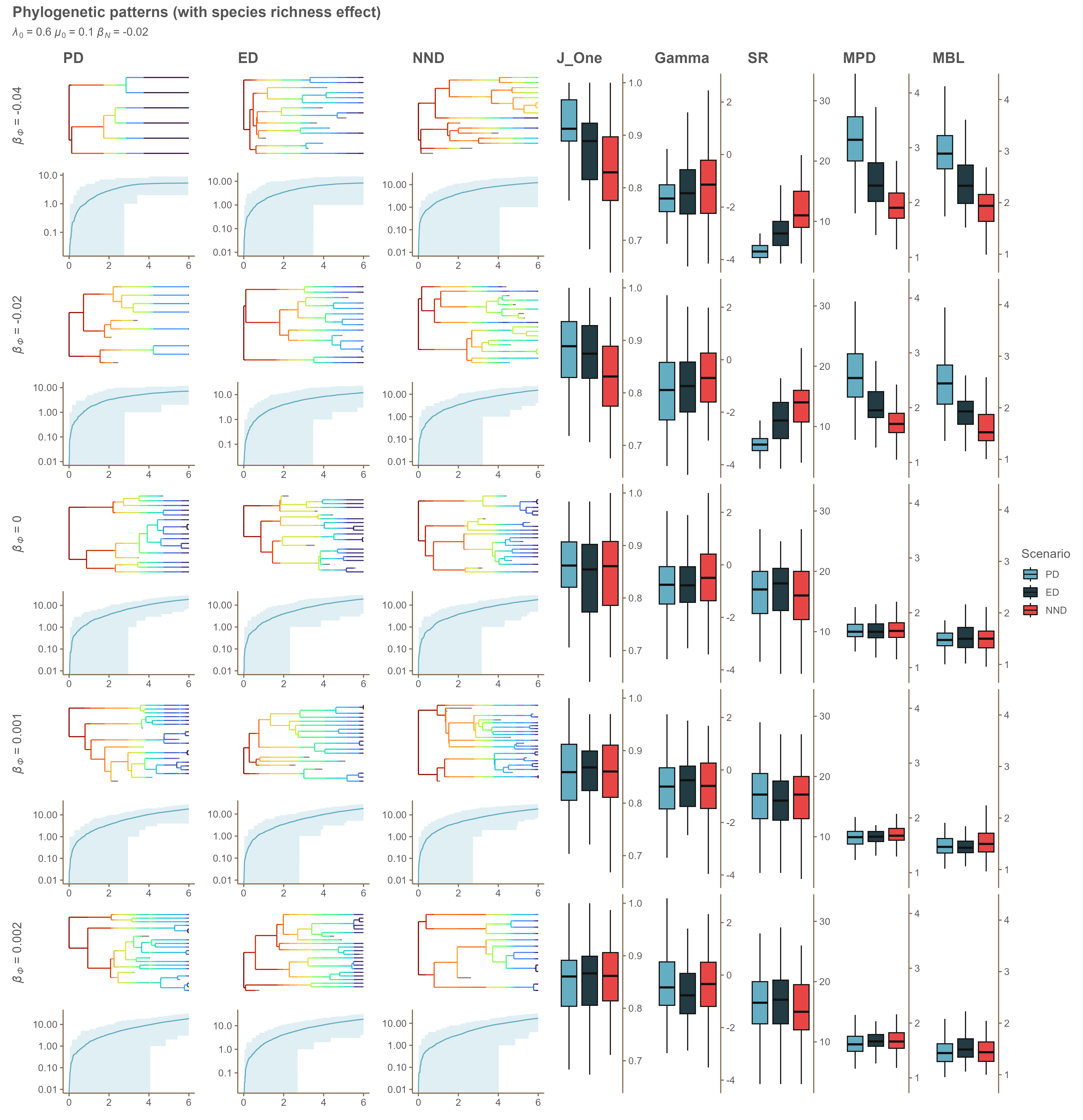

### Appendix D 0.6_0.1_-0.04.png

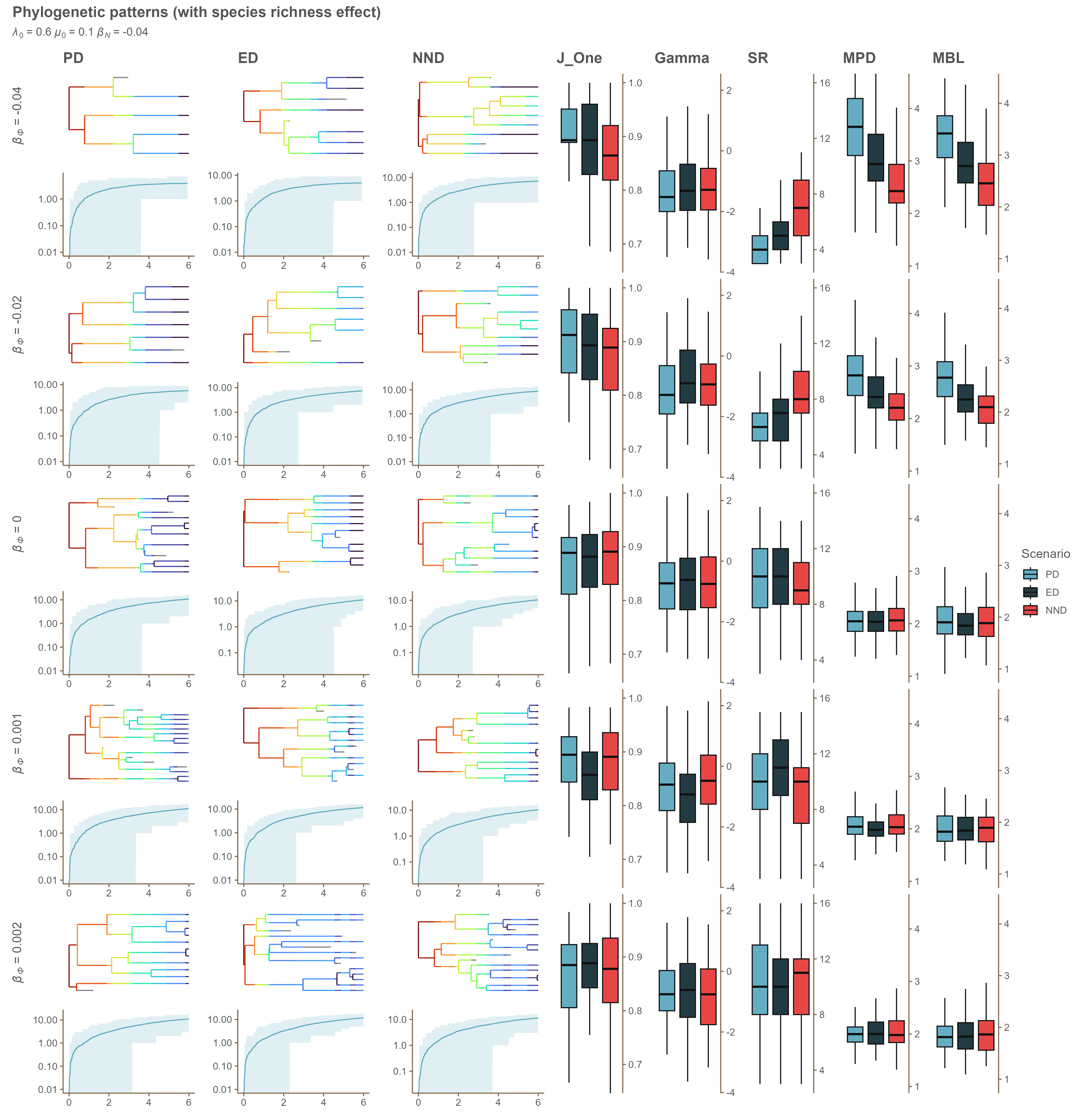

### Appendix D 0.6_0.2_0.png

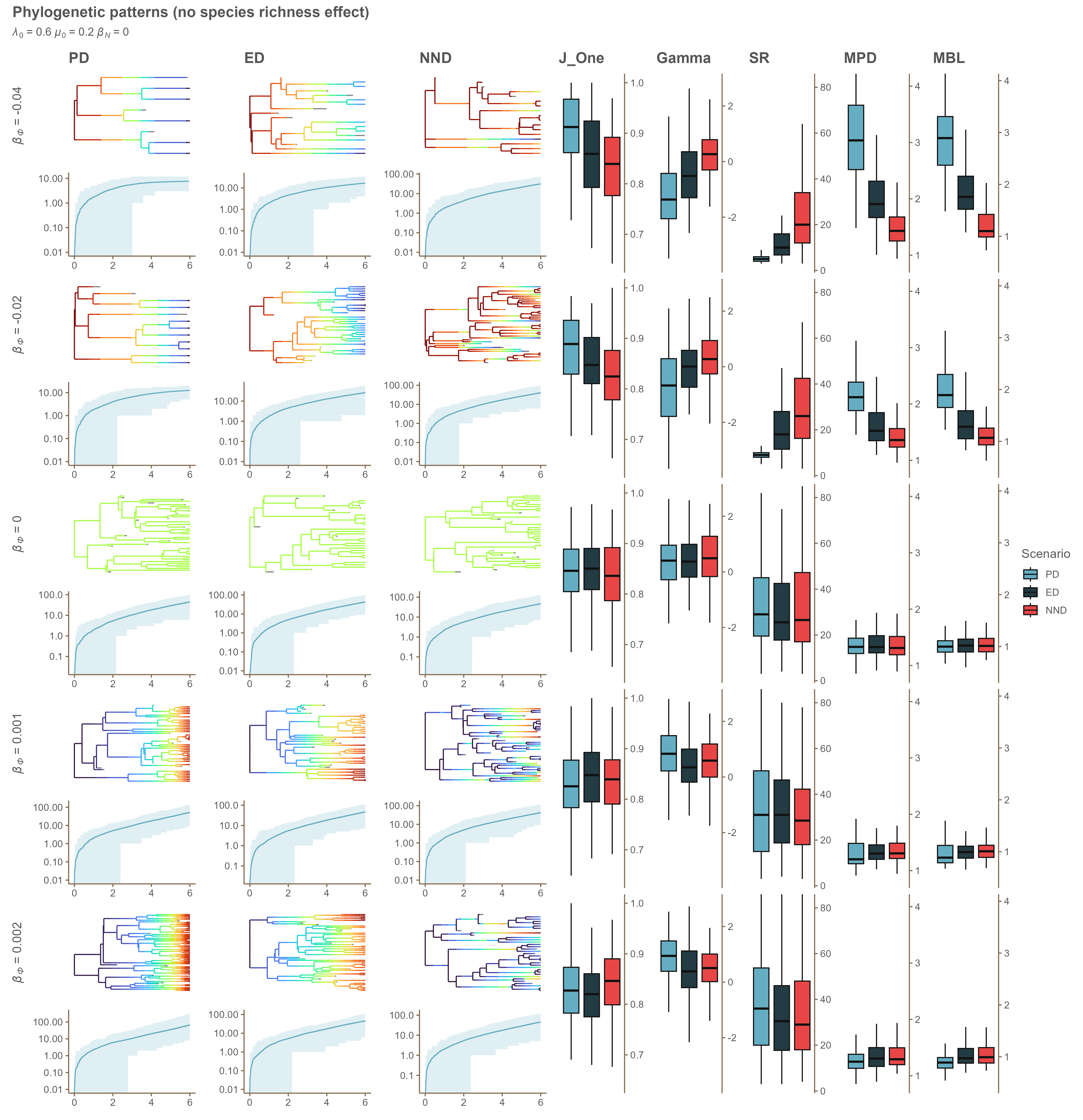

### Appendix D 0.6_0.2_-0.02.png

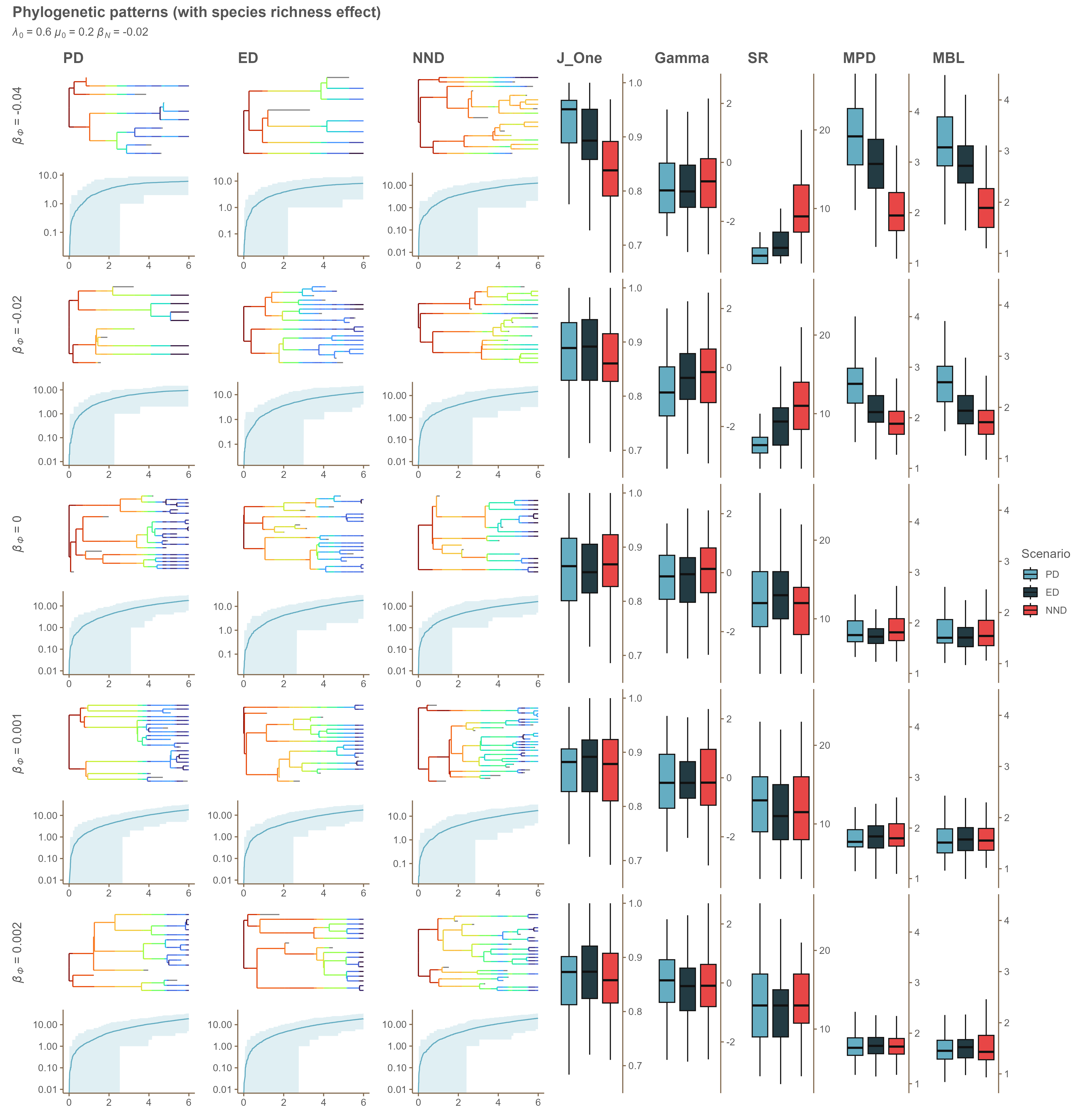

### Appendix D 0.6_0.2_-0.04.png

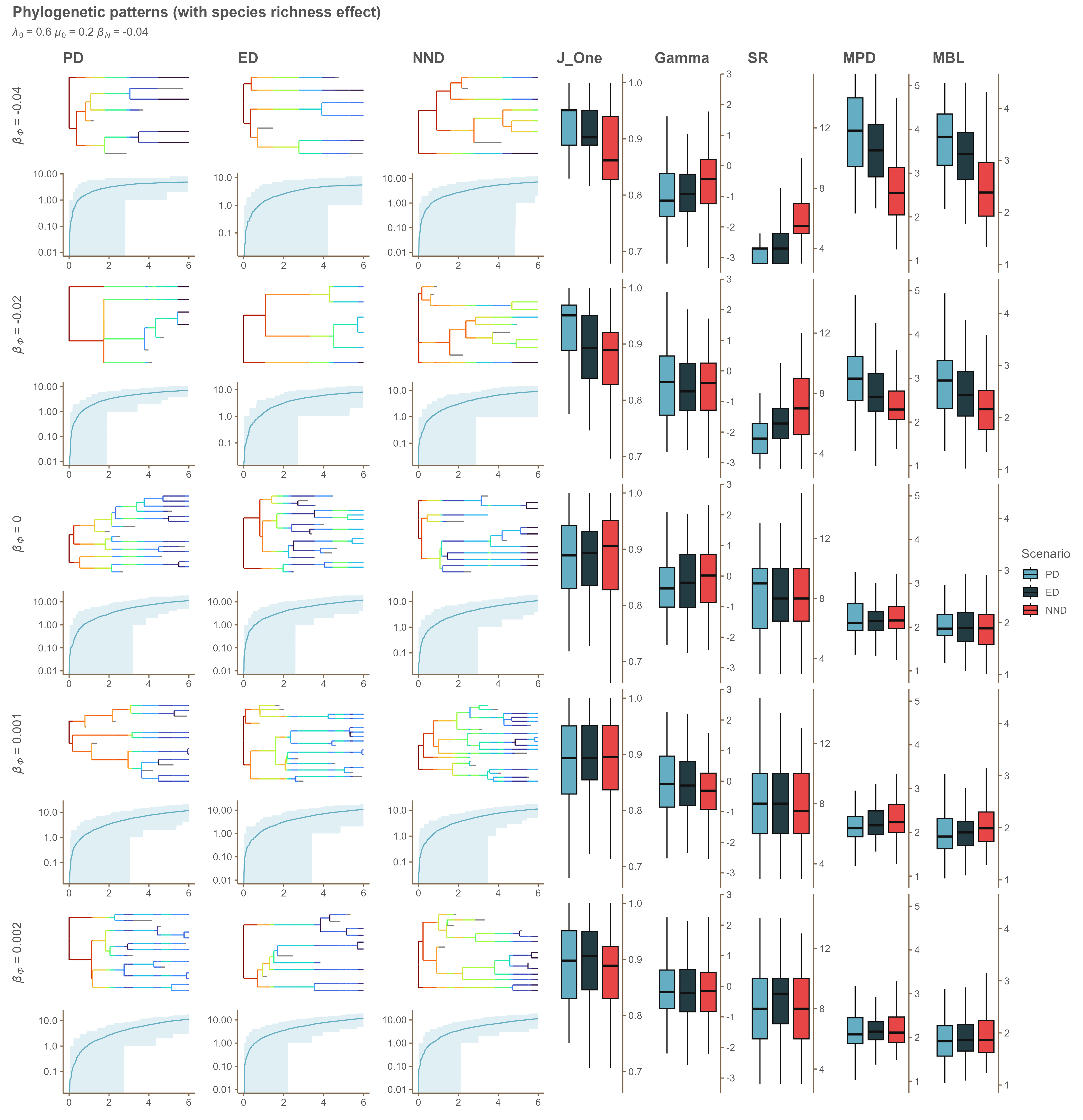

### Appendix D 0.6_0_0.png

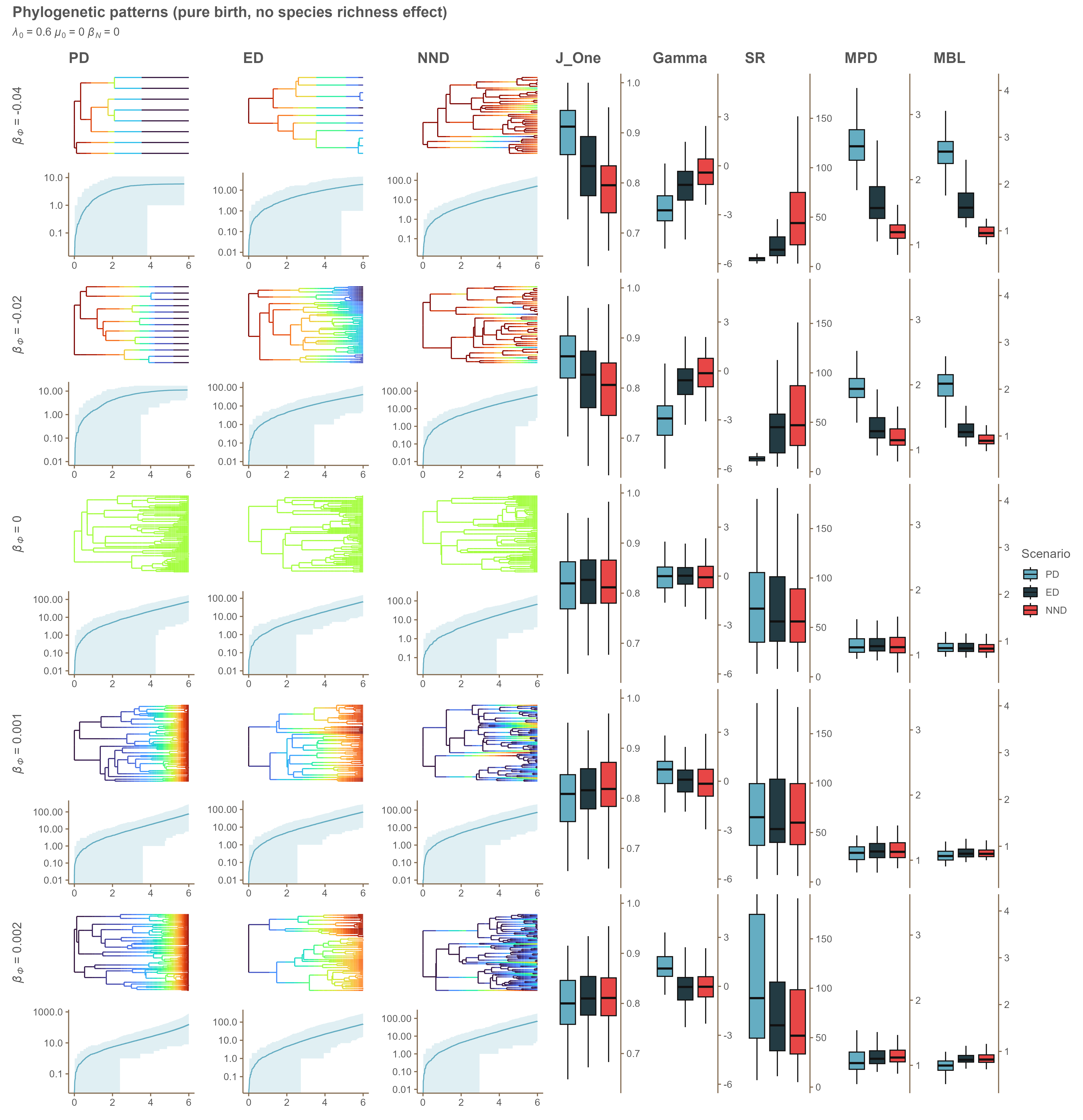

### Appendix D 0.6_0_-0.02.png

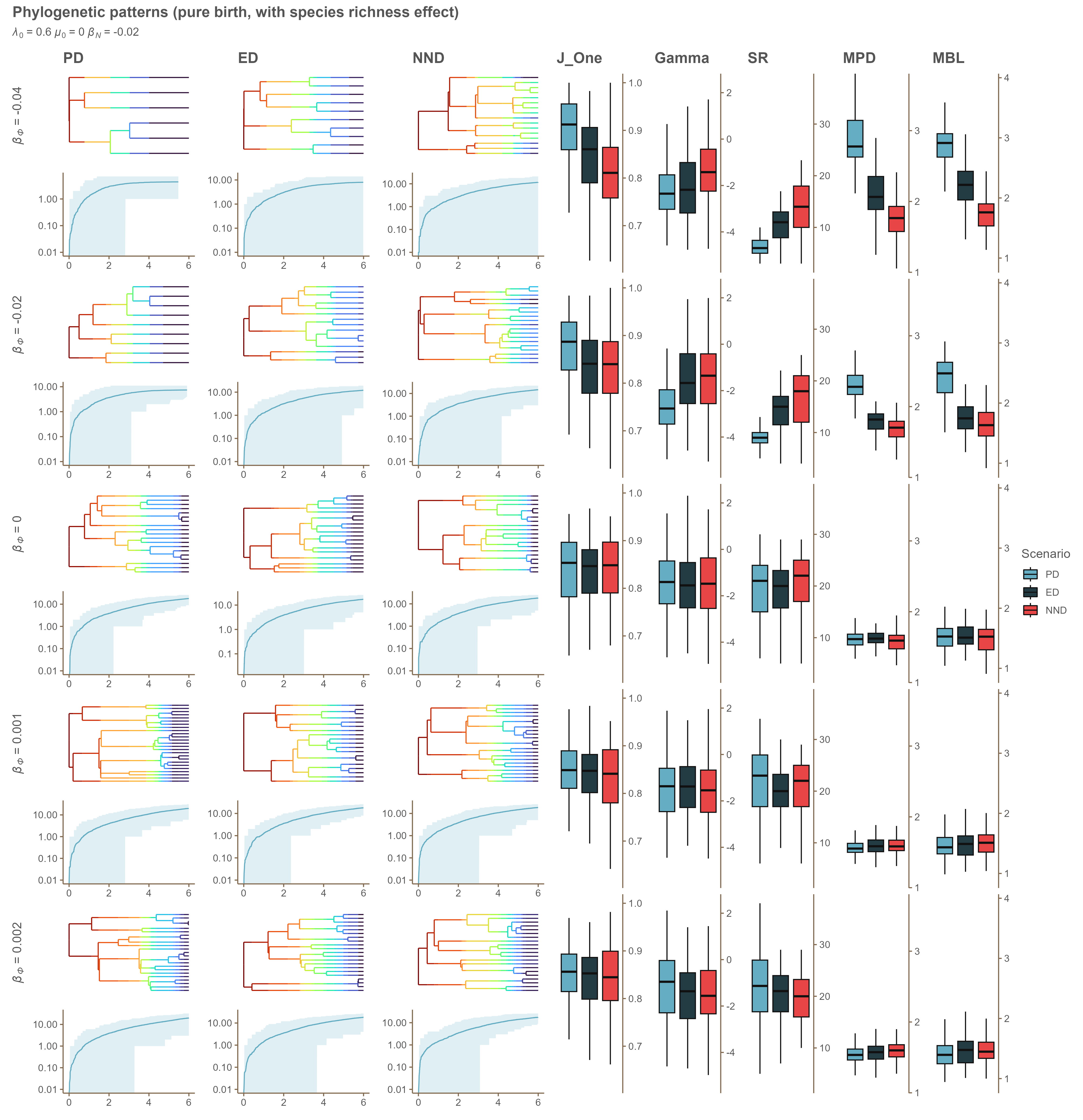

### Appendix D 0.6_0_-0.04.png

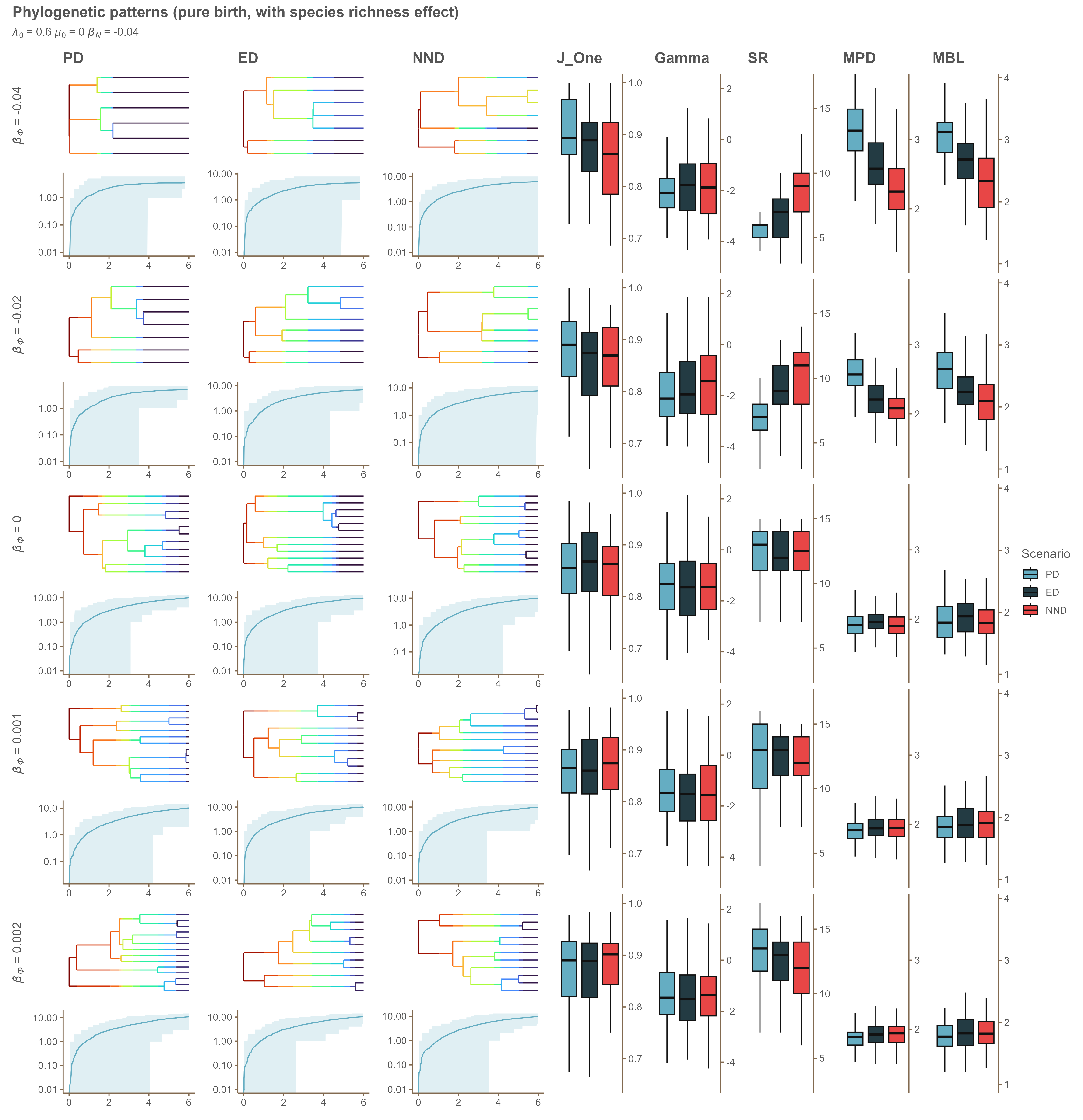
